# Epigenetic silencing of clustered tDNAs in Arabidopsis

**DOI:** 10.1101/2020.02.26.962373

**Authors:** Guillaume Hummel, Alexandre Berr, Stéfanie Graindorge, Valérie Cognat, Elodie Ubrig, David Pflieger, Jean Molinier, Laurence Drouard

## Abstract

Beyond their key role in translation, cytosolic transfer RNAs (tRNAs) are involved in a wide range of other biological processes. Nuclear tRNA genes (tDNAs) are transcribed by the RNA polymerase III (RNAP III) and *cis*-elements, *trans*-factors as well as genomic features are known to influence their expression. In Arabidopsis, besides a predominant population of dispersed tDNAs spread along the 5 chromosomes, some clustered tDNAs have been identified. Here, we demonstrate that these tDNA clusters are transcriptionally silent and that pathways involved in the maintenance of DNA methylation play a predominant role in their repression. Moreover, we show that clustered tDNAs exhibit repressive chromatin features whilst their dispersed counterparts contain permissive euchromatic marks. Our data highlight that the combination of both genomic environment and epigenomic landscape contribute to fine tune the differential expression of dispersed versus clustered tDNAs in Arabidopsis.

## Introduction

In eukaryotes, transcription of nuclear genes is carried by three conserved RNA polymerases (RNAP), named I, II and III. RNAP III transcribes a set of non-coding RNAs, including transfer RNAs (tRNAs), 5S ribosomal RNAs (rRNAs) and other small RNAs, such as the U6 small nuclear RNA (snRNA) or some small nucleolar RNAs (snoRNAs)^1^. As a major function, tRNAs serve as physical link between mRNA and the amino acid sequence of proteins. They are classified according to their anticodon and their cognate amino acid. For each amino acid, tRNAs with distinct anticodons are called tRNA isoacceptors^2^. During evolution, nuclear tRNA genes (tDNAs) have been duplicated several times leading to multi-copy tDNA isoacceptor families whose members are dispersed along chromosomes^2 3 4^. In addition, few nucleotide polymorphisms occur at certain tDNA isoacceptors, which are defined as isodecoders^2^. The number of tDNAs is highly variable among eukaryotic genomes^2 3 4 5 6^ and is not correlated to genome sizes. Nevertheless, it has been reported in *Saccharomyces cerevisiae, Drosophila melanogaster, Caenorhabditis elegans, Chlamydomonas reinhardtii* and in some higher plants that the number of tDNA isoacceptors is correlated with the codon usage and with global translational needs ^7 8 9 10^. Indeed, to enable an effective and rapid protein translation, the abundance of tRNAs cannot be the limiting factor. Therefore, under normal growth conditions, tDNAs were considered as “housekeeping genes” and their transcription by the RNAP III was believed to be constitutive and high^11^. However, several studies on the regulation of RNAP III transcription showed that the rate of tRNA transcription varies in response to various stress conditions^12^ and that identical tDNAs can be differentially expressed in cell-/time-specific manners (for reviews, see^1,13^). Moreover, half of the predicted tDNAs are transcriptionally silent in human^14^ suggesting that tight regulatory processes exist but remain to be deciphered.

Transcription of eukaryotic tDNAs is regulated *via* different ways (for reviews see^12,13^). First, their sequences contain two intrinsic RNAP III internal promoter elements, called A and B boxes. These evolutionary conserved motifs allow the specific recruitment of the general transcription factor (TF) TFIIIC and subsequently TFIIIB for RNAP III pre-initiation complex assembly. While these TFs together with non-core *trans*-factors (*e.g.* Maf1)^12^ play important roles in the regulation of tDNAs biosynthesis, other strategies to modulate tDNAs transcription have emerged during evolution. This includes the presence of other *cis*-elements on tDNAs such as the upstream AT-rich / TATA-like motifs and the downstream short poly-T stretches. This later motif releases RNAP III and allows the recruitment of another *trans*-factor stabilizing the precursor tRNA: the La protein^15^. Moreover, additional *cis*-elements can exist. Indeed, in higher plants, a CAA triplet located between positions −1 and −20 seems essential for transcription initiation^16^. The presence of several consecutive CAA motifs has also been identified^7^. In addition to the important roles of these *cis*-elements and *trans*-factors, a still limited set of data suggest that RNAP III-mediated tDNAs transcription in eukaryotes is submitted to a wider range of regulatory processes such as genomic and epigenomic environments as well as three-dimensional location (for reviews, see^13,17^).

Indeed, nowadays, in eukaryotic organisms, it is well established that normal growth, development and differentiation of distinct cell lineages are governed by transcriptional but also epigenetic mechanisms^18^. In plants, epigenetic processes such as DNA methylation (5-methyl cytosine: 5-mC) and histone post-translational modifications (PTM) are thought to contribute to the regulation of gene expression and the silencing of repeats ^19,20,21^. In mammals, DNA methylation of cytosine residues (5-mC) is restricted to the symmetric CG context, while in plants DNA methylation can occur at CG, CHG and CHH sites (where H=A, T, or C)^21^. In Arabidopsis, 5-mC is deposited by three main types of DNA methyltransferases: METHYLTRANSFERASE 1 (MET1), CHROMOMETHYLASE 2 and 3 (CMT2 and CMT3) and DOMAINS REARRANGED METHYLTRANSFERASE 2 (DRM2). These enzymes are responsible for maintaining methylation in the CG (MET1), CHG (CMT3) and CHH (CMT2 and DRM2) sequence contexts on newly synthetized DNA strands upon replication^21 22 23^. Additionally, cytosines can be methylated *de novo* through the RNA-directed DNA methylation pathway (RdDM) involving two plant-specific RNA polymerases evolutionarily related to RNAP II^24^, RNAP IV and RNAP V. RNAP IV, together with RNA-DEPENDENT RNA POLYMERASE 2 (RDR2), produces double-stranded RNA (dsRNA) precursors that are diced into 24-nt siRNAs by DICER-LIKE 3 (DCL3) and loaded into ARGONAUTE 4 (AGO4)^21 22 23 24 25^. These 24-nt siRNAs are thought to interact with scaffold transcripts produced by RNA POL V, thereby mediating *de novo* DNA methylation of cognate DNA sequences in the three cytosine contexts by DRM2^21,25^. Importantly, the SWI2/SNF2-related chromatin remodeling factor DECREASE IN DNA METHYLATION 1 (DDM1) provides DNA methyltransferases access in cooperation with the RdDM pathway^26^. In addition to DNA methylation, regulation of gene expression can be mediated by chromatin remodeling and histone modifying enzymes. In plants, the computational integration of high-throughput epigenomic data at the whole genome level has led to the identification of different epigenomic profiles named chromatin states^27-30^. Chromatin states are meaningful in several biological functions and impact the activity of gene expression during developmental processes and in response to environmental cues^31^.

In Arabidopsis, while most tDNAs are scattered throughout the nuclear genome, it was previously shown that part of the proline (Pro), serine (Ser) and tyrosine (Tyr) tDNAs are organized in clusters^7^ and, here, we identify a novel cluster formed by cysteine (Cys) tDNAs. In this study, we then demonstrate that while tDNAs dispersed along chromosomes are expressed, clustered tDNAs are transcriptionally silent although most of them possess the canonical RNAPIII *cis*-elements. Using *in silico* analyses we unveil that these clustered tDNAs contain elevated CG methylation levels compared to their dispersed counterparts. Molecular approaches allowed identifying that particular mutant plants defective for the maintenance of DNA methylation pathways exhibit a release of the expression of clustered tDNAs. Concomitantly to DNA methylation, we reveal that clustered tDNAs display heterochromatic features whereas their dispersed equivalents show euchromatic characteristics. Collectively, our findings reveal that both genomic environment and epigenetic landscape act to fine tune the differential expression of dispersed and clustered tDNAs in Arabidopsis.

## Results

### Extra copies of tDNAs are organized into clusters in Arabidopsis

In various organisms, a positive correlation exists between the number of tDNA copies specific for each amino acid and the corresponding frequency of occurrence of the same amino acid and codon(s) ^8 10^. We have previously shown^7^ that this correlation holds true in Arabidopsis except for the Pro, Ser and Tyr tDNA families. These families are in excess and lie clearly outside of the regression line between the number of genes for each tDNA families and the frequency of occurrence of each amino acid (Fig. 1a). According to our previously published analyses ^6 7^, this excess of copies is organized in several clusters with adjacent tDNAs in the same orientation (Figs. 1c; Supplementary Table 1). Two clusters of Pro tDNAs (*i.e.* one with 24 copies and one with 6 copies) are located on Arabidopsis Chromosome 1 (Chr1), nearby the centromeric/pericentromeric regions, and one cluster of 9 copies on Chr2 long arm (Fig. 1b). Clustered Ser and Tyr tDNAs are found on Chr1 arranged in interspaced Ser-Tyr-Tyr units tandemly repeated 27 times nearby the pericentromeric region (Fig. 1b). Interestingly, we further identified an additional cluster on Chr5 formed by 4 Cys tDNAs oriented in the same direction (Figs. 1b and 1c; Supplementary Table 1). Compared to the other clusters, the low number of gene copies in the Cys tDNAs cluster does not drastically affect the positioning of the Cys point on the regression line (Fig. 1a). Lastly, while Cys and Ser-Tyr-Tyr repeats are regularly interspaced, clustered Pro tDNAs are not (Fig. 1c).

**Fig. 1.**
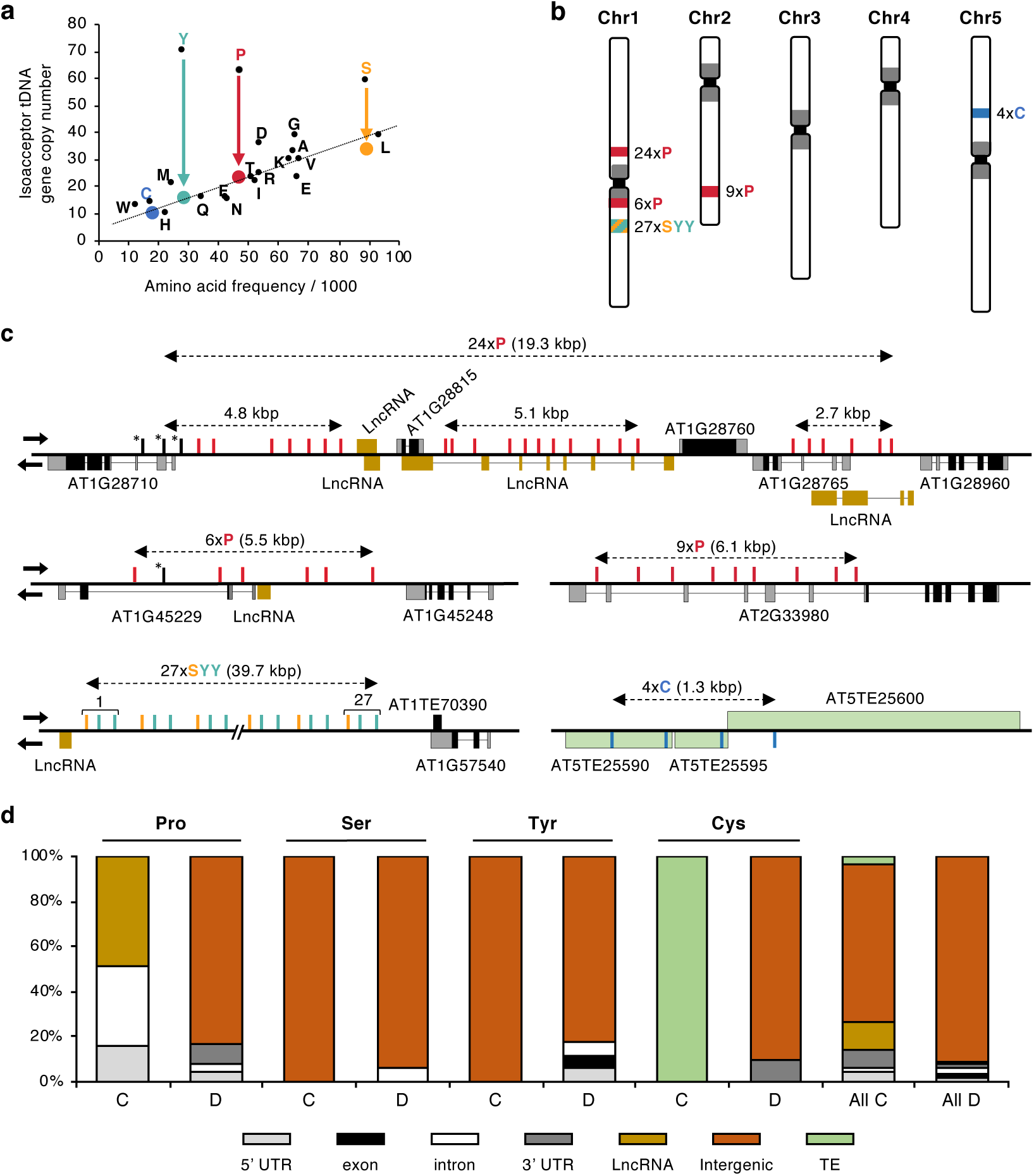
Characterization of tDNAs clusters and their genomic context in *Arabidopsis*. **a**, Correlation between the number of tDNAs specific for each amino acid and the frequency of occurrence of the same amino acid. For four amino acids, Ser, Tyr, Pro and Cys, the numbers corresponding to tDNAs found in clusters were subtracted and the colored dots represent the new correlation (colored arrows indicate the corresponding shifts). The one-letter code for amino acid is used and the color code is green for Y, red for P, yellow for S and blue for C. In this figure adapted from previously published work^7^, the presence of Cys tDNAs in cluster has been added following re-analysis of the data. **b**, Schematic representation of the tDNAs clusters loci present in chromosomes 1, 2 and 5 of *Arabidopsis*. The numbers of repeated tDNA copies are indicated. Pericentromeric regions are in gray. **c**, Scheme of the clustered tDNAs genomic regions. tDNAs are indicated by colored vertical bars (green for Y, red for P, yellow for S and blue for C). Accession numbers of protein coding genes and transposable elements (TE) is given. Black and gray boxes represent exons in CDS and UTRs respectively, lines represent introns. Brown boxes depict putative long non coding RNAs (LncRNAs). Arrows indicate transcription orientation. **d**, Distribution of the location of tDNAs on the Arabidopsis genome. Clustered (c) and dispersed (d) tDNAs specific for Pro, Ser, Tyr, Cys and Ala have been manually analyzed. Data were retrieved from the TAIR10.1 genome browser.

Unlike dispersed Pro tDNAs that are primarily intergenic, their clustered counterparts are predominantly located in long non-coding RNAs (LncRNAs) or in protein coding gene (PCG), within introns or 5’-untranslated regions (5’-UTR), and always in opposite orientation of the transcriptional unit (Figs. 1c and d). The Ser-Tyr-Tyr cluster location is exclusively restricted to a large intergenic region of around 40 kbp, resembling the genetic environment predominant for dispersed tDNAs. Finally, the Cys cluster is positioned at three transposable elements (TEs) belonging to the Helitron superfamily (Figs. 1c and d). Taken together, it emerges that Pro and Cys tDNA clusters stand in distinct genetic environments compared to dispersed tDNAs, while the Ser-Tyr-Tyr cluster does not.

### RNAP III *cis-*acting elements are well conserved among clustered and dispersed tDNAs

Transcription efficiency of plant tDNAs by RNAP III is based notably on the presence of several upstream and downstream *cis*-acting elements^16,32^. To further explore the genetic architecture of Arabidopsis clustered tDNAs, their promoter and terminator regions were analyzed in details and compared to those of dispersed tDNAs. Corresponding sequences were retrieved from the PlantRNA database ^6^. Whether or not from clustered tDNAs, promoters of Pro, Tyr and Cys tDNAs are A/T-rich and present a CAA triplets in the −7 to −3 region (Supplementary Fig. 1a) known to be important for efficient RNAP III transcription^16,32^. Promoters of clustered Ser tDNAs are also enriched in A and T residues but do not display a CAA motif (Supplementary Fig. 1a and 1b). Downstream, both clustered and dispersed tDNAs contain a short poly(T) tail (Supplementary Fig. 1a). These stretches of T residues are shorter (but of sufficient length, *i.e.* at least 4 Ts^14^) for clustered tDNAs compared to dispersed ones (Supplementary Fig. 1a). Altogether, these analyses demonstrate that almost all clustered tDNAs exhibit the required core components to be potentially transcribed by RNAP III.

### Clustered tDNAs are transcriptionally silent

Given that clustered tDNAs represent extra copies of tRNA genes and that most of them exhibit all features for RNAP III transcription, we were interested in assessing the level of expressed tRNAs originating from these clustered tDNAs in the whole tRNA population. In Arabidopsis, as in other eukaryotes, a high level of conservation between tRNA sequences among each tRNA isoacceptor family exists^6^ preventing a straightforward differential expression analysis. To overcome this impediment, the clustered and dispersed tDNA sequences were analyzed into details (Supplementary Fig. 2) to identify potential polymorphisms that would allow discriminating specifically their expression level by northern blotting. Interestingly, we found nucleotide polymorphisms between the most abundant clustered and dispersed tDNA Tyr and Pro copies (Supplementary Figs. 2 and 3), allowing the design of specific oligonucleotide probes (Fig. 2).

**Fig. 2.**
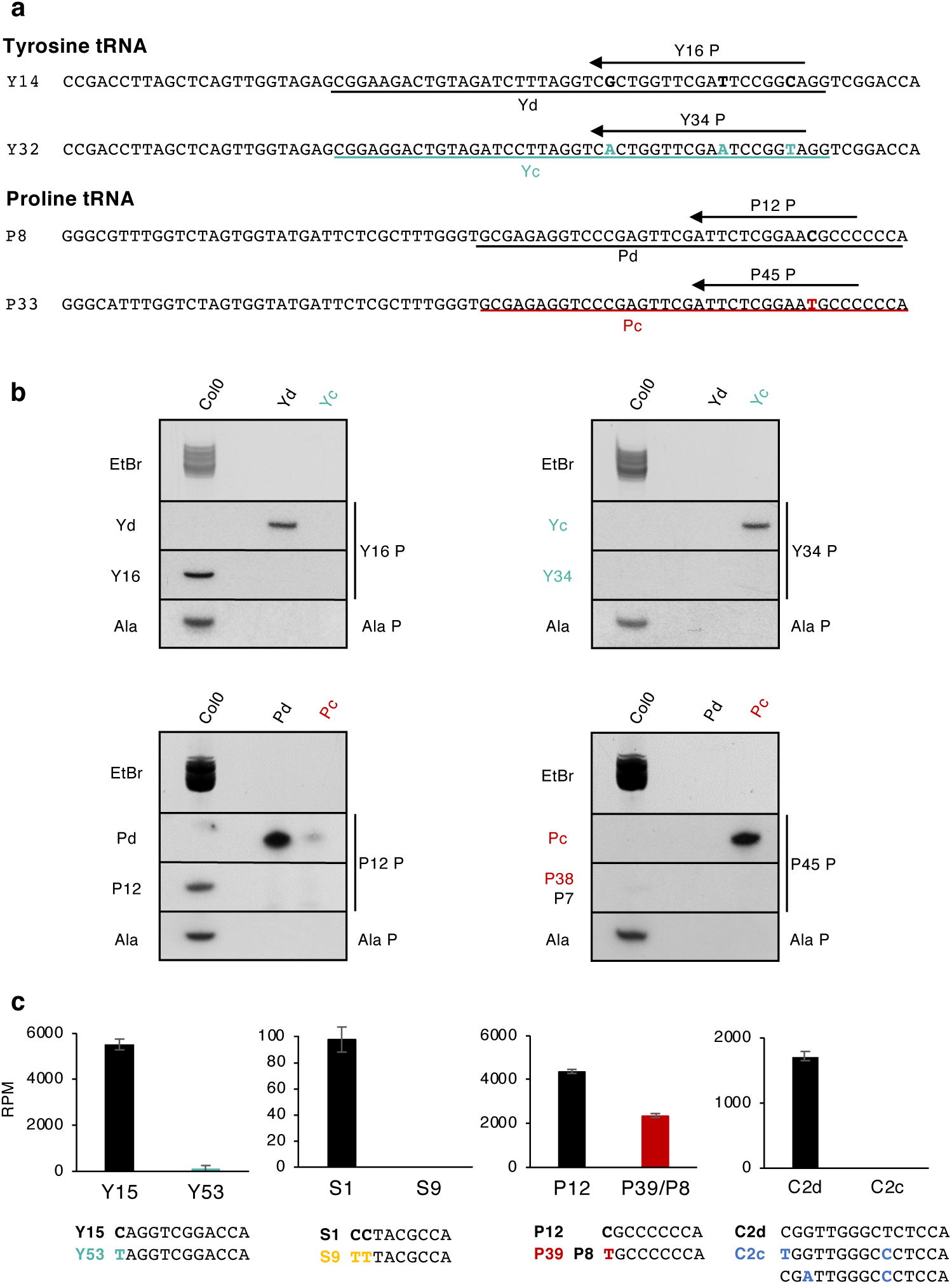
Expression analysis of *Arabidopsis* tDNA clusters. **a**, Predominant sequences of dispersed (Y14 and P8) and clustered (Y32 and P33) Tyr and Pro tDNAs. Nucleotide differences are in bold and colored in green and red for Tyr and Pro clustered tDNAs respectively. Underlined sequences (Yd, Yc, Pd and Pc) were used as controls of specificity and sequences indicated by arrows (Y16 P, Y34 P, P12 P and P45 P) were used as oligonucleotide probes during northern blot experiments. Note that the P45 P probe designed for the clustered Pro tDNAs (P38) is also complementary to 7 dispersed Pro tDNAs (P7), (Supplementary Fig. 2). **b**, northern blot detection of dispersed (upper panel left) and clustered (upper panel right) Tyr plus dispersed (lower panel left) and clustered (lower panel right) Pro tRNAs in Col0 WT shoots. Nomenclatures of the left and right sides refer to controls and probes as depicted in **a**. Staining with ethidium bromide (EtBr) and hybridization with a probe (Ala P) specific to Arabidopsis cytosolic Alanine tRNA were used as loading control. **c**, Histograms showing the abundance of tRFs specific to dispersed Tyrosine (Y15), Serine (Y1), Proline (P12) and Cysteine (C2d) tRNAs and clustered Tyrosine (Y53), Serine (S9), and Cysteine (C2c) tRNAs. The tRF sequence “TGCCCCCCA” is specific to 40 clustered Pro tRNAs (P40) and 8 dispersed Pro tRNAs (P8). tRFs sequences are indicated below each graph with nucleotide differences in bold and colored. tRFs with a size between 19- and 26-nt were analyzed. Error bars represent standard errors of mean of two independent libraries.

Among the Tyr tDNAs, 3 nucleotide polymorphisms (G49A, T59A and C65T) allows discriminating the sequences of the dispersed Tyr tDNAs from those of the predominant clustered Tyr tDNA copies. Thereby, the probe Y16 P would be specific to 16 dispersed Tyr tDNAs, while the probe Y34 P would be specific to 34 Tyr tDNAs out of the 54 clustered Tyr tDNAs (Fig. 2a and Supplementary Fig. 2). Northern blot analyses using two synthetic oligonucleotides (Yd and Yc), representative of these dispersed and clustered Tyr tDNAs, respectively (Fig. 2a), confirm that Y34 P and Y16 P probes allow discriminating specifically between Tyr tRNA originating from clustered *versus* dispersed tDNAs (Fig. 2b). Similarly, a single-nucleotide polymorphism (C67T) is sufficient to distinguish by northern blot a majority of clustered Pro tDNAs (*i.e.* 38 out of 39) from most dispersed ones (Fig. 2b and Supplementary Fig. 2). Indeed, the P45 P probe looks specific to 38 clustered Pro tDNAs, but also to 7 out of 24 dispersed ones since they share the same sequence. Conversely, the P12 P probe can exclusively recognize 12 dispersed Pro tDNAs (Supplementary Fig. 2a and Fig. 2a). The specificity of the two probes, P45 P and P12 P was confirmed by northern blot experiments using as controls two oligonucleotides, Pd and Pc, representative of the major dispersed and clustered Pro tDNAs respectively (Fig. 2b). Unfortunately, the small numbers of gene copies with sequence polymorphisms prevented the design of specific probes for the differential analysis of Ser and Cys tDNAs by northern blot (Supplementary Fig. 2b).

While Tyr tRNA isodecoders expressed from dispersed copies are detectable using the Y16 P probe, those originating from clustered copies (Y34 P probe) are below the limits of detection (Fig. 2b). Pro tDNAs expression is also detected for dispersed copies using the P12 P probe (Fig. 2b). Although a weak signal is observed using the P45 P probe (Fig. 2b), we believe that it most likely reflects the ability of this probe to recognize not only clustered Pro tDNAs, but also few copies of dispersed ones. Taken together, these molecular analyses suggest that clustered tDNAs are likely silenced in Arabidopsis.

To determine whether this lack of detection relies on a transcriptional or on a post-transcriptional regulatory process, we used publicly available small RNA libraries^33^ to undertake a degradome analysis of tRNA-derived fragments (tRFs) bearing polymorphic ribonucleotides (Fig. 2c, Supplementary Fig. 2). For this analysis, we assumed that tRF levels reflect tRNA abundance, although we cannot exclude that half-life between tRF species may differ. tRFs originating from tRNAs expressed from clustered Tyr tDNAs (Y53) are barely detectable, whereas those from dispersed Tyr tDNAs (Y15) are abundant (Fig. 2c). Similarly, while tRFs derived from a single dispersed Ser tRNA (S1) are present in the two analyzed small RNA libraries, tRFs arising from the 9 clustered Ser tDNAs (S9) are absent (Fig. 2c). For Pro tRNAs, the level of tRFs derived from 12 dispersed tRNA genes (P12) is higher than that derived from 40 clustered (P40) plus 8 dispersed tDNAs (P8), (Fig. 2c). Finally, tRFs arising from 2 clustered Cys tRNAs are not detected, whereas those arising from 2 dispersed Cys tDNAs exist (Fig. 2c).

Collectively, although we cannot fully exclude the existence of post-transcriptional regulatory processes, our analyses strongly suggest that the expression of clustered Ser, Tyr, Pro and Cys tDNAs is predominantly transcriptionally repressed in Arabidopsis.

### Clustered tDNA repeats display elevated DNA methylation levels

We assumed that the lack of expression characterizing clustered tDNAs may be due to the presence of repressive epigenetic marks, while dispersed ones may contain permissive marks to be expressed. Among repressive epigenetic marks, DNA methylation is well known to play a key role in plant transcriptional gene silencing^34^. Therefore, the DNA methylation landscape at both dispersed and clustered Ser, Tyr, Pro and Cys tDNAs in Arabidopsis plants was analyzed. Using publicly available data^35^, 5-mC levels in CG, CHG and CHH contexts were calculated for each single clustered tDNA locus within a genomic window starting 50 bp upstream (promoter) and ending 25 bp downstream (terminator) of each tDNA (Fig. 3a). Importantly, the cytosine content is homogeneous between all studied tDNAs, thus allowing comparative studies (Fig. 3b and Supplementary Fig. 4a). As shown in Fig. 3c, all clustered tDNAs exhibit higher methylation levels in the CG context compared to their dispersed counterparts. CHG and CHH methylation levels are slightly higher for clustered Ser, Tyr and Cys tDNAs compared to their dispersed equivalents, whereas this is not the case for clustered Pro tDNAs (Fig. 3c). Additionally, while analyzing the methylome at single-nucleotide-resolution we found that DNA methylation occurs mainly in the body of clustered tRNA genes (Fig. 3d). Cytosines at promoter and terminator regions are also methylated but to a low extent (Fig. 3d). Importantly, the dispersed Ala tDNAs, used as control in this study, display low DNA methylation levels in all contexts (Supplementary Figs. 4b and 4c).

**Fig. 3.**
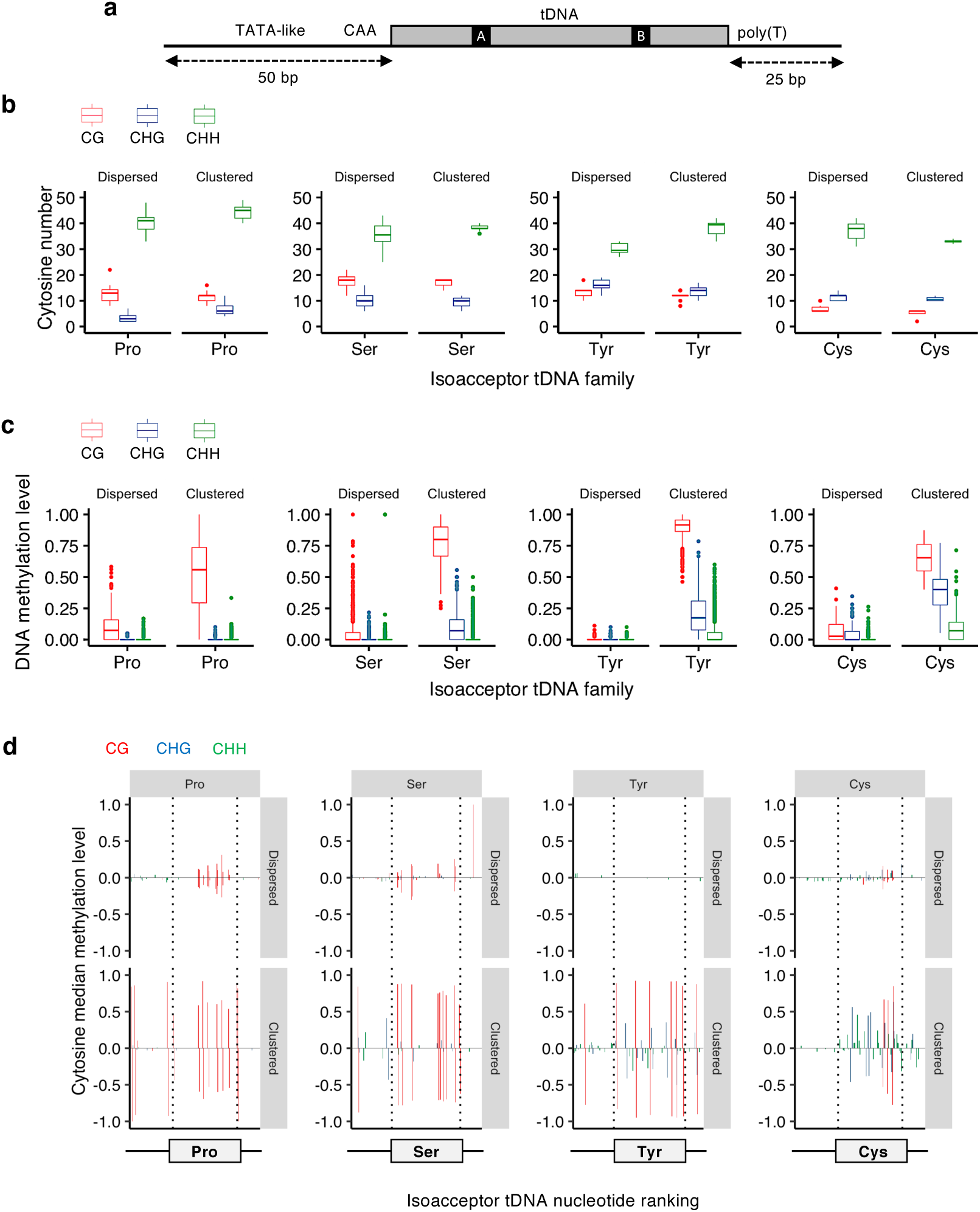
Methylation landscape at dispersed and clustered tDNA loci. **a**, Schematic representation of a tDNA locus with the transcriptional internal control regions called A and B boxes, the upstream TATA-like elements and CAA motif and the downstream poly-T tract^4^. For each tDNA, the region spanning from 50 bp upstream and 25 bp downstream was examined for its DNA methylation mean level in CG (red), CHG (green) and CHH (blue) contexts. **b**, Boxplots representing DNA cytosine counts present on dispersed and clustered Pro, Ser, Tyr and Cys tDNAs. **c**, Boxplots representing DNA cytosine methylation levels on dispersed and clustered Pro, Ser, Tyr and Cys tDNAs. **d**, DNA cytosine methylation median levels at single nucleotide resolution on dispersed and clustered Pro, Ser, Tyr and Cys tDNAs. To note, only predominant polymorphic sequences of identical length have been used for these representations. Positive and negative values are referring to positive and negative DNA strands, respectively. The regions corresponding to tDNA sequences are delimited by dotted lines. The statistical analysis using pairwise Wilcoxson tests are provided in Supplementary Data 1.

To test whether these DNA methylation profiles are modulated during plant development, we analyzed the DNA methylation landscapes of dispersed and clustered tDNAs in different tissues and organs using publicly available methylomes ^22 36 37 38 39 40 41 42^, (Supplementary Fig. 5, Supplementary Table 2). The CG context shows the higher methylation level in clustered tDNAs compared to dispersed tDNAs in all tested samples. Reproductive tissues contain higher CHG methylation levels for Ser, Tyr and Cys tDNA clusters compared to somatic tissues (Supplementary Fig. 5). Clustered Ser and Tyr tDNAs exhibit increased CHH methylation levels in the central cell compared to the other tissues (Supplementary Fig. 5). In contrast, the control dispersed Ala tDNAs do not display any strong variations. Furthermore, using northern blot analyses, we found that, the tRNA levels of clustered Ser, Tyr and Pro tDNAs were below the detection limit in all tested organs, while tRNAs originating from dispersed tDNAs were detectable (Supplementary Fig. 6).

Collectively, these data reflect that a negative correlation likely exists between the CG methylation levels and the expression of clustered tDNAs.

### Effectors of maintenance of DNA methylation regulate methylation landscape at clustered tDNAs

The methylome analyses of tDNA families reveal that clustered tDNAs contain high levels of DNA methylation compared to dispersed ones. To determine the involvement of particular DNA methylation pathways in these profiles, we re-analyzed the DNA methylation levels of clustered and dispersed tDNAs in mutant plants defective for the expression of the main Arabidopsis DNA methylation processes^35^ (Supplementary Table 2 and Fig. 4). In this way, as expected, we show that for all clustered tDNAs, the CG methylation level relies on the MET1 DNA methyltransferase (Fig. 4). In addition, the CG methylation pattern is strongly affected in the *ddm1* mutant plants for the Ser, Tyr and Cys clusters but not for the Pro one (Fig. 4). Enhanced CHG and CHH DNA methylation levels are observed in *ddm1* and *met1* for the Ser and Tyr cluster, respectively (Fig. 4) in agreement with the ectopic gain of CHH methylation already reported in both mutant plants^35^. Our analyses further show that for the Ser, Tyr and Cys clusters, CHG methylation relies on CMT3 and KYP. For all clustered tDNAs, CHH methylation levels are poorly affected in both *cmt2* and *drm2* mutant plants (Fig. 4), suggesting that both CMT2 and RdDM may act synergistically. Importantly, our data also show that the Pro tDNA clusters are under the control of the histone H3K9 demethylase INCREASE IN BONSAI METHYLATION 1 (IBM1), known to indirectly modulate CHG methylation levels^43^. Indeed, *ibm1* mutant plants exhibit enhanced CHG methylation level at clustered Pro tDNAs (Fig. 4). Interestingly, among the Pro clusters, one of them is located within the coding sequence of AT2G33980 (Fig. 1c), reported as an IBM1 target^43^. Thus, our analyses suggest that the dynamics of histone PTM (*i.e.* H3K9me2) may also influence DNA methylation levels at this particular locus (Fig. 4). Together, our data show that DDM1, MET1, CMT2, CMT3, KYP, IBM1 and DRM2 maintain high level of DNA methylation in clustered tDNAs.

**Fig. 4.**
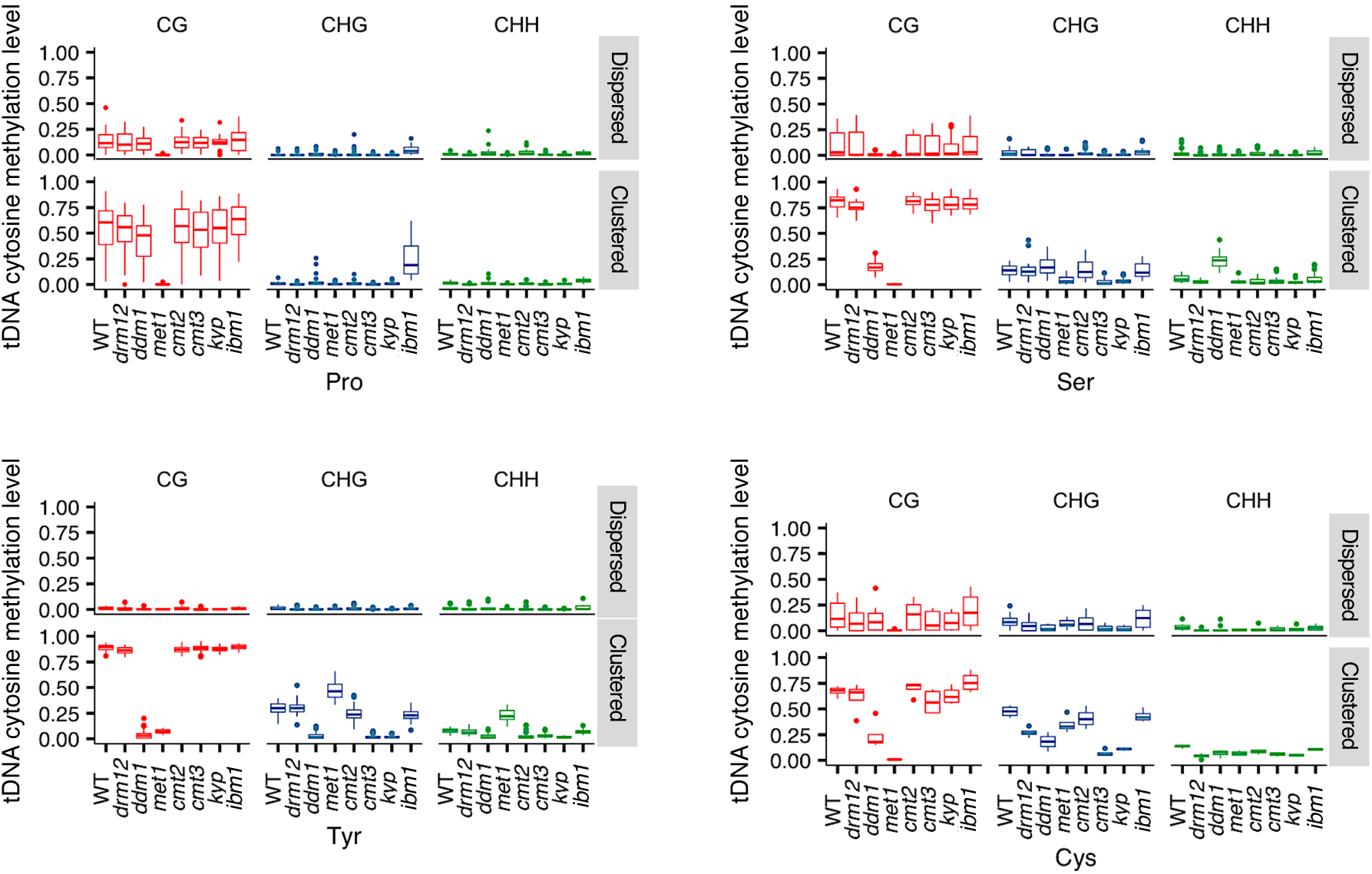
DNA methylation landscape at clustered tDNAs in DNA methylation mutant plants. Box plots representing DNA methylation levels in various mutant lines for dispersed and clustered Pro, Ser, Tyr and Cys tDNAs. CG (red), CHG (green) and CHH (blue) contexts are indicated. The statistical analysis using pairwise Wilcoxson tests are provided in Supplementary Data 2.

In order to determine whether DNA methylation restricts clustered tDNAs expression, we experimentally analyzed the tRNA steady state levels by northern blots using different strategies. In a first approach, we challenged the effect of a global loss of cytosine methylation on clustered Tyr and Pro tDNAs expression by assaying the effect of the cytidine analog 5-azacytidine (5-azaC)^44^. The 5-azaC treatment significantly released clustered Tyr tDNAs expression whilst no effect was detected for the Pro ones (Fig. 5a). This pharmacological approach highlights that DNA methylation represses clustered Tyr tDNAs expression. In a second approach, we used mutant plants deficient for the expression of RdDM factors and of the maintenance of DNA methylation pathways. In RdDM mutant plants, no release of clustered Tyr and Pro tDNAs expression could be detected (Supplementary Fig. 6), suggesting that this pathway does not play a predominant role in their silencing. Moreover, both *ddm1* and *met1* mutant plants display re-expression of clustered Tyr tDNAs (Fig. 5b), in agreement with the reduced CG methylation levels observed in these mutants (Fig. 4). Surprisingly, clustered Pro tDNAs do not display release of expression in *ddm1* and *met1* mutant plants, suggesting that additional factors may repress their expression (Fig. 5b). Importantly, no transcriptional reactivation is observed for clustered Tyr and Pro tDNAs in *cmt2, cmt3* and *kyp* plants (Fig. 5c), demonstrating that both CHG and CHH methylation are not the predominant contexts repressing clustered tDNAs expression.

**Fig. 5.**
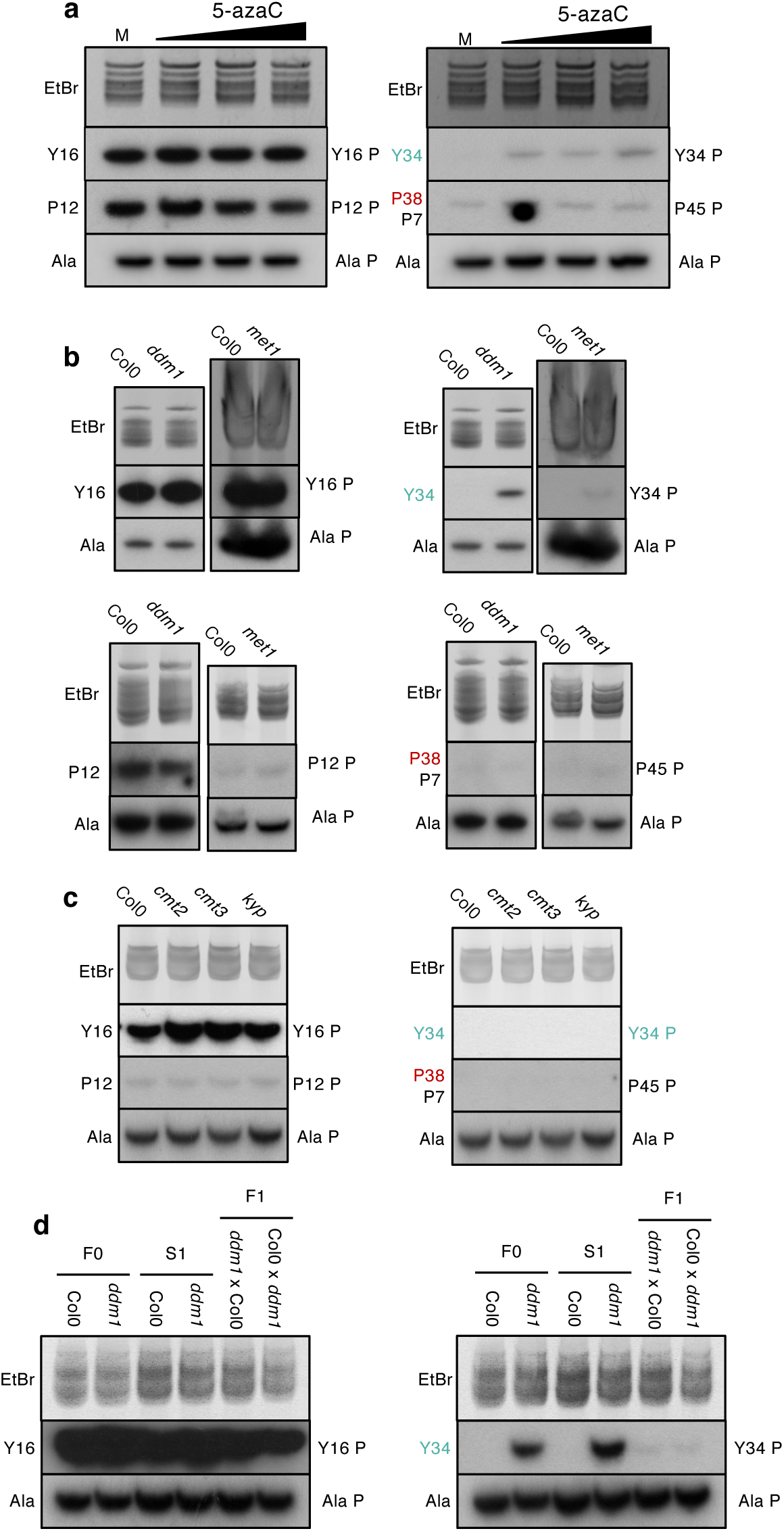
Effect of DNA hypomethylation on tDNAs cluster expression. **a**, northern blots analysis of total tRNA from untreated (M) or 5-azaC (0.25, 0.5 and 1 mM) treated WT Arabidopsis plants. **b** and **c**, northern blots analysis of total tRNA extracted from leaves of the indicated mutant genotypes. Experimental conditions and probes are similar to those described on Fig. 2b. **d**, northern blot analysis in self (S1) and reciprocal (F1) hybrids between Col0 and *ddm1* mutant plants. Names of probes are as in Fig. 2.

In addition to all tRNAs, RNAP III also transcribes the 5S rRNA^1^. In Arabidopsis, the helicase MORPHEUS MOLECULE 1 (MOM1) is known to mediate silencing of 5S rDNA repeats in a DNA methylation independent manner^45 46 47^. We therefore explored the possible impact of MOM1 on the silencing of clustered tRNA genes by northern analysis using Y34 P and Y16 P for Tyr and P45 P and P12 P for Pro (Fig. 2a). Confirming that DNA methylation likely predominantly regulates the silencing of clustered tDNAs, Tyr and Pro tDNAs were not re-activated in the *mom1* mutant (Supplementary Fig. 6c).

Given that DNA methylation must be efficiently maintained through mitosis and meiosis, we tested whether inheritance/maintenance of DNA methylation would repress the expression of clustered tDNAs through a directional parental effect. For that purpose, we analyzed clustered Tyr tDNAs expression in the progeny of reciprocal crosses between WT and *ddm1* mutant plants. As above reported, the expression of clustered Tyr tDNAs is released in the *ddm1* self-crossing plants (S1; Fig. 5d). In both F1 plants *ddm1*/Col0 and Col0/*ddm1*, the expression of clustered Tyr tDNAs could still be detected, albeit weaker than in *ddm1* parent plants (Fig. 5d). These data reflect that maintenance of DNA methylation is important to efficiently repress the expression of clustered tDNAs and demonstrate its bi-parental inheritance.

As a whole, the combination of *in silico* and molecular approaches allowed determining that DNA methylation plays a predominant role in the silencing of clustered tDNA repeats and that efficient maintenance of CG cytosine methylation is crucial to repress their expression.

### Clustered tDNAs overlap with heterochromatin states

Besides DNA methylation, histone modifications and histone variants are integral part of the molecular features that characterize the chromatin landscape in eukaryotic cells. As such, certain chromatin signatures of genomic elements establish particular gene expression patterns. Here, we examined the relationship between clustered tDNAs and each of the nine chromatin states (CS) previously described in Arabidopsis^29^. Dispersed Pro, Ser and Tyr tDNAs, as well as the Ala control, are similarly enriched in CS2 (Fig. 6a), a state characterized by the coexistence of active/repressive marks (*e.g.* H3K4me3 and H3K27me3) in proximal promoter regions and considered as the second most active state according to the first principal component analysis^29^. This enrichment in CS2 is significantly higher than the average distribution of CS observed over the entire Arabidopsis genome, with all *p* value < 0.005 (Supplementary Table 3). For the dispersed Cys tDNAs, the CS distribution is roughly similar to the average one, with CS1, CS2, CS4 and CS6 being the most abundant states. While dispersed tDNAs are mainly enriched in euchromatin states, Ser, Tyr and Cys clustered tDNAs are enriched in the heterochromatin state 8 (CS8), a repressive state found within intergenic regions and TE with high levels of the histone variant H3.1, H3K9me2, H3K27me1 and CG methylation^29^. Interestingly, while CS8 characterizes the chromatin of Ser, Tyr and Cys tDNA clusters, CS7 is predominantly present along the chromatin of the three Pro tDNA clusters (Fig. 6a). This state appears almost exclusively related to intragenic regions, as it colocalizes with coding sequences and introns. Moreover, CS7 strongly associates with transcription units longer than average^29^. The dichotomy in chromatin states between the CS8 of Ser, Tyr and Cys tDNA clusters and the CS7 of Pro tDNA clusters is further reinforced through exploiting RNAP II ChIP-seq data^48^. Indeed, tDNA clusters colocalizing with the intragenic CS7 are enriched in RNAP II, while clusters colocalizing with the intergenic CS8 are not (Supplementary Fig. 7).

**Fig. 6.**
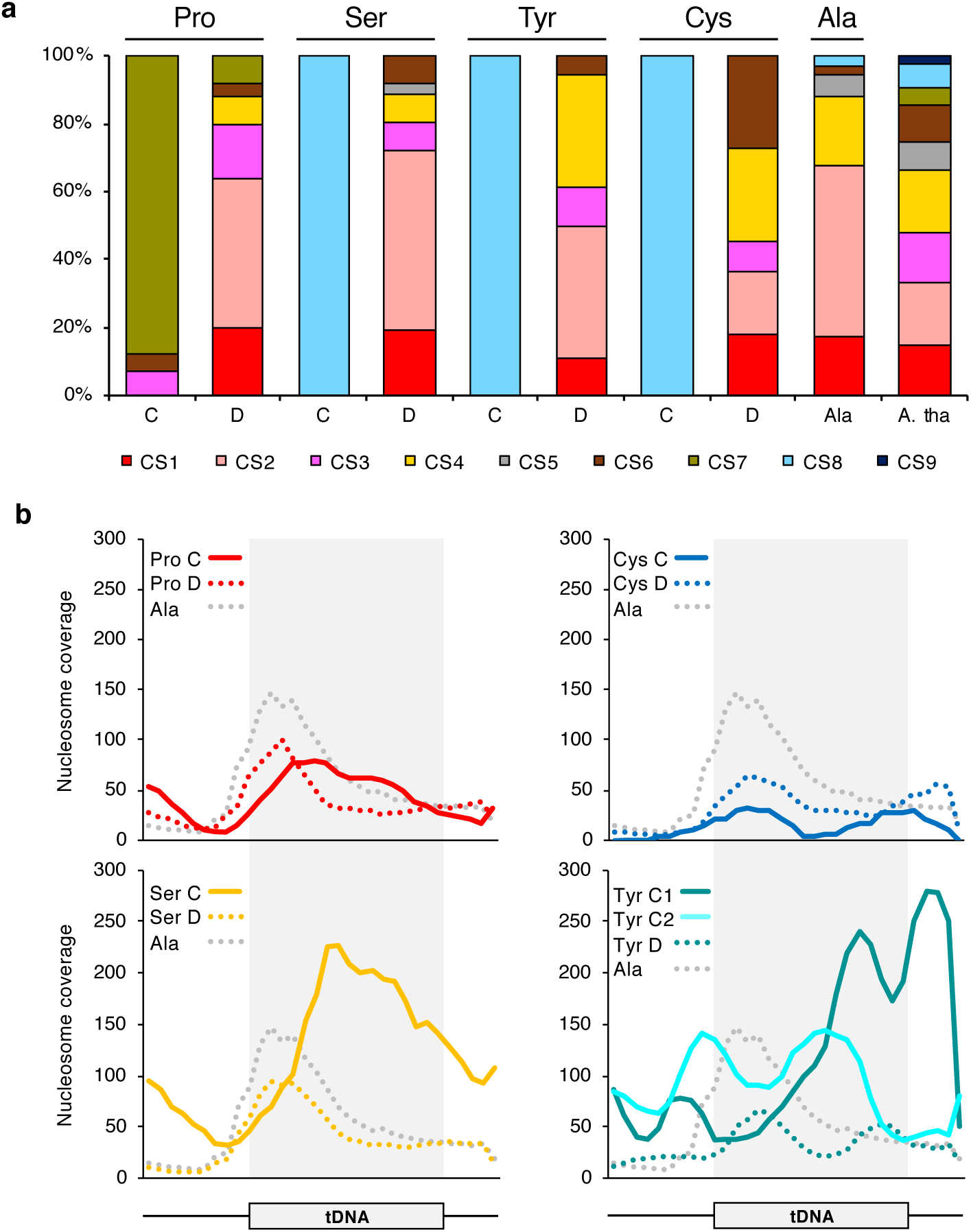
Chromatin states (CS) of dispersed (D) and clustered (C) tDNAs loci. **a**, Histogram showing the proportion of each of the 9 chromatin states (CS)^29^ for the dispersed (D) or clustered (C) tDNAs. Last column (A. tha) gives the proportion of each CS for the entire Arabidopsis genome. **b**, Nucleosome occupancy at Ser, Tyr, Pro, Ala, dispersed (D) and clustered (C) tDNA loci. The two Tyr tDNAs in the Ser-Tyr-Tyr tandemly repeated units are numbered 1 and 2, respectively.

Nucleosomes occupancy is another key factor impacting DNA accessibility in the nucleus. Methods combining digestion of unbound double-stranded DNA using enzymes such as micrococcal nuclease (MNase) and high throughput sequencing of the remaining protected nucleosome-bound DNA allow the generation of genome-wide maps of nucleosome positions. Furthermore, while being enriched at heterochromatin, nucleosome positioning was also reported to influence DNA methylation patterning throughout the genome. Indeed, DNA methyltransferases were found to preferentially target nucleosome-bound DNA^49^. We therefore decided to explore the nucleosome occupancy (NO) at clustered tDNAs in comparison to non-clustered ones using public MNase-seq data^50^. In general, NO was higher in the body of tRNA genes suggesting access for TF recruitment in the upstream genic region (Fig. 6b, Supplementary Fig. 8). Supporting our chromatin data, the NO was lower at Pro tDNA clusters (*i.e.* enriched in the euchromatic CS7) compared to Ser and Tyr tDNA clusters (*i.e.* enriched in the heterochromatic CS8). Considering clustered Ser and Tyr as interspaced Ser-Tyr-Tyr units tandemly repeated, we found that while nucleosomes often localized at the center of the body of the Ser and of the second Tyr tDNA repeats (Tyr C2), they are located toward the 3’ end of the first Tyr tDNAs (Tyr C1; Fig. 6b). Finally, despite the primarily heterochromatic state of Cys tDNA clusters, NOs were similar between clustered and dispersed Cys tDNAs, suggesting a mechanism of heterochromatic silencing different from that of clustered Ser and Tyr repeats.

Together, our analysis demonstrates that clustered tDNAs display particular chromatin features compared to their dispersed equivalents. Moreover, clustered tDNAs can be split into three categories based on their chromatin organization, (*i*) Pro with a predominant euchromatic state mainly associated with long genes and intronic regions, (*ii*) Ser and Tyr with their typical heterochromatic and inaccessible environment and (*iii*) Cys with its heterochromatic state and low NO.

## Discussion

Beyond the major role of tRNAs in protein synthesis and their multiple functions in other biological processes, the modulation of RNAP III transcription of the whole repertoire of tDNAs remains poorly documented^13^. Taken together, the findings described here provide evidence that DNA methylation and histone PTMs negatively affect RNAP III transcription of clustered tDNAs loci in Arabidopsis and demonstrate the influence of the epigenetic environment on the expression of tDNAs.

We report the differential expression levels of clustered *versus* dispersed copies of Ser, Tyr, Pro and Cys tDNAs in Arabidopsis. Indeed, under normal growth conditions, clustered tDNAs are repressed whilst their non-clustered equivalents are expressed. Most of these clustered tDNA loci contain the conserved *cis*-elements (A and B internal boxes, TA-rich and CAA upstream motifs, downstream poly-T stretches) involved in RNAP III transcriptional complex recognition, ruling out regulatory genetic alterations. Importantly, as clustered tRNAs (i) exhibit canonical cloverleaf tRNA structures (*e.g.* Supplementary Fig. 2) and (ii) possess all signatures involved in tRNA biogenesis to process them efficiently^51^, their lack of detection is unlikely caused by their rapid degradation following post-transcriptional events.

*In silico* analyses of epigenomic landscapes show that the silencing of clustered tDNAs is correlated with particular epigenetic states involving DNA methylation and repressive histone marks. As compared to dispersed tDNAs scattered in euchromatic environment, clustered tDNAs display high CG methylation levels suggesting that DNA methylation triggers their transcriptional repression as described for most repeats and TEs in Arabidopsis^52 53^. Indeed, epigenetic silencing at tDNA clusters is under the complex interplay between the DNA methyltransferases MET1, CMT2 and CMT3, as well as the chromatin remodeling factor DDM1 (Fig. 7). Such process shares some similarities with the transcriptional control of the heterochromatic 5S rDNA repeats^54^, strengthening the idea that DNA methylation efficiently prevents RNAP III transcription at several loci including clustered tDNAs. Conversely to the 5S rDNAs, we did not identify a predominant role neither for the RdDM pathway nor for the DNA methylation-independent pathway involving MOM1 in the transcriptional repression of clustered tDNAs. The particular interplay between genomic environments, epigenetic profiles and RNAP III transcription at clustered tDNAs loci would thus correspond to distinct and yet undetermined mechanisms (Fig. 7).

**Fig. 7.**
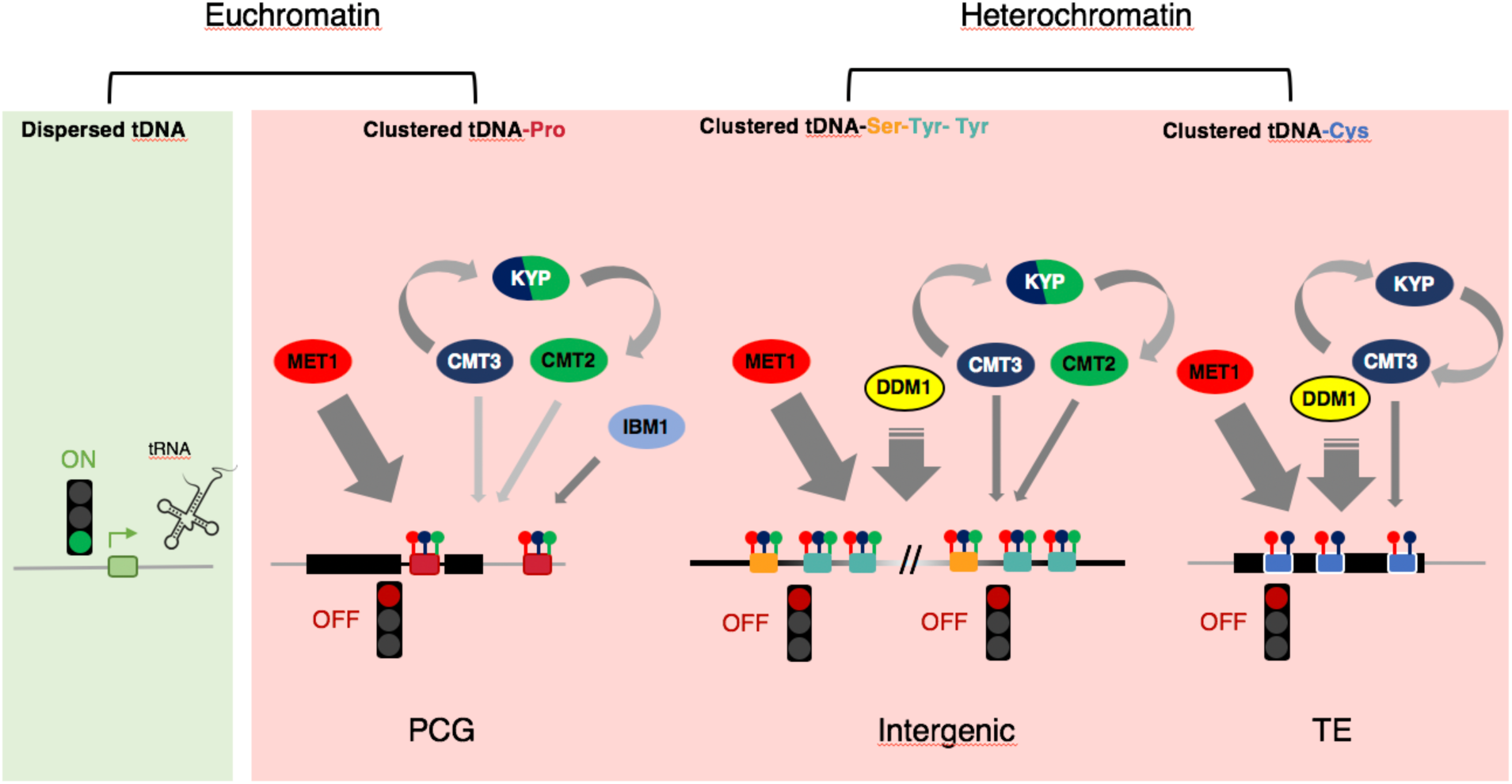
Model of epigenetic regulation of clustered tDNAs. Dispersed tDNAs are spread along the Arabidopsis chromosomes and clustered tDNAs repeats are located in distinct genomic (Chr 1, 2 and 5) and epigenomic (heterochromatic and euchromatic) environments. Dispersed tDNAs are found in a euchromatic permissive context allowing their expression (ON: Green light). Ser-Tyr-Tyr and Cys clusters overlap with heterochromatic state and are found within intergenic and TE genomic regions, respectively. Pro clusters overlap with euchromatic state and are located in the vicinity of protein coding genes (PCG) or within intron (black line) of PCG. These clustered tDNAs are found in a repressive chromatin context preventing their expression (OFF:Red light). In intergenic genomic regions, DNA methylation levels at Ser-Tyr-Tyr clusters is efficiently maintained through the combined actions of the DNA methyltransferases MET1 (CG context red circle), CMT3 (CHG context blue circle) and CMT2 (CHH context green circle). The H3K9 histone methyltransferase KYP allows directing DNA methylation in both CHG and CHH contexts via CMT3 and CMT2, respectively. In TE genomic regions, DNA methylation levels at Cys cluster is maintained through the predominant action of the DNA methyltransferases MET1 and CMT3. The H3K9 histone methyltransferase KYP allows directing DNA methylation in the CHG context via CMT3. In these heterochromatic clusters, the chromatin remodeling factor, DDM1, plays an important role to allow efficient maintenance of DNA methylation. In PCG region, DNA methylation levels at Pro clusters is maintained through the combined actions of the DNA methyltransferases MET1, and to a lower extent CMT2 and CMT3. The H3K9 histone methyltransferase KYP allows directing DNA methylation in both CHG and CHH contexts. Interestingly, the H3K9 histone demethylase, IBM1, likely acts to properly balance the H3K9me2 level and thus the CHG and CHH DNA methylation levels controlled by KYP, CMT3 and CMT2.

The Ser-Tyr-Tyr cluster is located like its dispersed counterparts in a large intergenic region. This region spans over about 40 Kbp on Chr1 and its DNA methylation at the three sequence contexts seems to rely on the DNA methylation maintenance effectors MET1, CMT2 and CMT3. In addition, the Ser-Tyr-Tyr cluster is embedded in an heterochromatic context enriched in nucleosomes and referred as CS8 (*i.e.* DNA methylation, H3K9me2 and H3K27me1), a chromatin state reported to map at intergenic regions^29^. Therefore, the repression of the Ser-Tyr-Tyr cluster may rely notably on the H3K9 histone methyltransferase KYP. Finally, the transcriptional release of clustered Tyr tDNAs in the *met1* mutant despite the ectopic gain of CHG and CHH methylations indicates that RNAP III can transcribe them in a high non-CG methylation context.

The silenced Pro clusters are located within introns of PCGs and/or overlapping with LncRNAs, while their expressed dispersed counterparts are predominantly found in intergenic regions, thus suggesting an impact of the genomic environment on the tDNA transcription. Also pinpointing that the unique genomic surrounding affects tRNA expression, intronic tRNAs were characterized by lower RNAP III occupancy in comparison with the RNAP III occupancy of non-intronic ones in *C. elegans*^55^. Regarding chromatin, Pro clusters are located within an euchromatic environment referred as CS7 (*i.e.* H3K4me1, H2Bub, and H3K36me3) typically found in intragenic regions and associated with transcription units longer than average^29^. In addition, Pro clusters display high CG DNA methylation levels and, conversely to the Ser-Tyr-Tyr cluster, both *ddm1* and *met1* mutations did not lead to the release of their expression. Thus, our finding suggests that additional epigenetic mechanisms may act to regulate the silencing of Pro clusters. In this sense, the histone H3K9 demethylase IBM1, known to prevent CHG and to some extent CHH methylation, was reported to bind to the long PCG (AT2G33980) in which one of the three Pro clusters is located^43^. Interestingly, this PCG displays several features characterizing IBM1/IBM2-EDM2 targets such as a long transcriptional unit (> 2 kbp), an alternative poly(A) site and intronic heterochromatin^43^. The control of H3K9me2 homeostasis by the antagonist action of IBM1 and KYP may therefore prevent gain of CHG and CHH methylation, thus establishing an epigenetic signature distinguishable from those of other tDNA clusters. Recent reports revealed the existence of a trade-off between heterochromatic TEs and gene functionality^56 57^. Also, heterochromatic introns containing clustered tDNAs behave similarly to the well characterized heterochromatic introns containing TE allowing the correct production of full transcripts^56^. Therefore, it is tempting to speculate that, at clustered Pro tDNAs, the fine tuning of H3K9me2 homeostasis involving KYP and IBM1 may help to balance the transcription of different mRNA isoforms and tRNA under particular growth conditions.

The Cys cluster is located on Chr5 within class II TEs from the Helitron family. As expected from its genetic environment, the Cys cluster resides in a constitutive heterochromatic environment (CS8) established by the DNA methyltransferases MET1 and CMT3, as well as the histone methyltransferase KYP. Despite its heterochromatic environment, the nucleosomal density at the Cys cluster was low compared to other clustered and non-clustered tDNAs. In the light of the repressive function of heterochromatin, a lower nucleosome occupancy is logically unexpected. Nucleosome depleted regions enriched in repetitive sequences were identified at heterochromatin in *S. pombe* and it was suggested that a wider spacing in heterochromatin may be more conducive to tighter forms of higher-order folding^58^. Supporting this, it has been proposed that tDNAs, as binding sites for TFs and architectural proteins, are playing important roles in the organization of the genome in animal and yeast nuclei^59^. Another clue is also the close genomic relationship of tDNAs with TEs as observed now for decades^60 61 62 63^. Given the ability of TE to be mobilized and to insert elsewhere in the genome as well as repeated sequences (*e.g.* tDNAs) to recombine, these TE-tDNA pairs might presumably be at the origin of both their numerous multi-copy dispersed isoacceptor families and clusters outstands^64^. Such genomic organization may reflect ancient transposition/recombination events followed by the establishment of repressive chromatin contexts leading to the silencing of both transposons and tDNAs.

The molecular mechanisms underlying non-coding RNA gene clusters formation and location are quite elusive. Genomic location of 5S rDNA loci are variable as well as their copy numbers^52^. Particular chromosomal rearrangements, such as unequal crossing overs and DNA break/repairs may have contributed to the formation and the shaping of tDNA clusters^65^. Moreover, considering the high variability of rRNA gene copy numbers through cell divisions in human and in plant clusters^66 67^, it seems that these particular genomic arrangements are submitted to a high flexibility. Interestingly, Arabidopsis plants defective for the cross link repair factor RTEL1 exhibit significant reduction of 45S rDNA copies^68^ highlighting that DNA repair factors also contribute to maintain repeats integrity. Addtionally, *ddm1* Arabidopsis plants display higher somatic and meiotic recombination frequencies^69^ suggesting that maintenance of both genome and methylome integrities are interconnected. Yet, it would be worth to know to which extend this dynamic could be applied to Arabidopsis tDNA repeat clusters. In *S. cerevisiae*, the role of tDNAs in genome instability through replication forks pausing was demonstrated, suggesting that such tDNA clusters might serve as motors for genome fitness and evolution^70^. Also, considering the link between chromatin status on replication properties^71^, this mechanism might also accompany genome instability or in contrary gently manage sensible genomic breakpoints at tDNA loci.

The biological relevance of the presence of repressed tDNA clusters may also reflect a role in the chromosome architecture. For example, certain tDNA clusters surrounding pericentromeres act as insulators preventing the spreading of silent heterochromatin in animal and yeast ^72^. Given the epigenomics and chromatin dynamics during plant development and upon stress conditions^73 74 75^ we can postulate that under certain developmental and/or growth conditions these tDNA clusters might be re-activated to sustain yet unknown functions related *e.g.* to adaptation to specific codon usage, regulation of protein synthesis or production of specific proteins such as proline-rich proteins^76^. Indeed, as the codon-tRNA balance is likely a major factor determining translation efficiency^77^, we can hypothesize that the shortage of a tRNA (*e.g.* Pro tRNA) would be a limiting factor for an optimal translation process to occur. Increasing the concentration of this tRNA by expressing clustered tDNAs would be a way to overcome this deleterious situation. Thus, it would be important to unveil the growth conditions releasing clustered tDNAs expression, if any.

## Materials and Methods

### Plant materials and growth conditions

*Arabidopsis thaliana* mutant plants used in this study are summarized in Supplementary Table 4 and are in the Col0 background. Plants were sown in batches on soil (Hawita Gruppe) supplied by a fertilizer N-P-K: 12-7-19 (Osmocote, 1g per 1L of compost) in 7 x 7 x 6.4 cm pots and cultured in growth chambers at 23°C/18°C for a photoperiod of 16h/8h (day/night).

For *in vitro* culture, after a 5-day stratification in darkness at 4°C, Col0 sterilized seeds were grown in Petri dishes (GBO, Reference: 664102) containing a MS222 (Duchefa Biochemie, Reference: M0222.0010) half-strength medium pH 5.8 supplemented with 0.5 g/L MES (Bio Basic Canada Inc., Reference: MB0341), 1% (w/v) sucrose, 0.68% (w/v) agar (Sigma Aldrich, Reference: 102067977), 250 μg/ml of sodium cefotaxime (Duchefa Biochemie, Reference: C0111) and 2 ml/L of Plant Preservative Mixture (Plant Cell Technology, PPM) in growth chambers 21°C/17°C for a photoperiod of 16h/8h (day/night).

### 5-azacytidine treatment

5-azacytidine (5-azaC) treatments were adapted from O. Mathieu et al.^44^. Col0 sterilized seeds were treated with 4 ml of water supplied with 250 μg/ml of sodium cefotaxime, 2 ml/L of PPM and either 0 mM, 0.25 mM, 0.5 mM, or 1 mM of 5-azaC freshly prepared every day. After germination, seeds were grown *in vitro* for 14 days.

### Total tRNA extraction and northern blot analysis

Total RNA was extracted from 22 days-old plants (except for *in vitro met1* plants) using TRI reagent (MRC) following the manufacturer instructions. For 4 days-old seedlings, roots, rosette leaves, inflorescences and green siliques of 7 week-old plants, total RNA was prepared according Chang et al.^78^. Total tRNA was enriched from total RNA as described in^33^. This protocol includes a LiCl precipitation step that allows enrichment, in the supernatant, of RNAs of a size smaller than 150 nt (*i.e*. 5S rRNA, tRNAs and small non-coding RNAs).

For tRNA northern blots^36^, up to 6μg of total tRNA was separated in 15% polyacrylamide gel, elecrotransferred onto Hybond N+ membrane (GE Healthcare Life Sciences) and hybridized to ^32^P radiolabeled oligonucleotides probes in 6X SSC, 0.5% (v/v) SDS at the following temperatures: Y14 P, Y32 P, P12 P, Ala P: 48°C, and P46 P: 44°C. Washing conditions were: 2 times 10 min in 2X SSC and 1 time 30 min in 2X SSC, 0.1% SDS, at hybridization temperature.

### tDNA and tRF analysis

Two small RNA libraries (GSM2301576, GSM2301577) were retrieved from the GEO database (http://www.ncni.nlm.nih.gov/geo/). tRF analysis was performed as described in Cognat et al.^33^. The adapter sequence was trimmed for the short read libraries with cutadapt (version 1.18)^79^. The data were mapped against the Arabidopsis tRNAs extracted from plantRNA database (http://plantrna.ibmp.cnrs.fr)^6^ with patMaN (version 1.2.2)^80^. The reads that can be assigned specifically to the clustered tRNAs and dispersed tRNAs have been counted to measure their expression.

The seqlogo of the upstream and downstream tDNA sequences were performed with WEBLOGO available at (https://weblogo.berkeley.edu).

### DNA methylation levels analysis

Published BS-Seq datasets (Supplementary Table 2) have been used to determine cytosine methylation levels of clustered and dispersed tDNA loci using 50 bp upstream (promoter) to 25 bp downstream (terminator) windows. Cytosine methylation levels in CG, CHG and CHH contexts were determined according to the method described in Daccord et al.^81^. Statistical analysis was performed using pairwise Wilcoxson tests. Graphical representations and pairwise Wilcoxson tests were done in R (https://www.r-project.org/).

### Chromatin states and nucleosome analyses

The genomic coordinates of Arabidopsis tDNAs were extracted from the plantRNA database^6^ and used to address the distribution of tDNAs across chromatin states. Genomic coordinates of the 9 chromatin states (CS) were downloaded from Sequeira-Mendes et al.^29^. Data for nucleosome positioning were downloaded from Liu et al.^48^ and converted in bigwig format using the UCSC wigToBigWig tool. A matrix of scores for dispersed and clustered tDNA regions (regions spanning from 50 bp upstream and 25 bp downstream) was calculated with the program computeMatrix from deeptools^82^ v. 3.1.3 (with parameter --binsize 5). The heatmaps were plotted with the plotHeatmap tool.

### Miscellaneous

The sequence of oligonucleotides used in this study are:

Yd:5’-CGGAAGACTGTAGATCTTTAGGTCGCTGGTTCGATTCCGGCAGG-3’

Yc:5’-CGGAGGACTGTAGATCCTTAGGTCACTGGTTCGAATCCGGTAGG-3’

Pd: 5’GCGAGAGGTCCCGAGTTCGATTCTCGGAACGCCCCCCA-3’

Pc: 5’-GCGAGAGGTCCCGAGTTCGATTCTCGGAATGCCCCCCA-3’

Y16 P: 5’-TGCCGGAATCGAACCAGCG-3’

Y34 P: 5’-TACCGGATTCGAACCAGTG-3’

P12 P: 5’-GGGCGTTCCGAGAAT-3’

P45 P: 5’-GGGCATTCCGAGAAT-3’

Ala P: 5’-ACCATCTGAGCTACATCCCC-3’

### Gel and blot images

Uncropped blots and gels are provided in Supplementary Fig. 9.

## Acknowledgements

The work was supported by the CNRS and the University of Strasbourg. We wish to thank Todd Blevins and Patryk Ngondo for helpful discussions.

## Author contributions

Study design: G.H., J.M., A.B. and L.D.; methylome analysis: S.G., D.P., V.C., G.H.; tRF and nucleosome analysis: V.C and A.B.; northern blot experiments: G.H., E.U.; writing: G.H., A.B., J.M and L.D.; supervision and funding acquisition: L.D.

## Additional information

Supplementary information is accessible online.

