## Supplemental data for "Epigenetic silencing of clustered tDNAs in Arabidopsis"

**Supplementary Fig. 1 a**, WEBLOGO analysis (http://weblogo.Berkeley.edu/) of the upstream and downstream regions of dispersed and clustered Tyr, Ser, Pro and Cys tDNAs. For each amino acid, the numbers of tDNAs used for the analysis are indicated. **b**, Percentage of AT nucleotides in the 50 nt upstream of pooled tDNAs. D and C stand for clustered and dispersed tDNAs respectively.

**a**

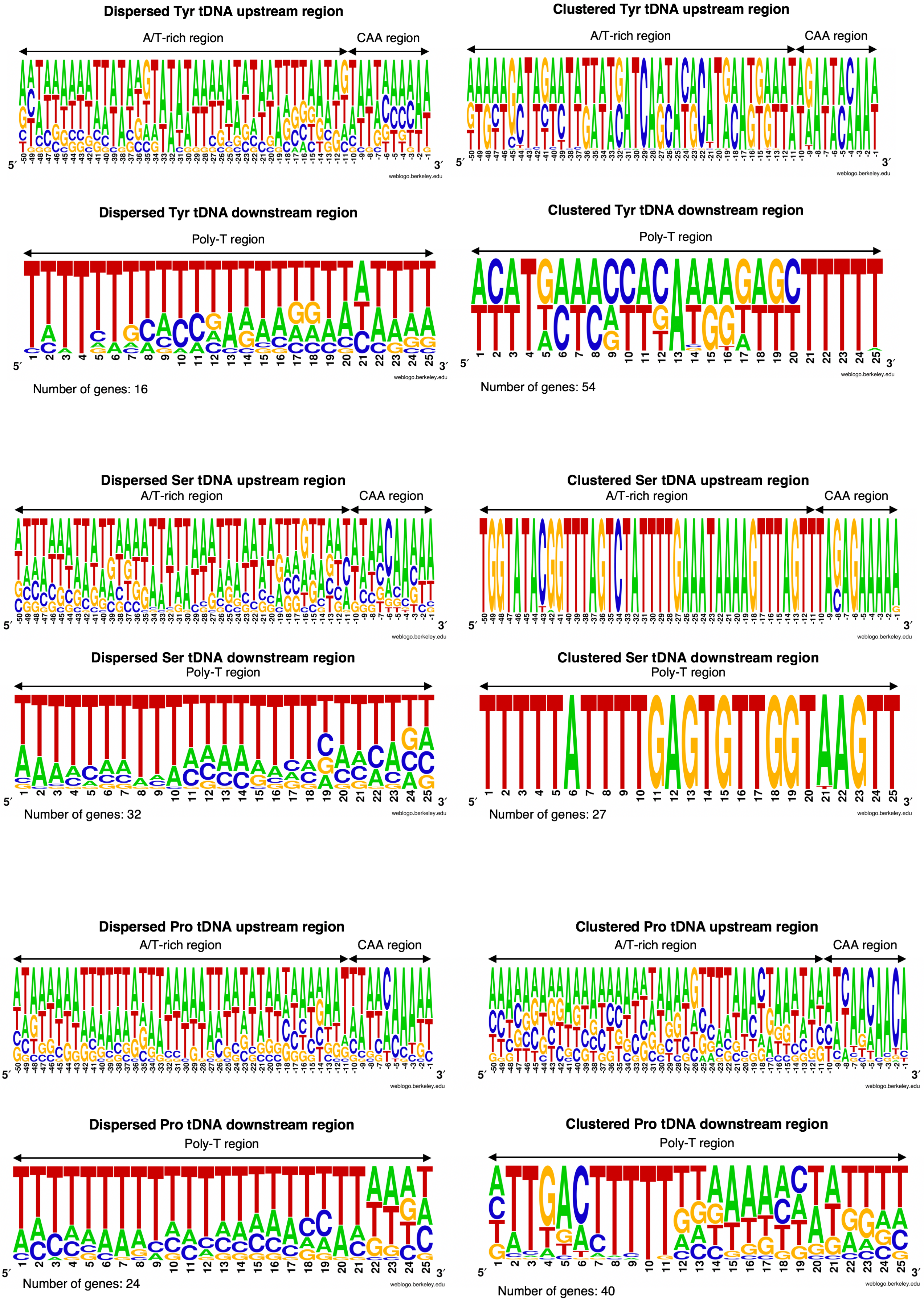

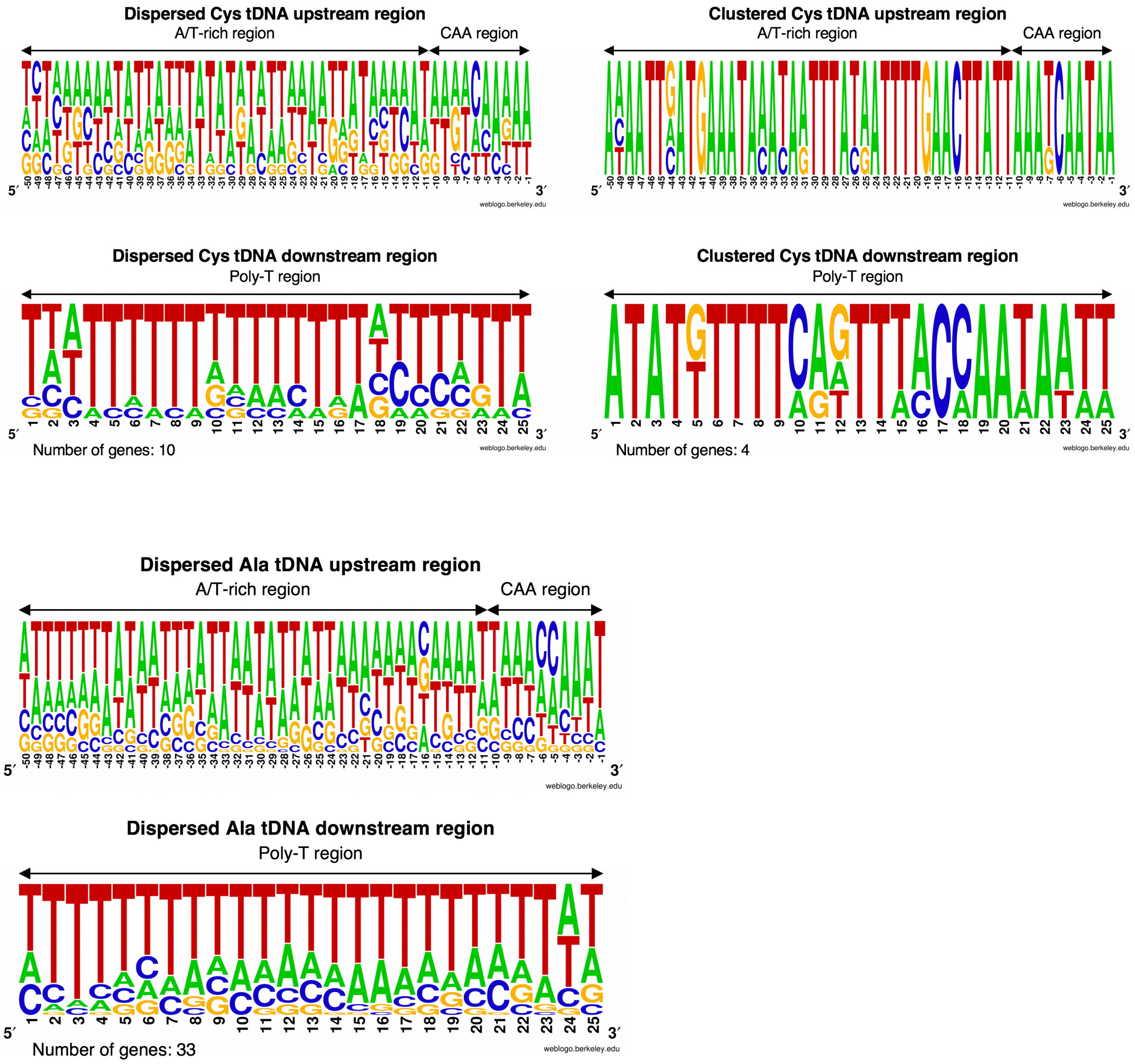
Dispersed Cys tDNA upstream region
A/T-rich region
CAA region
Clustered Cys tDNA upstream region
A/T-rich region
CAA region
Dispersed Cys tDNA downstream region
Poly-T region
Number of genes: 10
Clustered Cys tDNA downstream region
Poly-T region
Number of genes: 4
Dispersed Ala tDNA upstream region
A/T-rich region
CAA region
Dispersed Ala tDNA downstream region
Poly-T region
Number of genes: 33
5′
3′
weblogo.berkeley.edu

**b**

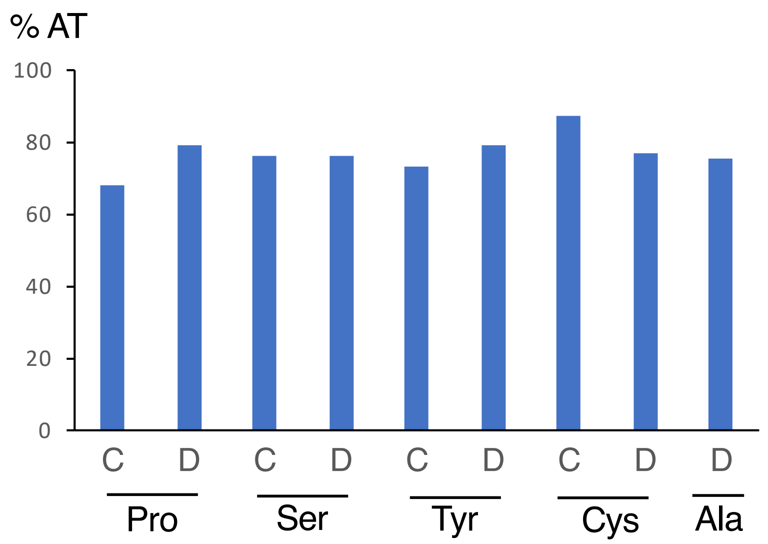
% AT
100
80
60
40
20
0
C
D
C
D
C
D
C
D
D
Pro
Ser
Tyr
Cys
Ala

**tRNA Tyr (Y) sequences**

| Sequences (positions marked: 49, 59, 65) | Genes Number | | Probes |
|---|---|---|---|
| CCGACCTTAGCTCAGTTGGTAGAGCGGAAGACT**GTA**AATCTTTAGGTCGCTGGTTCGATTCCGGCAGGTTGAACCA | 1 | dispersed | Y16 P |
| CCGACCTTAGCTCAGTTGGTAGAGCGGAAGACT**GTA**AATCTTTAGGTCGCTGGTTCGATTCCGG*CAGGTCGGACCA* | 1 | dispersed | Y16 P |
| CCGACCTTAGCTCAGTTGGTAGAGCGGAAGACT**GTA**GATCTTTAGGTCGCTGGTTCGATTCCGG*CAGGTCGGACCA* | 14 | dispersed | Y16 P |
| CCGACTTTAGCTCAGTTGGTAGAGCGGAGGACA**GTA**GATCCTTAGGTCACTGGTTCGAATCCGG*TAGGTCGGACCA* | 2 | clustered | Y34 P |
| CCGACCTTAGCTCAGTTGGTAGAGCGGAGGACT**GTA**GATCCTTAGGTCACTGGTTCGAATCCGG*TAGGTCGGACCA* | 32 | clustered | Y34 P |
| CCGACCTTAGCTCAGTTGGTAGAGCGGAGGACT**GTA**GATCCTTAGGTCATTGGTTCGAATCCGG*TAGGTCGGACCA* | 2 | clustered | |
| CCGACCTTAGCTCAGTTGGTAGAGCGGAGGACT**GTA**GATCCTTAGGTCATTGGTTCGAATCCGG*TAGGTCGGACCA* | 5 | clustered | |
| CCGACCTTAGCTCAGTTGGTAGAGCGGAGGACT**GTA**GATCCTTAGGTCATTGGTTCGAATCCGC*TAGGTCGGACCA* | 5 | clustered | |
| CCGACCTTAGCTCAGTTGGTAGAGCGGAGGACT**GTA**GATCCTTAGGTCACTGGTTCGAATTCGG*TAGGTCGGACCA* | 2 | clustered | |
| CCGACCTTAGCTCAGTTGGTAGAGCGGAGGACA**GTA**GATCCTTAGGTCATAGGTTCGAATCCGG*TAGGTCGGACCA* | 1 | clustered | |
| CCGACCTTAGCTCAGTTGGTAGAGCGGAGGACA**GTA**GATCCTTAGGTCACAGGTTCGAATCCGG*TAGGTCGGACCA* | 2 | clustered | |
| CCAACCTTAGCTCAGTTGGTAGAGCGGAGGACT**GTA**GATCCTTAGGTCACTGGTTCGAATTTGG*TAGGTCGGACCA* | 2 | clustered | |
| CCGACCTTAGCTCAGTTGGTAGAGCGGAGGACT**GTA**GATCCTTAGGTCACTGGTTCGAATCTGGTAGGTTTGACCA | 1 | clustered | |

**tRNA Pro (P) sequences**

| Sequences (position marked: 67) | Genes Number | | Probes |
|---|---|---|---|
| GGGCGTTTGGTCTAGTGGTATGATTCTCGCTT**TGG**GTGCGAGAGGTCCCGAGTTCGATTCTCGGAA*CGCCCCCCA* | 8 | dispersed | P12 P |
| GGGCGTTTGGTCTAGTGGTATGATTCTCGCTT**AGG**GTGCGAGAGGTCCCGAGTTCAATTCTCGGAA*CGCCCCCCA* | 4 | dispersed | P12 P |
| GGGTGTTTGGTCTAGTGGTATGATTCGCGCTT**CGG**GTGCGCGAGGTCCCGAGTTCGATTCTCGGAACACCCCCCA | 1 | dispersed | |
| GGGTGTTTGGTCTAGTGGTATGATTCTCGCTT**CGG**GTGCGAGAGGTCCCGAGTTCGATTCTCGGAACACCCCCCA | 3 | dispersed | |
| GGGCATTTGGTCTAGTGGTATGATTCTCGCTT**AGG**GTGCGAGAGGTCCGGAGTTCAATTCTCCGAA*TGCCCCCCA* | 1 | dispersed | |
| GGGCATTTGGTCTAGTGGTATGATTCTCGCTT**TGG**GTGCGAGAGGTCCCGAGTTCGATTCTCGGAA*TGCCCCCCA* | 1 | dispersed | P45 P |
| GGGCATTTGGTCTAGTGGTATGATTCTCGCTT**AGG**GTGCGAGAGGTCCCGAGTTCAATTCTCGGAA*TGCCCCCCA* | 3 6 | dispersed / clustered | P45 P |
| TGGCATTTGGTCTAGTGGTATGATTCTCGCTT**AGG**GTGCGAGAGGTCCCGAGTTCAATTCTCGGAA*TGCCCCCCA* | 1 | clustered | P45 P |
| GGGCATTTGGTCTAGTGGTATGATTCTCGCTT**CGG**GTGCGAGGGGTCCCGAGTTCGATTCTCGGAA*TGCCCCCCA* | 1 | clustered | P45 P |
| GGGCATTTGGTCTAGTGGTATGATTCTCGCTT**TGG**GTGCGAGAGGTCCCGAGTTCGATTCTCGGAA*TGCCCCCCA* | 33 | clustered | P45 P |
| GGGCATTTGGTCTAGTGGTATGATTCTCGCTT**TGG**GTGCGAGAGGTCCCGAGTTCGATTCTCGCAA*TGCCCCCCA* | 1 | clustered | P45 P |

**Supplementary Fig. 2a. Sequences alignment of Arabidopsis Tyrosine and Proline tDNAs.** Anticodons are in bold. Nucleotide differences are in red. Genes number for each sequence is indicated on the right. Black and red vertical bars indicate dispersed and clustered tDNAs respectively. tDNA regions chosen for probes (Y16 P, Y34 P, P12 P and P45 P) are underlined. Note that probe P45 P recognizes 38 dispersed proline tRNAs and 7 dispersed proline tRNAs. tDNA regions chosen for tRFs analysis are in italics. The nucleotidic positions of the polymorphisms in the regions used as probes and discriminating dispersed and clustered tRNAs are indicated.

**tRNA Ser(AGA) sequences**

| Sequences | Genes Number | |
|---|---|---|
| GTGGACGTGCCGGAGTGGT.T.ATCGGGCATGACT**AGA**AATCATGTGGGCTCTGCCCGCGCAGGTTCGAATCCTGCCGTTCACGCCA | 1 | |
| GTGGGCGTGCCGGAGTGGT.T.ATCGGGCATGACT**AGA**AATCATGTGGGCTTTGCCCGCGCAGGTTCGAATCCTGCCGCTCACGCCA | 1 | dispersed |
| GTAGGCGTGCCGGAGTGGT.T.ATCGGGCATGACT**AGA**AATCATGTGGGCTTTGCCCGCGCAGGTTCGAATCCTGCCG*CCTACGCCA* | 1 | |
| GTGGACGTGCCGGAGTGGT.T.ATCGGGCATGACT**AGA**AATCATGTGGGCTTTGCCCGCGCAGGTTCGAATCCTGCCGTTCACGCCA | 6 14 | |
| GTGGAAGTGGCGGAGTGGT.T.ATCGGGCATGACT**AGA**AATCATGTGGGCTTTGCCCGCGCAGGTTCGAATCCTGCCGTTCACGCCA | 2 | |
| GTGGACGTGCCGGAGTGGT.T.ATCGGGCATGACT**AGA**AATCATGTGGGTTTTGCCCGCGCAGGTTCGAATCCTGCCGTTCACGCCA | 2 | |
| GTGGACGTGCCGGAGTGGT.T.ATCGGGCATAACT**AGA**AATCATGTGGGCTTTGCCCGCGCAGGTTCGAATCCTGCCGTTCACGCCA | 1 | |
| GTGGAAGTGCCGGAGTGGT.T.ATCGGGAATGACT**AGA**AATCATGGGGGCTTTGCCCGCGCAGGTTTGAATCTTGCCG*TTTACGCCA* | 2 | clustered |
| GTGGACGTGCCGGAGTGGT.T.ATCGGGAATGACT**AGA**AATCATGGGGGCTTTGCCCGCGCAGGTTTGAATCTTGCCG*TTTACGCCA* | 3 | |
| GTGGACGTGCCGGAGTGGT.T.ATCGGGAATGACT**AGA**AATCATGGAGGCTTTGCCCGCGCACGTTTGAATCTTGCCG*TTTACGCCA* | 2 | |
| GTGGACATGCCGGAGTGGTGTTATCGGGCATAACT**AGA**AATCATGTGGGCTTTGCCCGCGCAGGTTCGAATCATGCCG*TTTACGCCA* | 2 | |

**tRNA Cys sequences**

| Sequences | Genes Number | |
|---|---|---|
| GGGTCCTTAGCTCAGTGGTAGAGCAATTGACT**GCA**GATCAATAGGTCACCGGTTCGAATCCGGTAGGGCCCTCCA | 2 | |
| GGGCTCATAGCTCAGTGGTAGAGCATTCGACT**GCA**GATCGAGAGGTCACCGGTTCGAACCCGGTTGGGCCCTCCA | 1 | |
| GAGCCTATAGCTCAGTGGTAGAGCAATTGACT**GCA**GATCAATAGGTCACCGGTTCGAATC*CGGTTGGGCTCTCCA* | 2 | dispersed |
| GGGTCCTTAGCTCAGTGGTAGAGCATTTGACT**GCA**GATCAAGAGGTCACCGGTTCGAACCCGGTAGGGCCCTCCA | 1 | |
| GGGTCCATAGCTCAGTGGTAGAGCATTTGACT**GCA**GATCAAGAGGTCACCGGTTCGAACCCGGTTGGGCCCTCCA | 1 | |
| GGGCCCATAGCTCAGTGGTAGAGCATTCGACT**GCA**GATCGAGAGGTCACCGGTTCGAACCCGGTTGAGCCCTCCA | 1 | |
| GGGTCCATAGCTCAGTGGTAGAGCAATTGACT**GCA**GATCAATAGGTCACCGGTTCGAACCCGGTTGGGCCCTCCA | 1 | |
| GGGTCCATAGCTCAGTGATAGAGCAATTGACT**GCA**GATCAATAGGTCACCGGTTCGAACCCGGTTGGGCCCTCCA | 1 1 | |
| GGGTTCATAGCTCAGTGGTAGAGCAATTGACT**GCA**GATCAATAGGTCACCGGTTCGAACCCGGTTGGGCCCTCCA | 1 | |
| AGGTCCATAGCTCAGTGGTAGAGCAATTGACT**GCA**GATCAATAGGTCACCGGTTTGAACC*TGGTTGGGCCCTCCA* | 1 | clustered |
| AGGTCCATAGCTCAGTGGTAGAGCAATTGACT**GCA**GATCAATAGGTCACCGGTTCGAACC*CGATTGGGCCCTCCA* | 1 | |

**Supplementary Fig. 2b. Sequences alignment of Arabidopsis Serine (AGA) and Cysteine tDNAs.** Anticodons are in bold. Nucleotide differences are in red. Genes number for each sequence is indicated on the right. Black and red vertical bars indicate dispersed and clustered tDNAs respectively. tDNA regions chosen for tRFs analysis are in italics.

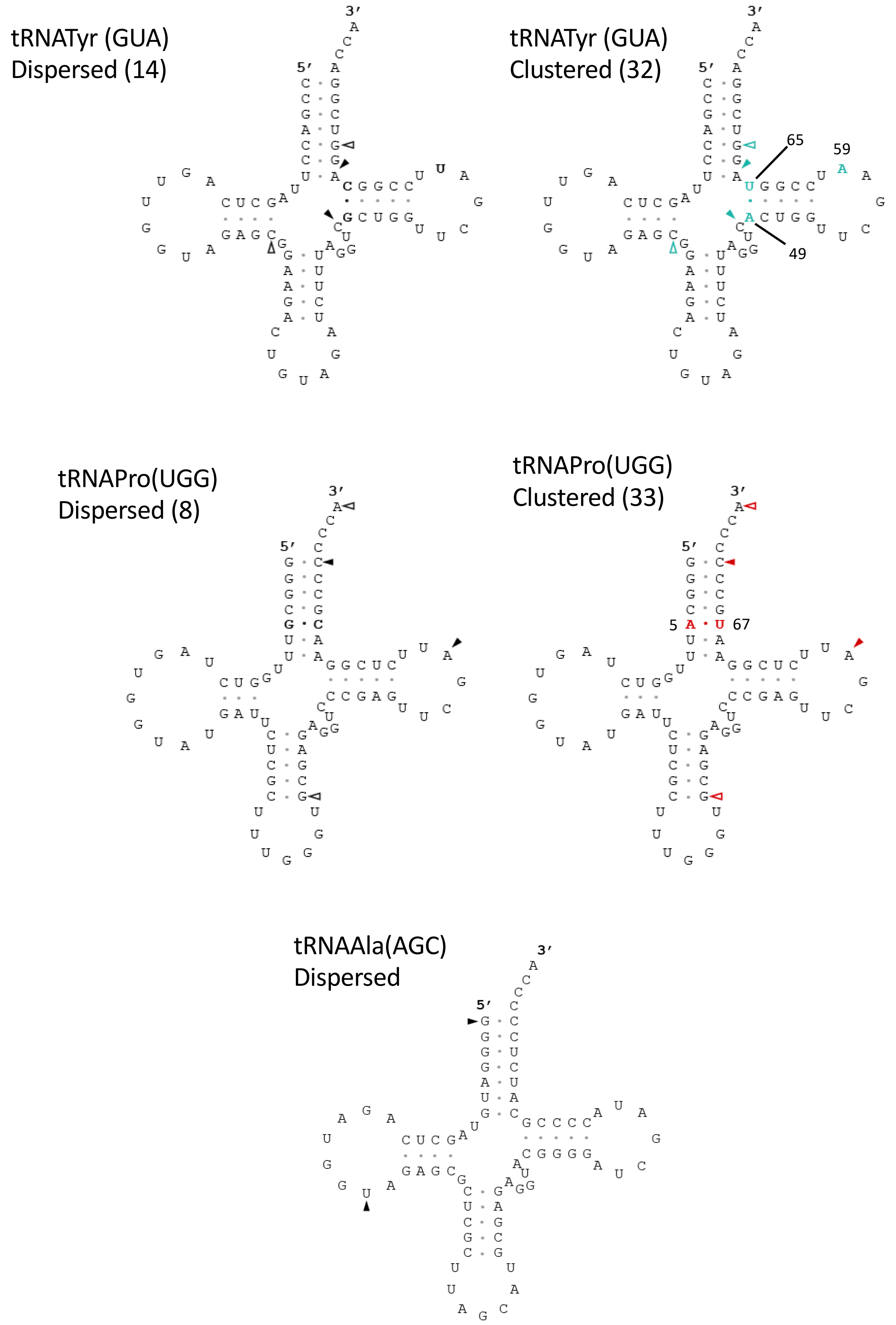


**Supplementary Fig. 3** Cloverleaf secondary structures of dispersed and clustered Tyr, Pro and Ala mature tRNAs. Sequences correspond to isodecoder tRNAs characterized by predominant gene copy numbers as reported on Supplementary Fig. 2. Empty and filled arrows indicate the extremities of the regions of the oligonucleotides used as controls and probes respectively as depicted in Fig. 2. Nucleotidic polymorphisms between dispersed and clustered tDNAs are in green (Tyr) or red (Pro) and their positions indicated. Number in parentheses indicate the number of tDNA copies.

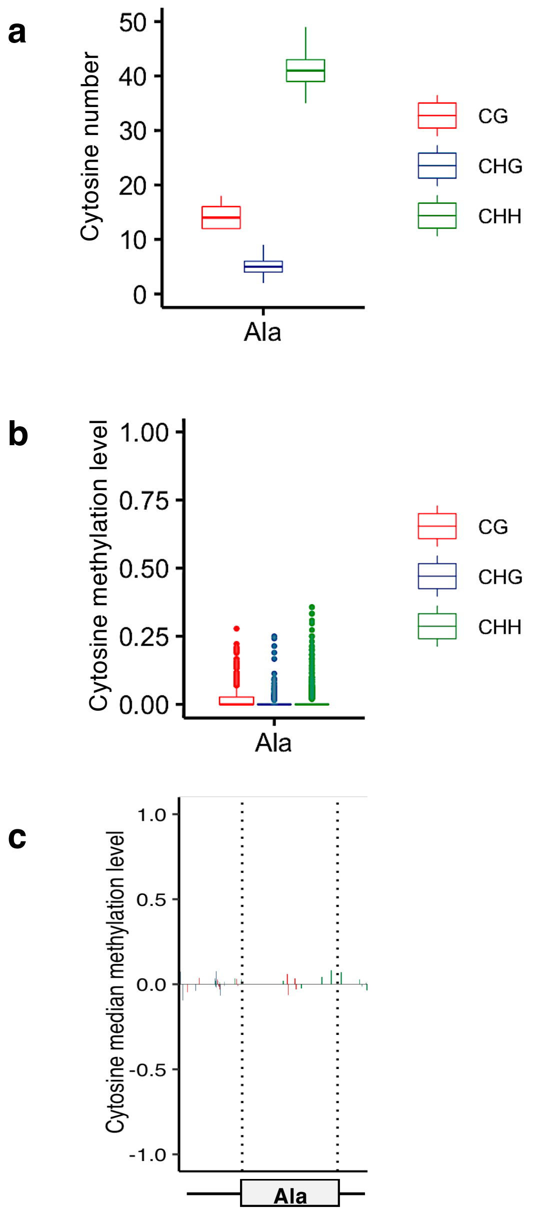


**Supplementary Fig. 4. Methylation landscape at dispersed Ala tDNA loci. a,** boxplot of DNA cytosine counts**. b,** boxplot of DNA cytosine methylation levels. **c**, DNA cytosine methylation median levels at single nucleotide resolution. Positive and negative values are referring to positive and negative DNA strands, respectively. The region corresponding to tDNA sequence is delimited by dotted lines.

**Supplementary Fig. 5** Heatmaps of DNA cytosine methylation levels for dispersed and clustered Pro, Ser, Tyr, Cys and Ala tDNAs upon Arabidopsis development. Each box corresponds to a single tDNA. Bubbles on the right indicate clustered tDNAs. CG heatmaps serve as reference for CHG and CHH ones.

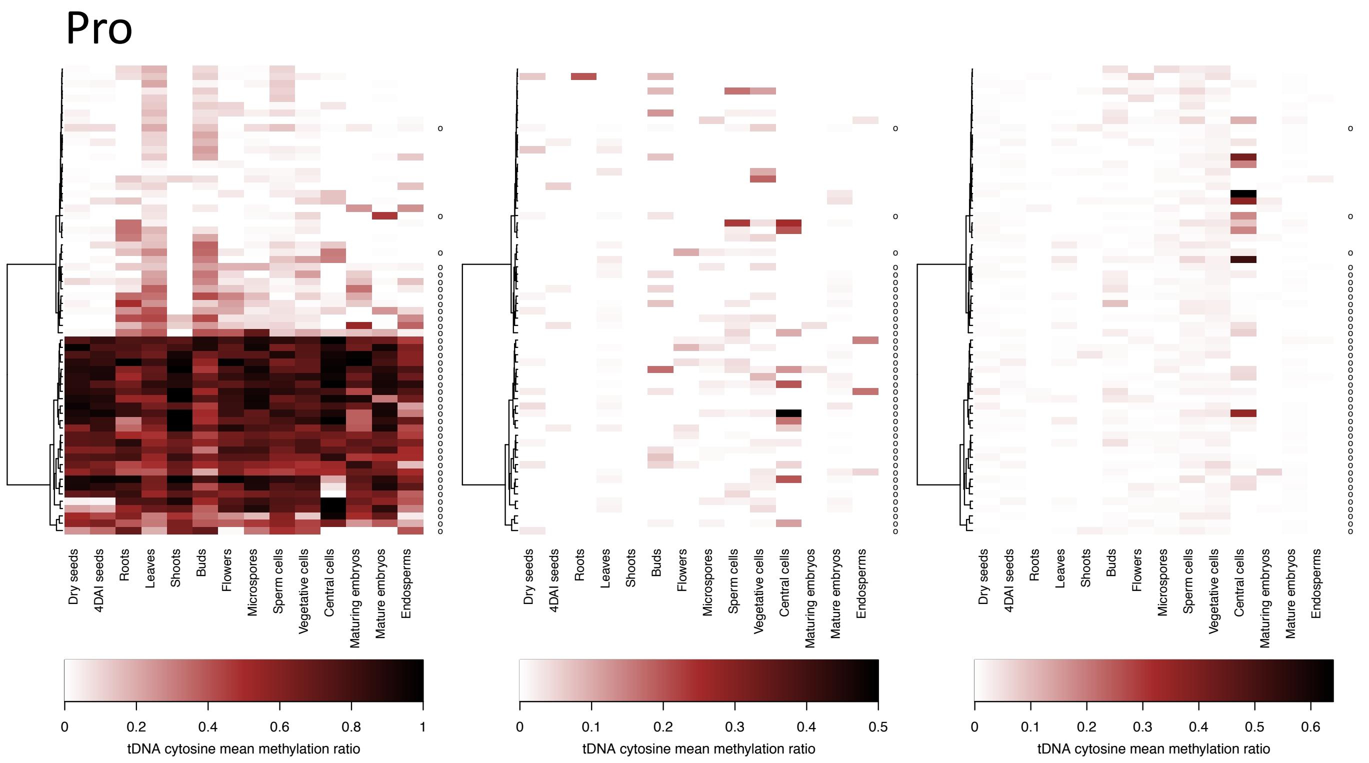


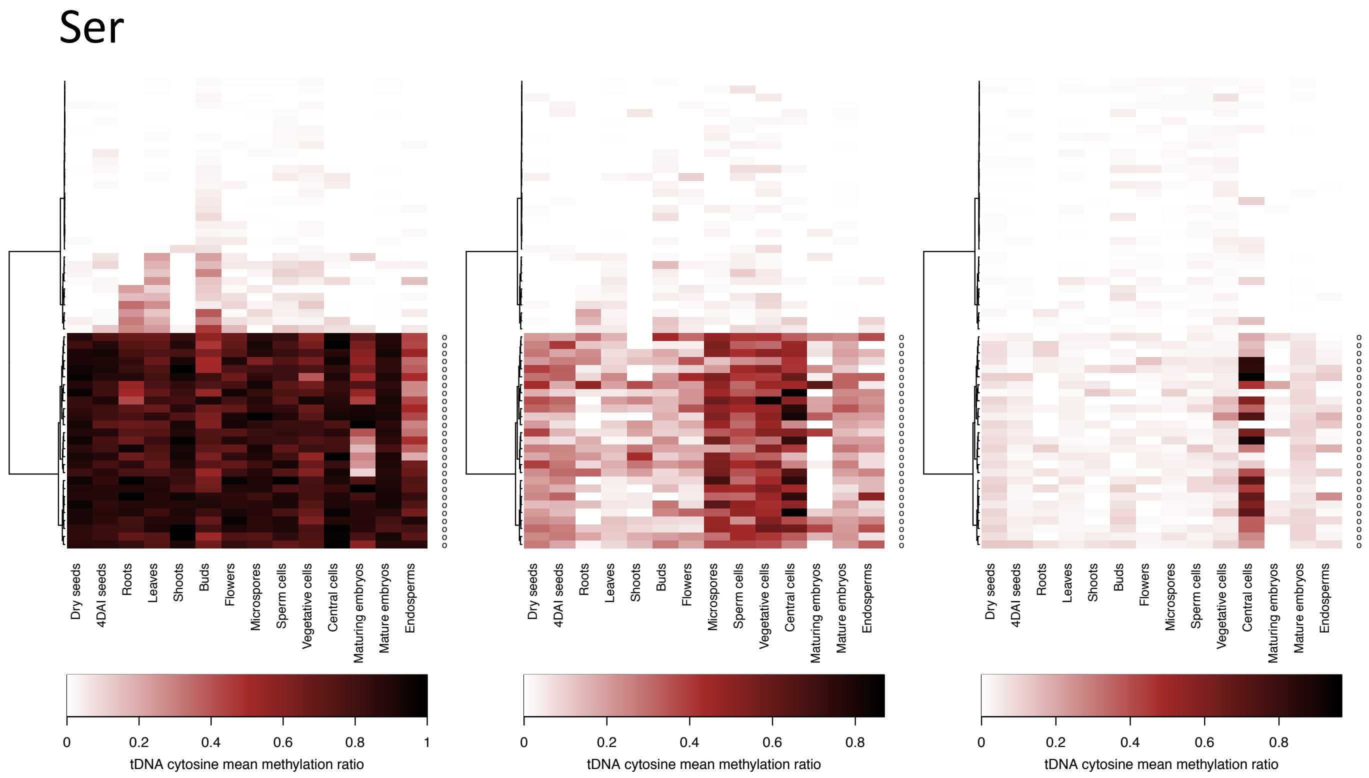

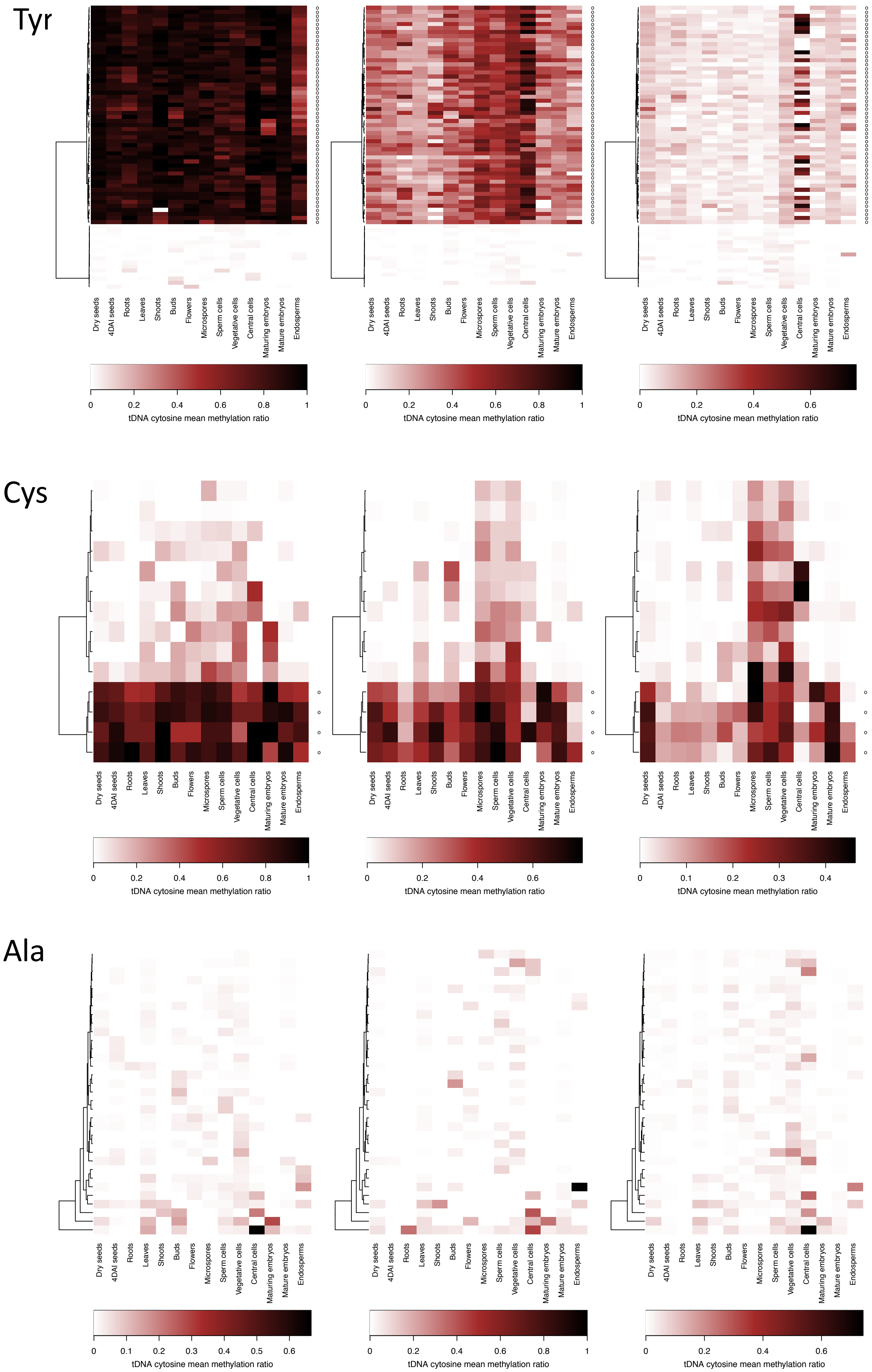
Tyr
Dry seeds
4DAI seeds
Roots
Leaves
Shoots
Buds
Flowers
Microspores
Sperm cells
Vegetative cells
Central cells
Maturing embryos
Mature embryos
Endosperms
0 0.2 0.4 0.6 0.8 1
tDNA cytosine mean methylation ratio
0 0.2 0.4 0.6 0.8 1
tDNA cytosine mean methylation ratio
0 0.2 0.4 0.6
tDNA cytosine mean methylation ratio
Cys
0 0.2 0.4 0.6 0.8 1
tDNA cytosine mean methylation ratio
0 0.2 0.4 0.6
tDNA cytosine mean methylation ratio
0 0.1 0.2 0.3 0.4
tDNA cytosine mean methylation ratio
Ala
0 0.1 0.2 0.3 0.4 0.5 0.6
tDNA cytosine mean methylation ratio
0 0.2 0.4 0.6 0.8 1
tDNA cytosine mean methylation ratio
0 0.2 0.4 0.6
tDNA cytosine mean methylation ratio

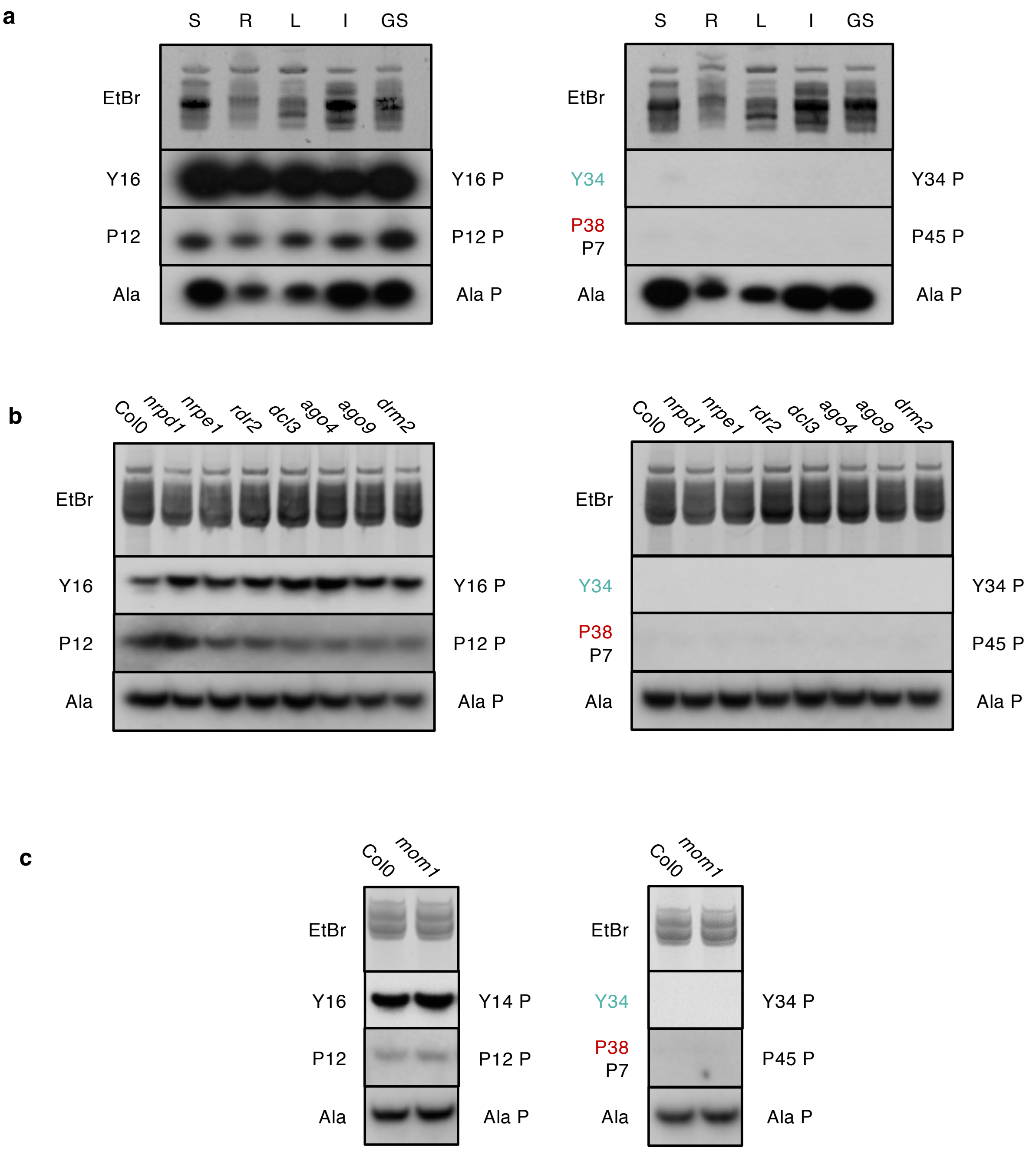


**Supplementary Fig. 6** Northern blot analyses of dispersed (left) and clustered (right) Tyr or Pro tRNAs. **a,** upon Arabidopsis development. S, seedlings; R, roots; L, leaves; I, inflorescences; GS, green siliques**. b**, in RdDM mutant plants. **c,** in *mom1* mutant plants. Staining with Ethidium Bromide (EtBr) and hybridization with a probe (Ala P) specific to Arabidopsis cytosolic Alanine tRNA were used as a loading control.

**Supplementary Fig. 7** RNAPII enrichment over clustered tDNA genes. Genome browser view of RNAPII enrichment over the five clustered tDNAs genomic regions identified in A. thaliana based on RNAPII ChiP-seq data from Liu et al.[48]. Clustered tDNAs genomic regions are presented below each track as in Fig. 1c. All tracks are equally scaled.

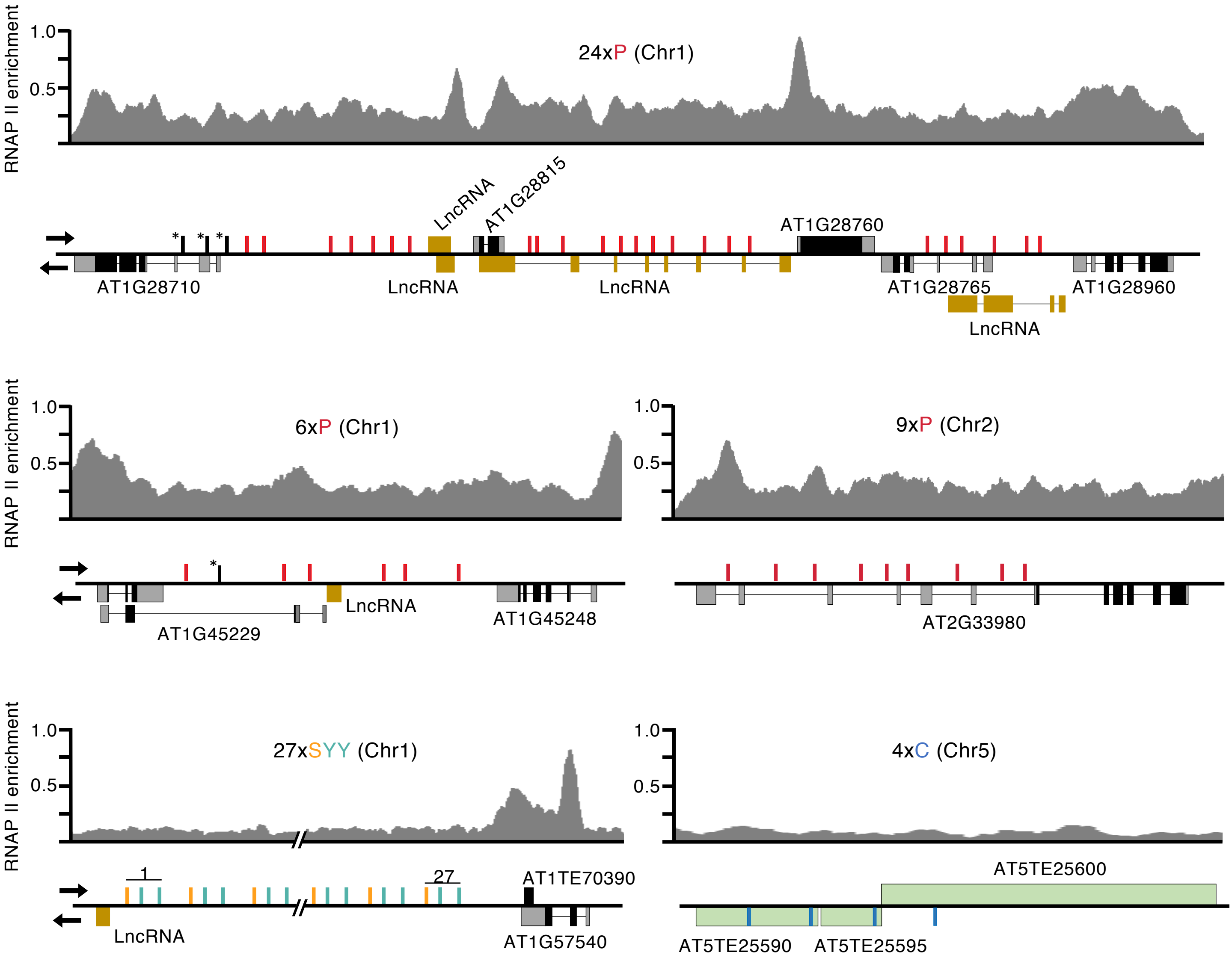

**Supplementary Fig. 8. Nucleosome occupancy at tDNA genes.** Heatmaps of nucleosome occupancy around clustered (C) and dispersed (D) tRNA genes in bins of 5 bp were derived from Liu et al[49]. For each tDNA, the region spanning from 50 bp upstream and 25 bp downstream is aligned on the X axis. Colors correspond to the value of data points at a given position. In general, nucleosome occupancy is indicated in blue and nucleosome depletion is indicated in red.

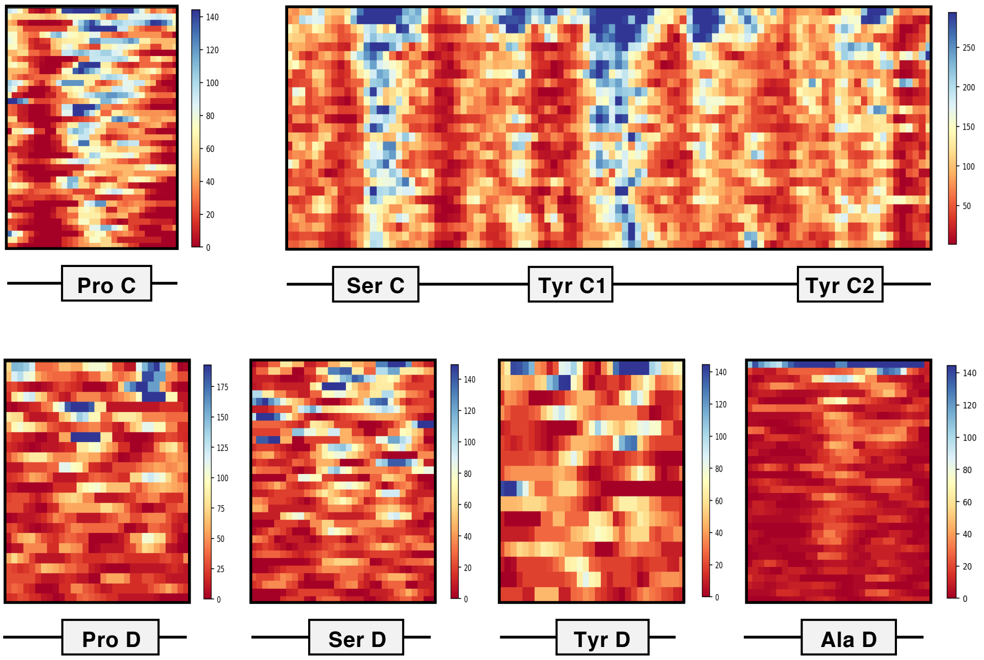

**Supplementary Figure 9.** Uncropped blots and gels.

Fig. 2b

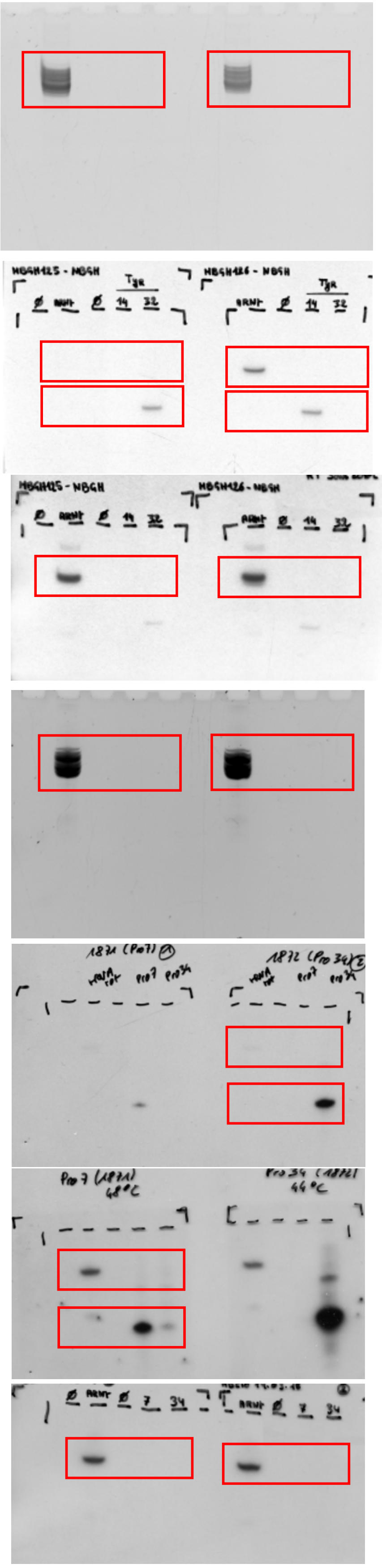

Fig. 5a

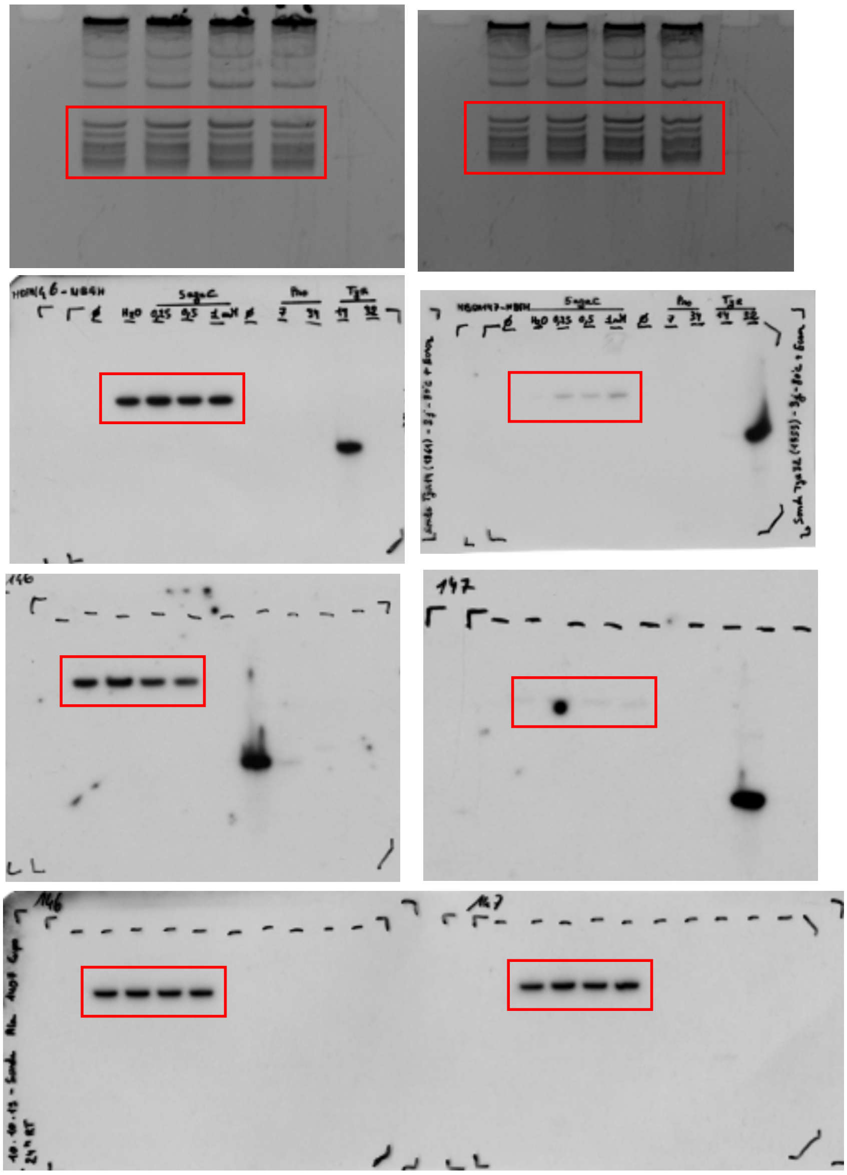

Fig. 5b (ddm1 – Y)

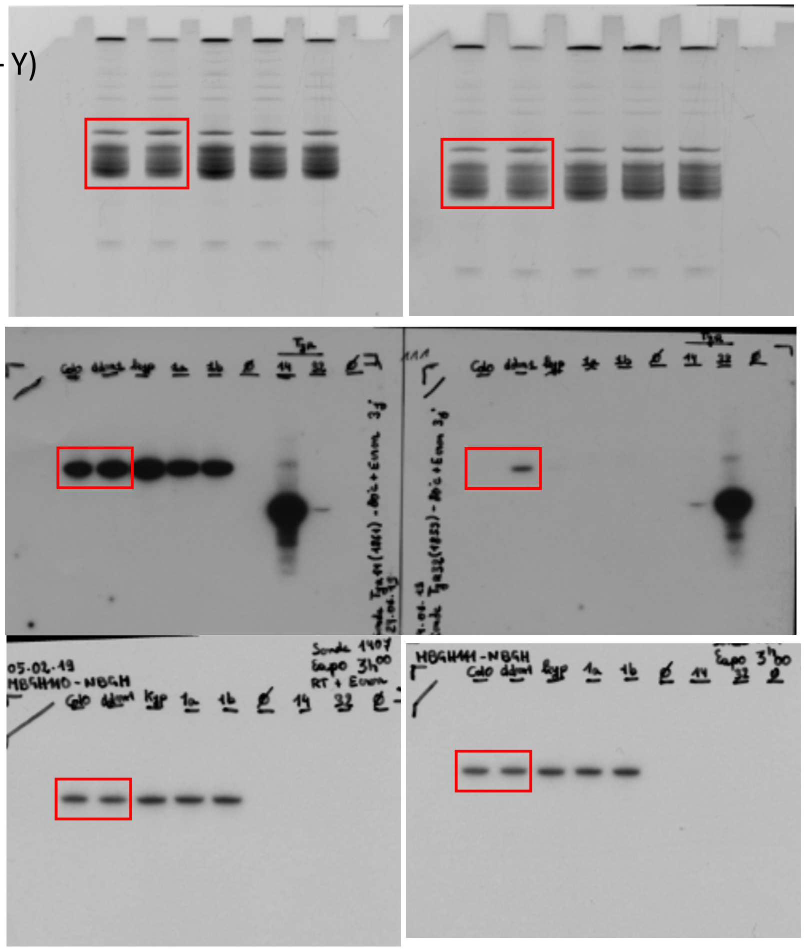

Fig. 5b
(ddm1- P)

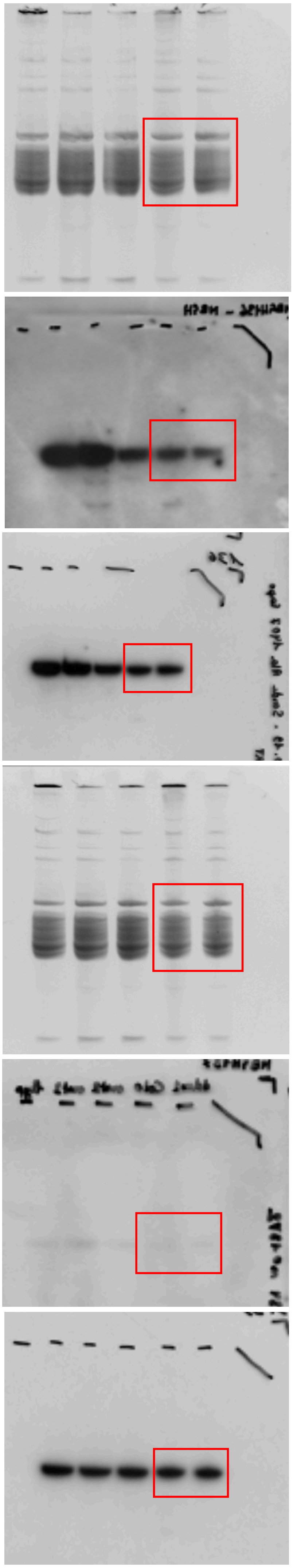

Fig. 5b
(met1- Y)

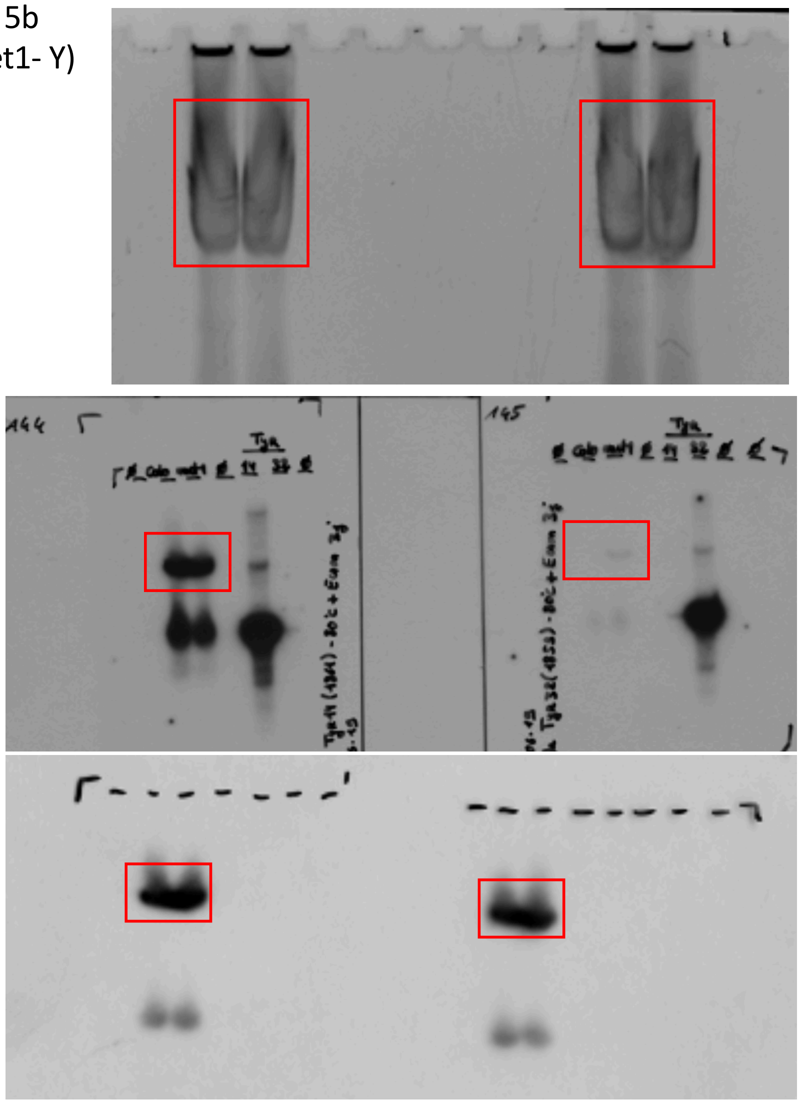

Fig. 5b
(met1- P)

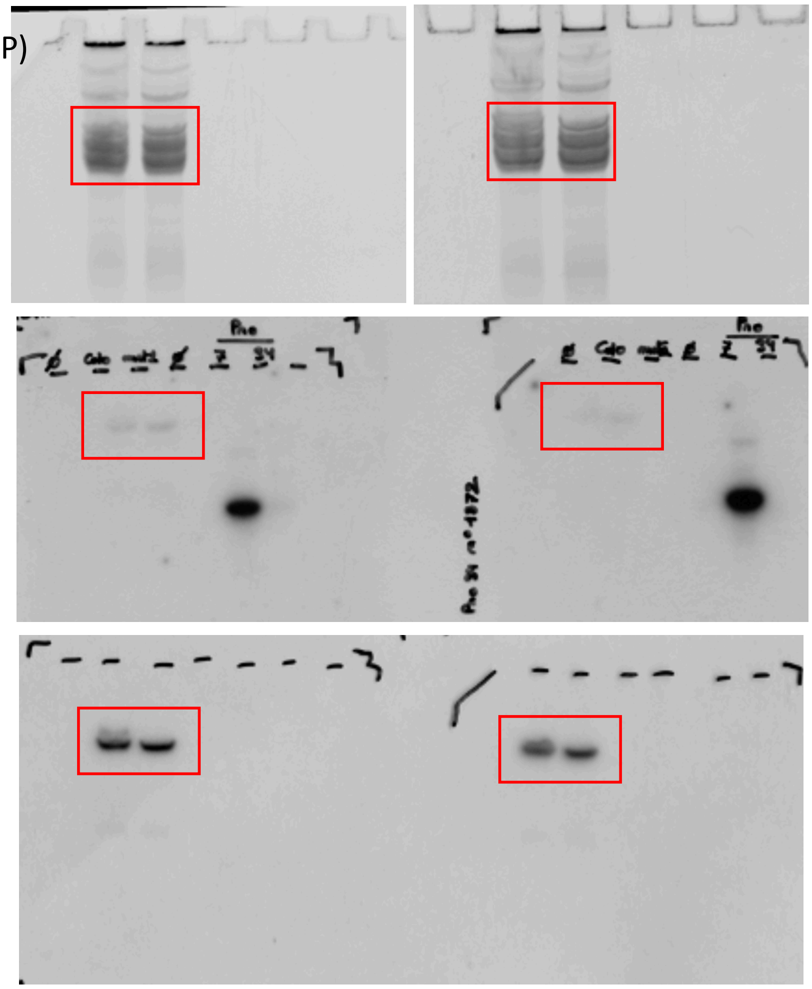

Fig. 5c

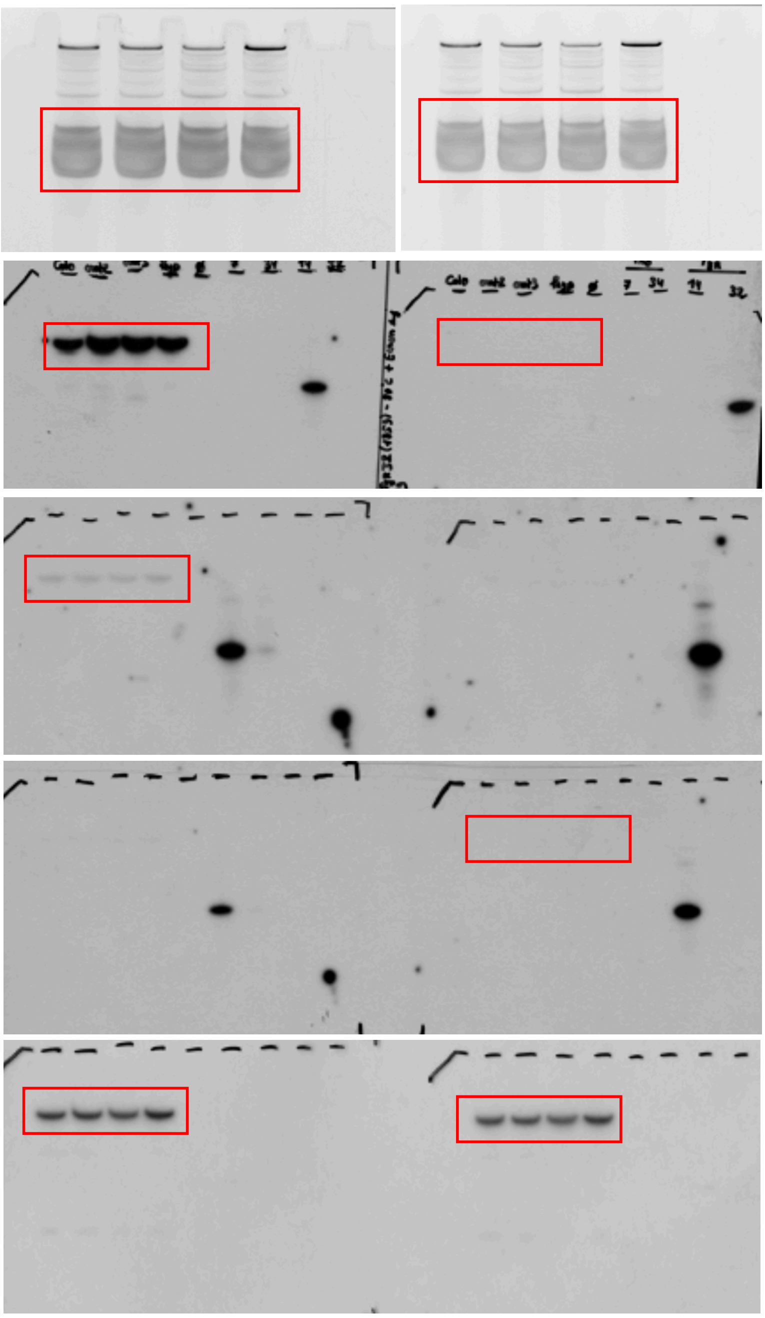

Fig. 5d

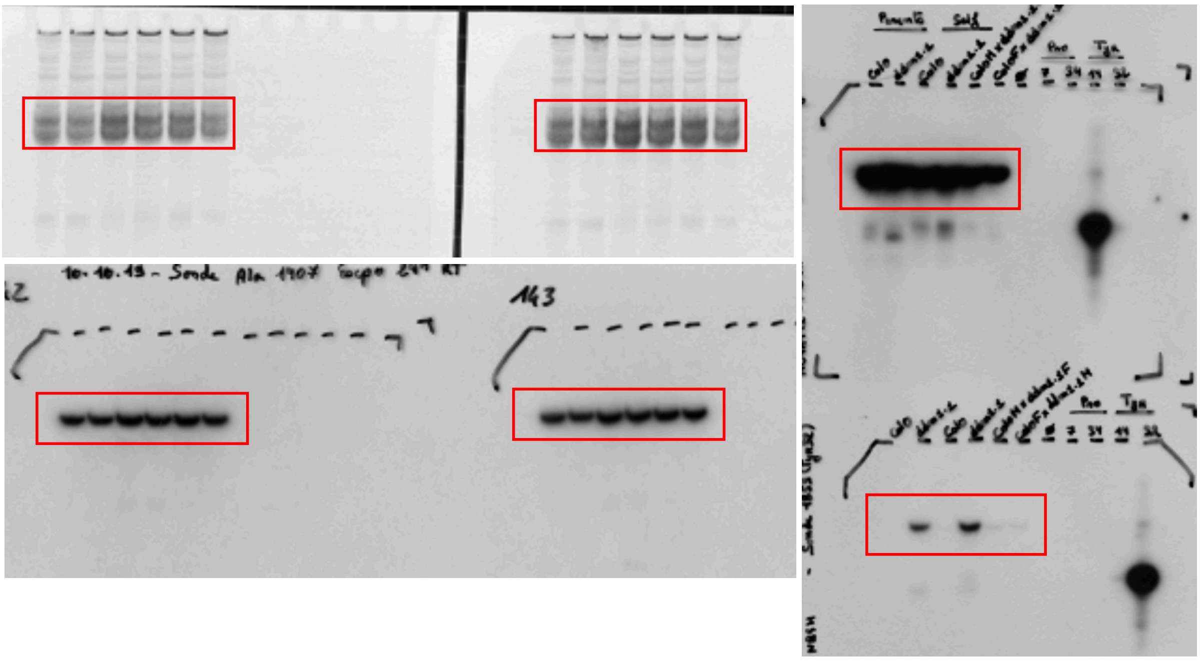

Sup Fig. 6a

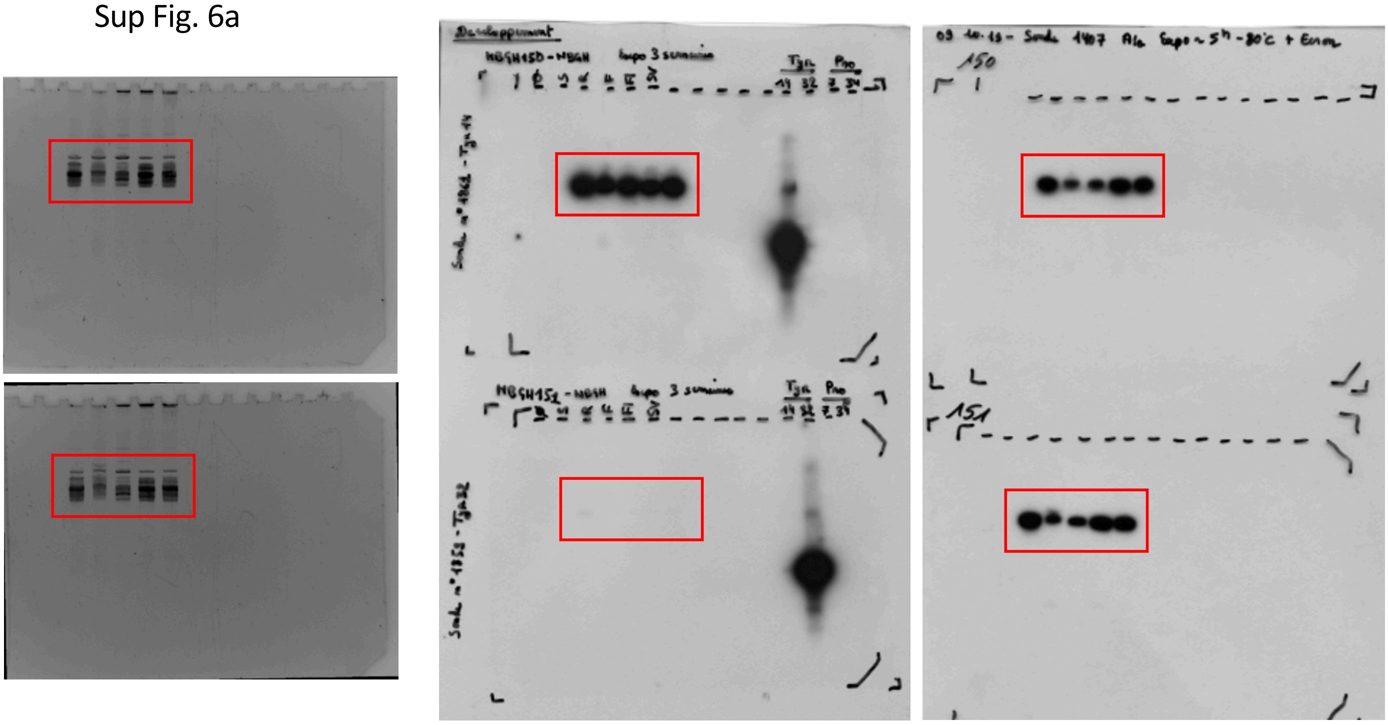

Sup Fig. 6b

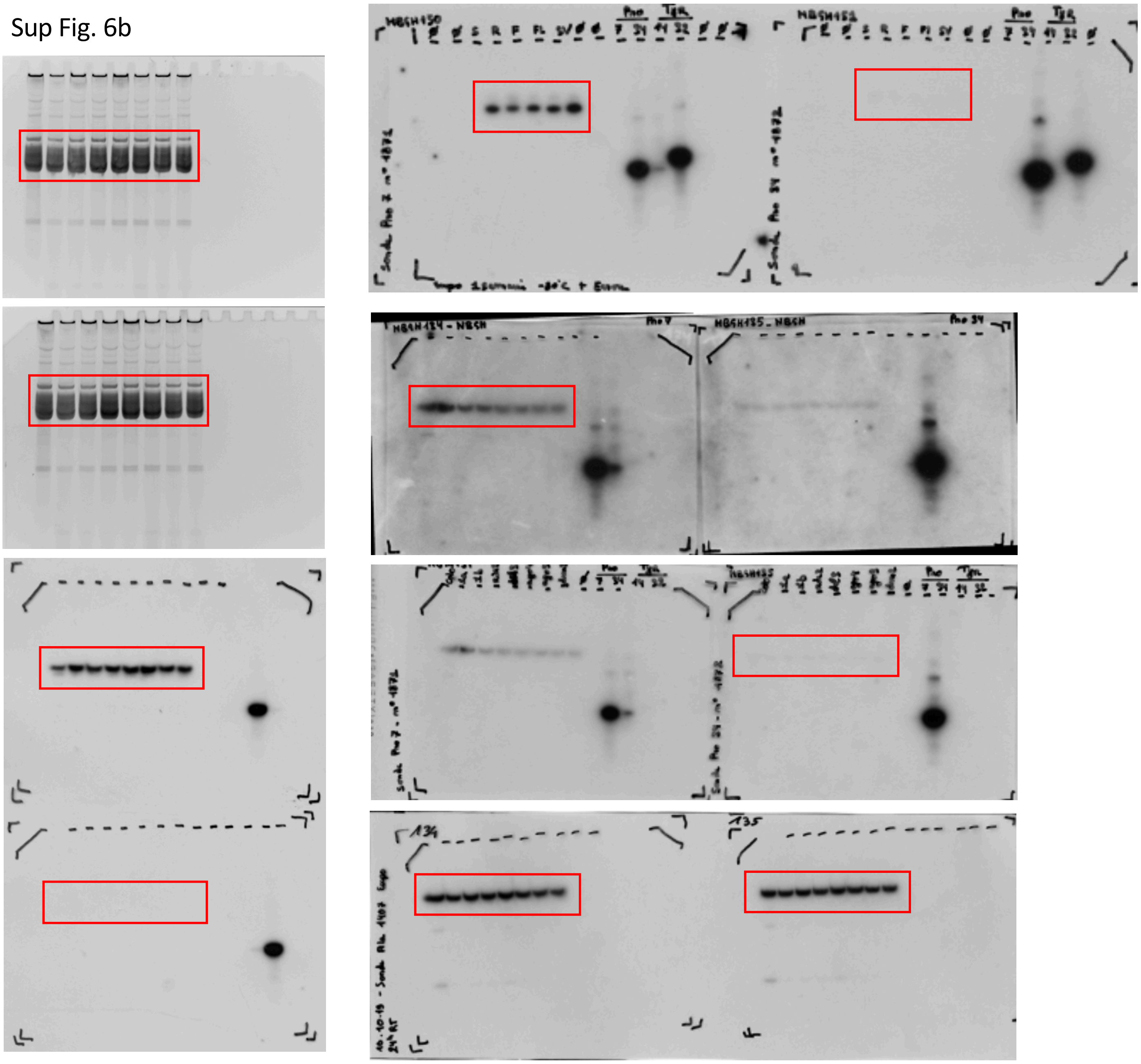

Sup Fig. 6c

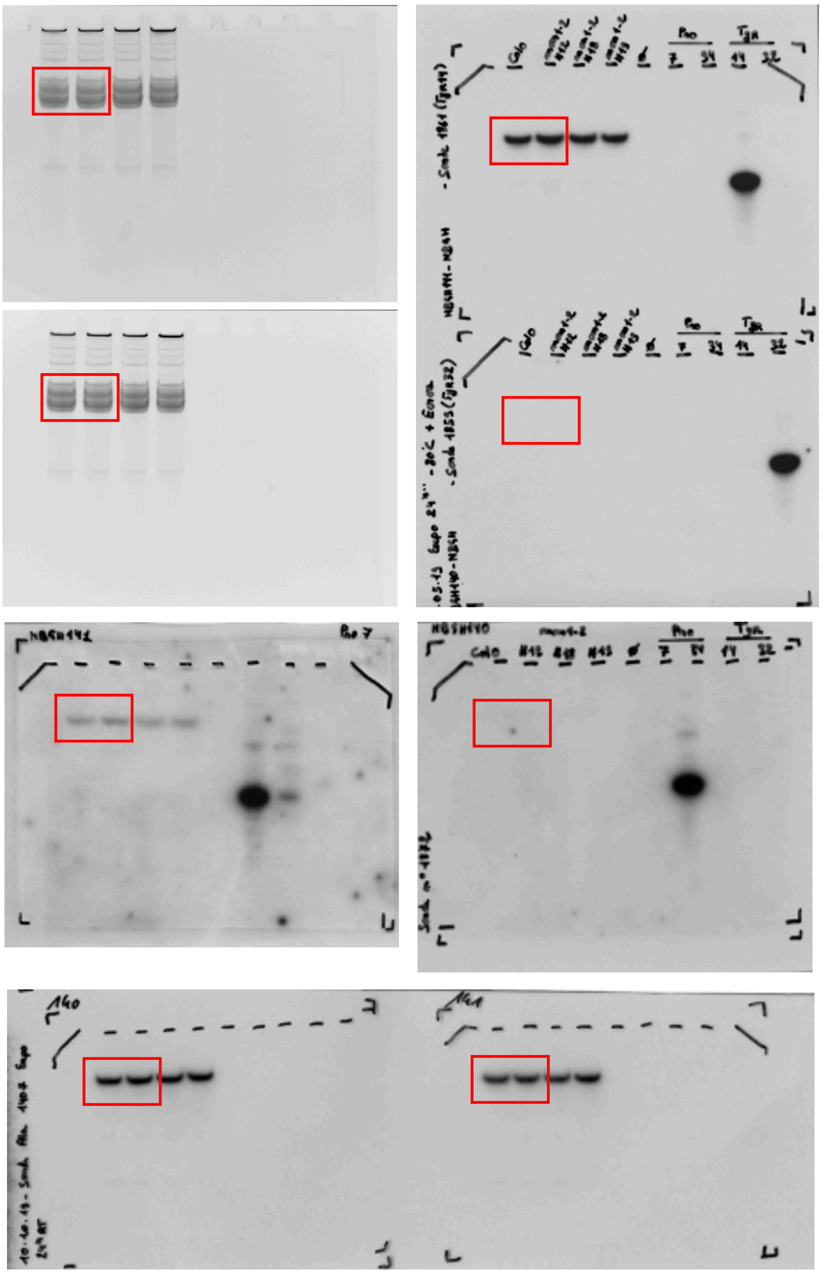

**Supplementary Table 1.** ***A. thaliana*** **genomic regions containing clustered tDNAs**

| Pro tDNAs - cluster 1 | Genomic_Feature | ID | Location | Strand | Start | End |
|---|---|---|---|---|---|---|
| | Nucleotide_diphospho_sugar_transferase_family_protein | AT1G28710.1 | Chr1 | Minus | 10086379 | 10088466 |
| | tRNA_(Pro_GGG)_pseudo_gene | AT1G44580.1 | Chr1 | Plus | 10088864 | 10088935 |
| | tRNA_(Pro_TGG)_pseudo_gene | AT1G28720.1 | Chr1 | Plus | 10089437 | 10089508 |
| | tRNA_(Pro_AGG)_pseudo_gene | AT1G28730.1 | Chr1 | Plus | 10089898 | 10089969 |
| | tRNA_(Pro_TGG)_gene | AT1G28740.1 | Chr1 | Plus | 10090368 | 10090439 |
| | tRNA_(Pro_TGG)_gene | AT1G28750.1 | Chr1 | Plus | 10090769 | 10090840 |
| | tRNA_(Pro_TGG)_gene | AT1G28770.1 | Chr1 | Plus | 10092325 | 10092396 |
| | tRNA_(Pro_TGG)_gene | AT1G28780.1 | Chr1 | Plus | 10092801 | 10092872 |
| | tRNA_(Pro_TGG)_gene | AT1G28790.1 | Chr1 | Plus | 10093309 | 10093380 |
| | tRNA_(Pro_TGG)_gene | AT1G28800.1 | Chr1 | Plus | 10093746 | 10093817 |
| | tRNA_(Pro_TGG)_gene | AT1G28810.1 | Chr1 | Plus | 10094173 | 10094244 |
| | lncRNA_gene | AT1G06173.1 | Chr1 | Plus | 10094622 | 10095145 |
| | lncRNA_gene | AT1G06177.1 | Chr1 | Minus | 10094805 | 10095222 |
| | Hypothetical_protein | AT1G28815.1 | Chr1 | Plus | 10095674 | 10096374 |
| | lncRNA_gene | AT1G28715.1 | Chr1 | Minus | 10095813 | 10102928 |
| | tRNA_(Pro_AGG)_gene | AT1G28820.1 | Chr1 | Plus | 10096951 | 10097022 |
| | tRNA_(Pro_TGG)_gene | AT1G28830.1 | Chr1 | Plus | 10097114 | 10097185 |
| | tRNA_(Pro_TGG)_gene | AT1G28840.1 | Chr1 | Plus | 10097718 | 10097789 |
| | tRNA_(Pro_TGG)_gene | AT1G28850.1 | Chr1 | Plus | 10098653 | 10098724 |
| | tRNA_(Pro_TGG)_gene | AT1G28860.1 | Chr1 | Plus | 10099079 | 10099150 |
| | tRNA_(Pro_AGG)_gene | AT1G28870.1 | Chr1 | Plus | 10099425 | 10099496 |
| | tRNA_(Pro_TGG)_gene | AT1G28880.1 | Chr1 | Plus | 10099827 | 10099898 |
| | tRNA_(Pro_TGG)_gene | AT1G28890.1 | Chr1 | Plus | 10100273 | 10100344 |
| | tRNA_(Pro_TGG)_gene | AT1G28900.1 | Chr1 | Plus | 10101028 | 10101099 |
| | tRNA_(Pro_TGG)_gene | AT1G28910.1 | Chr1 | Plus | 10101600 | 10101671 |
| | tRNA_(Pro_TGG)_gene | AT1G28920.1 | Chr1 | Plus | 10101993 | 10102064 |
| | Inner_nuclear_membrane_protein_A | AT1G28760.1 | Chr1 | Plus | 10103081 | 10104883 |
| | Hypothetical_protein | AT1G28765.1 | Chr1 | Minus | 10105047 | 10107645 |
| | tRNA_(Pro_TGG)_gene | AT1G28930.1 | Chr1 | Plus | 10106090 | 10106161 |
| | tRNA_(Pro_TGG)_gene | AT1G28940.1 | Chr1 | Plus | 10106528 | 10106599 |
| | misc_RNA_gene | AT1G09045.1 | Chr1 | Minus | 10106600 | 10109323 |
| | tRNA_(Pro_TGG)_gene | AT1G28950.1 | Chr1 | Plus | 10106898 | 10106969 |
| | tRNA_(Pro_TGG)_gene | AT1G28970.1 | Chr1 | Plus | 10107649 | 10107720 |
| | tRNA_(Pro_TGG)_gene | AT1G28980.1 | Chr1 | Plus | 10108389 | 10108460 |
| | tRNA_(Pro_TGG)_gene | AT1G28990.1 | Chr1 | Plus | 10108710 | 10108781 |

| | | | | | | |
|---|---|---|---|---|---|---|
| | Nudix_hydrolase_homolog_15 | AT1G28960.1 | Chr1 | Minus | 10109618 | 10111738 |
| **Pro tDNAs - cluster 2** | **Genomic_Feature** | **ID** | **Location** | **Strand** | **Start** | **End** |
| | Transmembrane_protein | AT1G45229.1 | Chr1 | Minus | 17154865 | 17156150 |
| | tRNA_(Pro_TGG)_gene | AT1G45234.1 | Chr1 | Plus | 17156607 | 17156678 |
| | tRNA_(Leu_AAG)_pseudogene | AT1G45236.1 | Chr1 | Plus | 17157360 | 17157431 |
| | tRNA_(Pro_TGG)_gene | AT1G45238.1 | Chr1 | Plus | 17158531 | 17158602 |
| | tRNA_(Pro_TGG)_gene | AT1G45240.1 | Chr1 | Plus | 17158979 | 17159050 |
| | lncRNA_gene | AT1G07313.1 | Chr1 | Minus | 17159423 | 17159704 |
| | tRNA_(Pro_TGG)_gene | AT1G45242.1 | Chr1 | Plus | 17160528 | 17160599 |
| | tRNA_(Pro_TGG)_gene | AT1G45244.1 | Chr1 | Plus | 17160969 | 17161040 |
| | tRNA_(Pro_TGG)_gene | AT1G45246.1 | Chr1 | Plus | 17162022 | 17162093 |
| | Nucleolar_histone_methyltransferase_related_protein | AT1G45248.1 | Chr1 | Minus | 17162963 | 17164779 |
| **Pro tDNAs - cluster 3** | **Genomic_Feature** | **ID** | **Location** | **Strand** | **Start** | **End** |
| | Nudix_hydrolase_homolog_22 | AT2G33980.1 | Chr2 | Minus | 14349764 | 14359901 |
| | tRNA_(Pro_TGG)_gene | AT2G33890.1 | Chr2 | Plus | 14350386 | 14350457 |
| | tRNA_(Pro_TGG)_gene | AT2G33900.1 | Chr2 | Plus | 14351359 | 14351430 |
| | tRNA_(Pro_TGG)_gene | AT2G33910.1 | Chr2 | Plus | 14352191 | 14352262 |
| | tRNA_(Pro_AGG)_gene | AT2G33920.1 | Chr2 | Plus | 14353143 | 14353214 |
| | tRNA_(Pro_TGG)_gene | AT2G33930.1 | Chr2 | Plus | 14353656 | 14353727 |
| | tRNA_(Pro_TGG)_gene | AT2G33940.1 | Chr2 | Plus | 14354098 | 14354169 |
| | tRNA_(Pro_CGG)_gene | AT2G33950.1 | Chr2 | Plus | 14355137 | 14355208 |
| | tRNA_(Pro_AGG)_gene | AT2G33960.1 | Chr2 | Plus | 14356055 | 14356126 |
| | tRNA_(Pro_TGG)_gene | AT2G33970.1 | Chr2 | Plus | 14356521 | 14356592 |
| **Ser/Tyr tDNAs -cluster** | **Genomic_Feature** | **ID** | **Location** | **Strand** | **Start** | **End** |
| | lncRNA_gene | AT1G08147.1 | Chr1 | Minus | 21268098 | 21268414 |
| | tRNA_(Ser_AGA)_gene | AT1G56730.1 | Chr1 | Plus | 21268787 | 21268868 |
| | tRNA_(Tyr_GTA)_gene | AT1G56740.1 | Chr1 | Plus | 21269122 | 21269206 |
| | tRNA_(Tyr_GTA)_gene | AT1G56750.1 | Chr1 | Plus | 21269552 | 21269636 |
| | tRNA_(Ser_AGA)_gene | AT1G56760.1 | Chr1 | Plus | 21270296 | 21270377 |
| | tRNA_(Tyr_GTA)_gene | AT1G56770.1 | Chr1 | Plus | 21270631 | 21270715 |
| | tRNA_(Tyr_GTA)_gene | AT1G56780.1 | Chr1 | Plus | 21271061 | 21271145 |
| | tRNA_(Ser_AGA)_gene | AT1G56790.1 | Chr1 | Plus | 21271805 | 21271886 |
| | tRNA_(Tyr_GTA)_gene | AT1G56800.1 | Chr1 | Plus | 21272140 | 21272224 |
| | tRNA_(Tyr_GTA)_gene | AT1G56810.1 | Chr1 | Plus | 21272570 | 21272654 |
| | tRNA_(Ser_AGA)_gene | AT1G56820.1 | Chr1 | Plus | 21273314 | 21273395 |
| | tRNA_(Tyr_GTA)_gene | AT1G56830.1 | Chr1 | Plus | 21273649 | 21273733 |
| | tRNA_(Tyr_GTA)_gene | AT1G56840.1 | Chr1 | Plus | 21274079 | 21274163 |
| | tRNA_(Ser_AGA)_gene | AT1G56850.1 | Chr1 | Plus | 21274823 | 21274904 |
| | tRNA_(Tyr_GTA)_gene | AT1G56860.1 | Chr1 | Plus | 21275158 | 21275242 |
| | tRNA_(Tyr_GTA)_gene | AT1G56870.1 | Chr1 | Plus | 21275609 | 21275693 |

| | | | | | |
|---|---|---|---|---|---|
| tRNA_(Ser_AGA)_gene | AT1G56880.1 | Chr1 | Plus | 21276352 | 21276433 |
| tRNA_(Tyr_GTA)_gene | AT1G56890.1 | Chr1 | Plus | 21276687 | 21276771 |
| tRNA_(Tyr_GTA)_gene | AT1G56900.1 | Chr1 | Plus | 21277117 | 21277201 |
| tRNA_(Ser_AGA)_gene | AT1G56910.1 | Chr1 | Plus | 21277861 | 21277942 |
| tRNA_(Tyr_GTA)_gene | AT1G56920.1 | Chr1 | Plus | 21278196 | 21278280 |
| tRNA_(Tyr_GTA)_gene | AT1G56930.1 | Chr1 | Plus | 21278647 | 21278731 |
| tRNA_(Ser_AGA)_gene | AT1G56940.1 | Chr1 | Plus | 21279390 | 21279471 |
| tRNA_(Tyr_GTA)_gene | AT1G56950.1 | Chr1 | Plus | 21279720 | 21279804 |
| tRNA_(Tyr_GTA)_gene | AT1G56960.1 | Chr1 | Plus | 21280178 | 21280262 |
| tRNA_(Ser_AGA)_gene | AT1G56970.1 | Chr1 | Plus | 21280922 | 21281003 |
| tRNA_(Tyr_GTA)_gene | AT1G56980.1 | Chr1 | Plus | 21281252 | 21281336 |
| tRNA_(Tyr_GTA)_gene | AT1G56990.1 | Chr1 | Plus | 21281710 | 21281794 |
| tRNA_(Ser_AGA)_gene | AT1G57000.1 | Chr1 | Plus | 21282454 | 21282535 |
| tRNA_(Tyr_GTA)_gene | AT1G57010.1 | Chr1 | Plus | 21282784 | 21282868 |
| tRNA_(Tyr_GTA)_gene | AT1G57020.1 | Chr1 | Plus | 21283242 | 21283326 |
| tRNA_(Ser_AGA)_gene | AT1G57030.1 | Chr1 | Plus | 21283986 | 21284067 |
| tRNA_(Tyr_GTA)_gene | AT1G57040.1 | Chr1 | Plus | 21284316 | 21284400 |
| tRNA_(Tyr_GTA)_gene | AT1G57050.1 | Chr1 | Plus | 21284774 | 21284858 |
| tRNA_(Ser_AGA)_gene | AT1G57060.1 | Chr1 | Plus | 21285519 | 21285600 |
| tRNA_(Tyr_GTA)_gene | AT1G57070.1 | Chr1 | Plus | 21285849 | 21285933 |
| tRNA_(Tyr_GTA)_gene | AT1G57080.1 | Chr1 | Plus | 21286306 | 21286390 |
| tRNA_(Ser_AGA)_gene | AT1G57090.1 | Chr1 | Plus | 21287050 | 21287131 |
| tRNA_(Tyr_GTA)_gene | AT1G57100.1 | Chr1 | Plus | 21287380 | 21287464 |
| tRNA_(Tyr_GTA)_gene | AT1G57110.1 | Chr1 | Plus | 21287837 | 21287921 |
| tRNA_(Ser_AGA)_gene | AT1G57120.1 | Chr1 | Plus | 21288581 | 21288662 |
| tRNA_(Tyr_GTA)_gene | AT1G57130.1 | Chr1 | Plus | 21288910 | 21288994 |
| tRNA_(Tyr_GTA)_gene | AT1G57140.1 | Chr1 | Plus | 21289368 | 21289452 |
| tRNA_(Ser_AGA)_gene | AT1G57150.1 | Chr1 | Plus | 21290112 | 21290193 |
| tRNA_(Tyr_GTA)_gene | AT1G57160.1 | Chr1 | Plus | 21290445 | 21290529 |
| tRNA_(Tyr_GTA)_gene | AT1G57170.1 | Chr1 | Plus | 21290903 | 21290987 |
| tRNA_(Ser_AGA)_gene | AT1G57180.1 | Chr1 | Plus | 21291461 | 21291542 |
| tRNA_(Tyr_GTA)_gene | AT1G57190.1 | Chr1 | Plus | 21291791 | 21291875 |
| tRNA_(Tyr_GTA)_gene | AT1G57200.1 | Chr1 | Plus | 21292248 | 21292332 |
| tRNA_(Ser_AGA)_gene | AT1G57210.1 | Chr1 | Plus | 21292992 | 21293073 |
| tRNA_(Tyr_GTA)_gene | AT1G57220.1 | Chr1 | Plus | 21293325 | 21293409 |
| tRNA_(Tyr_GTA)_gene | AT1G57230.1 | Chr1 | Plus | 21293783 | 21293867 |
| tRNA_(Ser_AGA)_gene | AT1G57240.1 | Chr1 | Plus | 21294341 | 21294422 |
| tRNA_(Tyr_GTA)_gene | AT1G57250.1 | Chr1 | Plus | 21294671 | 21294755 |
| tRNA_(Tyr_GTA)_gene | AT1G57260.1 | Chr1 | Plus | 21295128 | 21295212 |
| tRNA_(Ser_AGA)_gene | AT1G57270.1 | Chr1 | Plus | 21295872 | 21295953 |

| | | | | | |
|---|---|---|---|---|---|
| tRNA_(Tyr_GTA)_gene | AT1G57280.1 | Chr1 | Plus | 21296205 | 21296289 |
| tRNA_(Tyr_GTA)_gene | AT1G57290.1 | Chr1 | Plus | 21296663 | 21296747 |
| tRNA_(Ser_AGA)_gene | AT1G57300.1 | Chr1 | Plus | 21297221 | 21297302 |
| tRNA_(Tyr_GTA)_gene | AT1G57310.1 | Chr1 | Plus | 21297551 | 21297635 |
| tRNA_(Tyr_GTA)_gene | AT1G57320.1 | Chr1 | Plus | 21298009 | 21298093 |
| tRNA_(Ser_AGA)_gene | AT1G57330.1 | Chr1 | Plus | 21298753 | 21298834 |
| tRNA_(Tyr_GTA)_gene | AT1G57340.1 | Chr1 | Plus | 21299086 | 21299170 |
| tRNA_(Tyr_GTA)_gene | AT1G57350.1 | Chr1 | Plus | 21299544 | 21299628 |
| tRNA_(Ser_AGA)_gene | AT1G57360.1 | Chr1 | Plus | 21300102 | 21300183 |
| tRNA_(Tyr_GTA)_gene | AT1G57370.1 | Chr1 | Plus | 21300432 | 21300516 |
| tRNA_(Tyr_GTA)_gene | AT1G57380.1 | Chr1 | Plus | 21300890 | 21300974 |
| tRNA_(Ser_AGA)_gene | AT1G57390.1 | Chr1 | Plus | 21301634 | 21301715 |
| tRNA_(Tyr_GTA)_gene | AT1G57400.1 | Chr1 | Plus | 21301967 | 21302051 |
| tRNA_(Tyr_GTA)_gene | AT1G57410.1 | Chr1 | Plus | 21302425 | 21302509 |
| tRNA_(Ser_AGA)_gene | AT1G57420.1 | Chr1 | Plus | 21302983 | 21303064 |
| tRNA_(Tyr_GTA)_gene | AT1G57430.1 | Chr1 | Plus | 21303317 | 21303401 |
| tRNA_(Tyr_GTA)_gene | AT1G57440.1 | Chr1 | Plus | 21303777 | 21303861 |
| tRNA_(Ser_AGA)_gene | AT1G57450.1 | Chr1 | Plus | 21304520 | 21304603 |
| tRNA_(Tyr_GTA)_gene | AT1G57460.1 | Chr1 | Plus | 21304856 | 21304935 |
| tRNA_(Tyr_GTA)_gene | AT1G57470.1 | Chr1 | Plus | 21305311 | 21305395 |
| tRNA_(Ser_AGA)_gene | AT1G57480.1 | Chr1 | Plus | 21306055 | 21306136 |
| tRNA_(Tyr_GTA)_gene | AT1G57490.1 | Chr1 | Plus | 21306389 | 21306473 |
| tRNA_(Tyr_GTA)_gene | AT1G57500.1 | Chr1 | Plus | 21306849 | 21306933 |
| tRNA_(Ser_AGA)_gene | AT1G57510.1 | Chr1 | Plus | 21307592 | 21307675 |
| tRNA_(Tyr_GTA)_gene | AT1G57520.1 | Chr1 | Plus | 21307928 | 21308012 |
| tRNA_(Tyr_GTA)_gene | AT1G57530.1 | Chr1 | Plus | 21308387 | 21308471 |
| 40S_ribosomal_protein | AT1G57540.1 | Chr1 | Minus | 21309895 | 21311521 |
| Transposable_element_gene | AT1TE70390 | Chr1 | Plus | 21309950 | 21310162 |

**Cys tDNAs - cluster**

| Genomic_Feature | ID | Location | Strand | Start | End |
|---|---|---|---|---|---|
| Transposable_element_gene | AT5TE25590 | Chr5 | Minus | 7074005 | 7073198 |
| tRNA_(Cys_GCA)_gene | AT5G20852.1 | Chr5 | Minus | 7073601 | 7073530 |
| tRNA_(Cys_GCA)_gene | AT5G20854.1 | Chr5 | Minus | 7074014 | 7073943 |
| Transposable_element_gene | AT5TE25595 | Chr5 | Minus | 7074433 | 7074006 |
| tRNA_(Cys_GCA)_gene | AT5G20856.1 | Chr5 | Minus | 7074427 | 7074356 |
| Transposable_element_gene | AT5TE25600 | Chr5 | Plus | 7074434 | 7076706 |
| tRNA_(Cys_GCA)_gene | AT5G20858.1 | Chr5 | Minus | 7074843 | 7074772 |

**Supplementary Table 2. Publicly available methylomes used in this work.**

| Sample | Ecotype | SRR file | Reference |
|---|---|---|---|
| Dry seeds | Col0 | SRR5239966 | 36 |
| 4DAI seeds | Col0 | SRR5239976 | 36 |
| Roots | Col0 | SRR578936 | 22 |
| Shoots | Col0 | SRR578938 | 22 |
| Leaves | Col0 | SRR988547 | 37 |
| Unopened flower buds | Col0 | SRR013306 | 38 |
| Flowers | Col0 | SRR1039503 | 39 |
| Microspores | Col0 | SRR548296 | 40 |
| Vegetative cells | Col0 | SRR548294 | 40 |
| Sperm cells | Col0 | SRR548295 | 40 |
| Central cells | Col0 | SRR5014630 | 41 |
| Maturing embryos | Col0 | SRR017253 | 42 |
| Mature embryos | Col0 | SRR4051137 | * |
| Endosperms | Col0 | SRR017257 | 42 |
| Wild type | Col0 | SRR534177 | |
| *nrpb2-3* | Col0 | SRR534180 | |
| *nrpd1-4* | Col0 | SRR534181 | |
| *nrpe1-11* | Col0 | SRR534182 | |
| *rdr2-2* | Col0 | SRR534186 | |
| *rdr6-15* | Col0 | SRR534187 | |
| *dcl2-1, dcl3-1, dcl4-2* | Col0 | SRR534214 | |
| *ago4-5* | Col0 | SRR534197 | |
| *ago6-2* | Col0 | SRR534199 | 35 |
| *ago9-2* | Col0 | SRR534202 | |
| *drm1-2, drm2-2* | Col0 | SRR534222 | |
| *ddm1-2* | Col0 | SRR534215 | |
| *met1-3* | Col0 | SRR534239 | |
| *cmt2* | Col0 | SRR869314 | |
| *cmt3-11* | Col0 | SRR534209 | |
| *kyp* | Col0 | SRR534250 | |
| *mom1-2* | Col0 | SRR534178 | |
| *ibm1* | Col0 | SRR534234 | |

* : Bouyer, D. et al. DNA methylation dynamics during early plant life. Genome Biol 18, 179-190 (2017)

**Supplementary Table 3. Stastistical analysis of the chromatin states distribution**

| | S+C vs. A. tha | | | | S vs. A. tha | | | |
|---|---|---|---|---|---|---|---|---|
| | odds ratio [a] | *p*-value [b] | RF [c] | 95% CI | odds ratio [a] | *p*-value [b] | RF [c] | 95% CI |
| CS1 | 1.13 | 0.12 | 1.11 | 0.95 | ***1.46*** | ***0.36e$^{-3}$*** | ***1.36*** | ***1.22*** |
| CS2 | ***2.03*** | ***1.57e$^{-15}$*** | ***1.71*** | ***1.76*** | ***2.77*** | ***< 2.2e$^{-16}$*** | ***2.09*** | ***2.38*** |
| CS3 | 0.28 | 1.00 | 0.31 | 0.19 | 0.31 | 1.00 | 0.34 | 0.22 |
| CS4 | 0.89 | 0.88 | 0.91 | 0.74 | 1.15 | 0.11 | 1.11 | 0.95 |
| CS5 | 0.25 | 1.00 | 0.26 | 0.15 | 0.31 | 1.00 | 0.32 | 0.18 |
| CS6 | 0.85 | 0.89 | 0.87 | 0.67 | 1.03 | 0.42 | 1.03 | 0.81 |
| CS7 | 1.12 | 0.27 | 1.11 | 0.83 | 0.07 | 1.00 | 0.07 | 0.01 |
| CS8 | ***2.19*** | ***2.46e$^{-10}$*** | ***2.01*** | 1.80 | 0.23 | 1.00 | 0.25 | 0.12 |
| CS9 | 0.11 | 1.00 | 0.11 | 0.02 | 0.13 | 1.00 | 0.14 | 0.02 |

[a] A greater than 1 odds ratio (indicated in bolt and italic in the table together with corresponding *p*-value) implies a positive

[b] The null hypothesis that there's no difference between the means was rejected when the *p*-value was <0.005 (considered as

[c] A Representation Factor (RF) >1 indicates more overlap than expected between two populations.

**Supplementary Table 4 - Mutant genotypes used in this study**

| Mutant line | Type | Collection ID | Locus ID | Protein | Experiment |
|---|---|---|---|---|---|
| *nrpb2-3* | EMS | / | AT4G21710 | DNA-DIRECTED RNA POLYMERASE SUBUNIT 2 | Methylation profiles |
| *nrpd1-1* | T-DNA | SALK_583051 | AT1G63020 | DNA-DIRECTED RNA POLYMERASE SUBUNIT 1A | Northern blot |
| *nrpd1-4* | T-DNA | SALK_083051 | AT1G63020 | DNA-DIRECTED RNA POLYMERASE SUBUNIT 1A | Methylation profiles |
| *nrpe1-11* | T-DNA | SALK_029919 | AT2G40030 | DNA-DIRECTED RNA POLYMERASE SUBUNIT 1B | Northern blot / Methylation profiles |
| *rdr2-1* | T-DNA | SAIL_1277H08 | AT4G11130 | RNA-DEPENDENT RNA POLYMERASE 2 | Northern blot |
| *rdr2-2* | T-DNA | SALK_059661 | AT4G11130 | RNA-DEPENDENT RNA POLYMERASE 2 | Methylation profiles |
| *rdr6-15* | T-DNA | SAIL_617_H07 | AT3G49500 | RNA-DEPENDENT RNA POLYMERASE 6 | Methylation profiles |
| *dcl2-1* | T-DNA | SALK_064627 | AT3G03300 | DICER-LIKE PROTEIN 2 | Methylation profiles |
| *dcl3-1* | T-DNA | SALK_005512 | AT3G43920 | DICER-LIKE PROTEIN 3 | Northern blot / Methylation profiles |
| *dcl4-2* | T-DNA | GABI_160G05 | AT5G20320 | DICER-LIKE PROTEIN 4 | Methylation profiles |
| *ago4-1* | EMS | / | AT2G27040 | ARGONAUTE 4 | Northern blot |
| *ago4-5* | T-DNA | CS9927 | AT2G27040 | ARGONAUTE 4 | Methylation profiles |
| *ago6-2* | T-DNA | SALK_031553 | AT2G32940 | ARGONAUTE 6 | Methylation profiles |
| *ago9-1* | T-DNA | SALK_127358 | AT5G21150 | ARGONAUTE 9 | Northern blot |
| *ago9-2* | T-DNA | SALK_112059 | AT5G21150 | ARGONAUTE 9 | Methylation profiles |
| *drm1-2* | T-DNA | SALK_031705 | AT5G15380 | DNA METHYLTRANSFERASE 1 | Methylation profiles |
| *drm2-2* | T-DNA | SALK_150863 | AT5G14620 | DNA METHYLTRANSFERASE 2 | Methylation profiles |
| *drm2-10* | T-DNA | SALK_129477 | AT5G14620 | DNA METHYLTRANSFERASE 2 | Northern blot |
| *ddm1-1* | EMS | / | AT5G66750 | DEFICIENT IN DNA METHYLATION 1 | Crossing/Northern blot |
| *ddm1-2* | EMS | / | AT5G66750 | DEFICIENT IN DNA METHYLATION 1 | Northern blot / Methylation profiles |
| *met1-3* | T-DNA | CS16394 | AT5G49160 | METHYLTRANSFERASE 1 | Methylation profiles |
| *met1-7* | T-DNA | SALK_076522.32.35 | AT5G49160 | METHYLTRANSFERASE 1 | Northern blot |
| *cmt2* | T-DNA | WISCDSLOX7E02 | AT4G19020 | CHROMOMETHYLASE 2 | Methylation profiles |
| *cmt2-3* | T-DNA | SALK_012874 | AT4G19020 | CHROMOMETHYLASE 2 | Northern blot |
| *cmt3-11* | EMS | / | AT1G69770 | CHROMOMETHYLASE 3 | Northern blot / Methylation profiles |
| *kyp* | T-DNA | SALK_41474 | AT5G13960 | KRYPTONITE (SUVH4) | Methylation profiles |
| *kyp* | T-DNA | SALK_069326 | AT5G13960 | KRYPTONITE (SUVH4) | Northern blot |
| *mom1-2* | T-DNA | SAIL_610_G01 | AT1G08060 | MORPHEUS MOLECULE 1 | Northern blot / Methylation profiles |
| *ibm1* | T-DNA | SALK_006042 | AT3G07610 | INCREASE IN BONSAI METHYLATION 1 | Methylation profiles |

;

### Supplementary data 1

**Pairwise Wilcoxson tests related to the boxplots presented on Figures 3 and 4.**

**[1] "Pro"**

Wilcoxon rank sum test with continuity correction

data: Leaves_tDNAs_methylations.dt[context == "CG" & type == "Clustered"]$perc_methylation and Leaves_tDNAs_methylations.dt[context == "CG" & type == "Dispersed"]$perc_methylation
W = 121464, **p-value < 2.2e-16**
alternative hypothesis: true location shift is not equal to 0

Wilcoxon rank sum test with continuity correction

data: Leaves_tDNAs_methylations.dt[context == "CHG" & type == "Clustered"]$perc_methylation and Leaves_tDNAs_methylations.dt[context == "CHG" & type == "Dispersed"]$perc_methylation
W = 10720, **p-value = 0.6026**
alternative hypothesis: true location shift is not equal to 0

Wilcoxon rank sum test with continuity correction

data: Leaves_tDNAs_methylations.dt[context == "CHH" & type == "Clustered"]$perc_methylation and Leaves_tDNAs_methylations.dt[context == "CHH" & type == "Dispersed"]$perc_methylation
W = 878277, **p-value = 0.3653**
alternative hypothesis: true location shift is not equal to 0

**[1] "Ser"**

Wilcoxon rank sum test with continuity correction

data: Leaves_tDNAs_methylations.dt[context == "CG" & type == "Clustered"]$perc_methylation and Leaves_tDNAs_methylations.dt[context == "CG" & type == "Dispersed"]$perc_methylation
W = 254752, **p-value < 2.2e-16**
alternative hypothesis: true location shift is not equal to 0

Wilcoxon rank sum test with continuity correction

data:  Leaves_tDNAs_methylations.dt[context == "CHG" & type == "Clustered"]$perc_methylation and Leaves_tDNAs_methylations.dt[context == "CHG" & type == "Dispersed"]$perc_methylation
W = 70038, **p-value < 2.2e-16**
alternative hypothesis: true location shift is not equal to 0

Wilcoxon rank sum test with continuity correction

data:  Leaves_tDNAs_methylations.dt[context == "CHH" & type == "Clustered"]$perc_methylation and Leaves_tDNAs_methylations.dt[context == "CHH" & type == "Dispersed"]$perc_methylation
W = 664228, **p-value = 7.883e-16**
alternative hypothesis: true location shift is not equal to 0

**[1] "Tyr"**

Wilcoxon rank sum test with continuity correction

data:  Leaves_tDNAs_methylations.dt[context == "CG" & type == "Clustered"]$perc_methylation and Leaves_tDNAs_methylations.dt[context == "CG" & type == "Dispersed"]$perc_methylation
W = 138240, **p-value < 2.2e-16**
alternative hypothesis: true location shift is not equal to 0

Wilcoxon rank sum test with continuity correction

data:  Leaves_tDNAs_methylations.dt[context == "CHG" & type == "Clustered"]$perc_methylation and Leaves_tDNAs_methylations.dt[context == "CHG" & type == "Dispersed"]$perc_methylation
W = 185790**, p-value < 2.2e-16**
alternative hypothesis: true location shift is not equal to 0

Wilcoxon rank sum test with continuity correction

data:  Leaves_tDNAs_methylations.dt[context == "CHH" & type == "Clustered"]$perc_methylation and Leaves_tDNAs_methylations.dt[context == "CHH" & type == "Dispersed"]$perc_methylation
W = 679052, **p-value < 2.2e-16**
alternative hypothesis: true location shift is not equal to 0

**[1] "Cys"**

Wilcoxon rank sum test with continuity correction

data:  Leaves_tDNAs_methylations.dt[context == "CG" & type == "Clustered"]$perc_methylation and Leaves_tDNAs_methylations.dt[context == "CG" & type == "Dispersed"]$perc_methylation
W = 1399, **p-value = 3.179e-12**
alternative hypothesis: true location shift is not equal to 0

Wilcoxon rank sum test with continuity correction

data:  Leaves_tDNAs_methylations.dt[context == "CHG" & type == "Clustered"]$perc_methylation and Leaves_tDNAs_methylations.dt[context == "CHG" & type == "Dispersed"]$perc_methylation
W = 4762.5, **p-value < 2.2e-16**
alternative hypothesis: true location shift is not equal to 0

Wilcoxon rank sum test with continuity correction

data:  Leaves_tDNAs_methylations.dt[context == "CHH" & type == "Clustered"]$perc_methylation and Leaves_tDNAs_methylations.dt[context == "CHH" & type == "Dispersed"]$perc_methylation
W = 39989, **p-value < 2.2e-16**
alternative hypothesis: true location shift is not equal to 0

**Supplementary data 2 :** p-value of pairwise Wilcoxson test

**Pro CG clustered**

| | **Col0** | ***drm12*** | ***ddm1*** | ***met1*** | ***cmt2*** | ***cmt3*** | ***kyp*** | ***mom1*** |
|---|---|---|---|---|---|---|---|---|
| ***drm12*** | 1 | | | | | | | |
| ***ddm1*** | 0.724 | 1 | | | | | | |
| ***met1*** | 0 | 0 | 0 | | | | | |
| ***cmt2*** | 1 | 1 | 0.6446 | 0 | | | | |
| ***cmt3*** | 1 | 1 | 1 | 0 | 1 | | | |
| ***kyp*** | 1 | 1 | 1 | 0 | 1 | 1 | | |
| ***mom1*** | 1 | 1 | 1 | 0 | 1 | 1 | 1 | |
| ***ibm1*** | 1 | 1 | 0.02493 | 0 | 1 | 1 | 1 | 1 |

**Pro CHG clustered**

| | **Col0** | ***drm12*** | ***ddm1*** | ***met1*** | ***cmt2*** | ***cmt3*** | ***kyp*** | ***mom1*** |
|---|---|---|---|---|---|---|---|---|
| ***drm12*** | 1 | | | | | | | |
| ***ddm1*** | 1 | 0.00531 | | | | | | |
| ***met1*** | 1 | 1 | 0.45803 | | | | | |
| ***cmt2*** | 1 | 1 | 1 | 1 | | | | |
| ***cmt3*** | 1 | 1 | 0.08958 | 1 | 1 | | | |
| ***kyp*** | 1 | 0.57919 | 1 | 1 | 1 | 1 | | |
| ***mom1*** | 1 | 0.39589 | 1 | 1 | 1 | 1 | 1 | |
| ***ibm1*** | 0 | 0 | 0 | 0 | 0 | 0 | 0 | 0 |

**Pro CHH clustered**

| | **Col0** | ***drm12*** | ***ddm1*** | ***met1*** | ***cmt2*** | ***cmt3*** | ***kyp*** | ***mom1*** |
|---|---|---|---|---|---|---|---|---|
| ***drm12*** | 0.00006 | | | | | | | |
| ***ddm1*** | 1 | 0.0364 | | | | | | |
| ***met1*** | 0.01912 | 0.24146 | 1 | | | | | |
| ***cmt2*** | 1 | 0.04282 | 1 | 1 | | | | |
| ***cmt3*** | 0.01933 | 1 | 1 | 1 | 1 | | | |
| ***kyp*** | 0.89595 | 0.12026 | 1 | 1 | 1 | 1 | | |
| ***mom1*** | 1 | 0.00322 | 1 | 1 | 1 | 1 | 1 | |
| ***ibm1*** | 0.00019 | 0 | 0.00003 | 0 | 0 | 0 | 0 | 0 |

**Supplementary data 2:** p-value of pairwise Wilcoxson test

#### Ser CG clustered

| | Col0 | *drm12* | *ddm1* | *met1* | *cmt2* | *cmt3* | *kyp* | *mom1* |
|---|---|---|---|---|---|---|---|---|
| ***drm12*** | 0.61554 | | | | | | | |
| ***ddm1*** | 0 | 0 | | | | | | |
| ***met1*** | 0 | 0 | 0 | | | | | |
| ***cmt2*** | 1 | 0.16538 | 0 | 0 | | | | |
| ***cmt3*** | 1 | 1 | 0 | 0 | 1 | | | |
| ***kyp*** | 1 | 1 | 0 | 0 | 1 | 1 | | |
| ***mom1*** | 1 | 1 | 0 | 0 | 1 | 1 | 1 | |
| ***ibm1*** | 1 | 1 | 0 | 0 | 1 | 1 | 1 | 1 |

#### Ser CHG clustered

| | Col0 | *drm12* | *ddm1* | *met1* | *cmt2* | *cmt3* | *kyp* | *mom1* |
|---|---|---|---|---|---|---|---|---|
| ***drm12*** | 1 | | | | | | | |
| ***ddm1*** | 1 | 1 | | | | | | |
| ***met1*** | 0 | 0.00023 | 0 | | | | | |
| ***cmt2*** | 1 | 1 | 1 | 0.00039 | | | | |
| ***cmt3*** | 0 | 0.00001 | 0 | 1 | 0.00002 | | | |
| ***kyp*** | 0 | 0.00003 | 0 | 1 | 0.00003 | 1 | | |
| ***mom1*** | 1 | 1 | 1 | 0.00001 | 1 | 0 | 0 | |
| ***ibm1*** | 1 | 1 | 1 | 0.00005 | 1 | 0 | 0 | 1 |

#### Ser CHH clustered

| | Col0 | *drm12* | *ddm1* | *met1* | *cmt2* | *cmt3* | *kyp* | *mom1* |
|---|---|---|---|---|---|---|---|---|
| ***drm12*** | 0.06932 | | | | | | | |
| ***ddm1*** | 0 | 0 | | | | | | |
| ***met1*** | 0.02273 | 1 | 0 | | | | | |
| ***cmt2*** | 0.04065 | 1 | 0 | 1 | | | | |
| ***cmt3*** | 0.53932 | 1 | 0 | 1 | 1 | | | |
| ***kyp*** | 0.00005 | 1 | 0 | 1 | 1 | 1 | | |
| ***mom1*** | 1 | 1 | 0 | 1 | 1 | 1 | 0.03562 | |
| ***ibm1*** | 1 | 1 | 0 | 1 | 0.95621 | 1 | 0.03806 | 1 |

**Supplementary data 2 :** p-value of pairwise Wilcoxson test

**Tyr CG clustered**

| | **Col0** | ***drm12*** | ***ddm1*** | ***met1*** | ***cmt2*** | ***cmt3*** | ***kyp*** | ***mom1*** |
|---|---|---|---|---|---|---|---|---|
| ***drm12*** | 0.00242 | | | | | | | |
| ***ddm1*** | 0 | 0 | | | | | | |
| ***met1*** | 0 | 0 | 0 | | | | | |
| ***cmt2*** | 0.09534 | 1 | 0 | 0 | | | | |
| ***cmt3*** | 1 | 0.04701 | 0 | 0 | 1 | | | |
| ***kyp*** | 0.57129 | 0.79006 | 0 | 0 | 1 | 1 | | |
| ***mom1*** | 1 | 0.04237 | 0 | 0 | 1 | 1 | 1 | |
| ***ibm1*** | 1 | 0.00001 | 0 | 0 | 0.00121 | 0.66269 | 0.01029 | 0.37379 |

**Tyr CHH clustered**

| | **Col0** | ***drm12*** | ***ddm1*** | ***met1*** | ***cmt2*** | ***cmt3*** | ***kyp*** | ***mom1*** |
|---|---|---|---|---|---|---|---|---|
| ***drm12*** | 1 | | | | | | | |
| ***ddm1*** | 0 | 0 | | | | | | |
| ***met1*** | 0 | 0 | 0 | | | | | |
| ***cmt2*** | 0.00048 | 0.00015 | 0 | 0 | | | | |
| ***cmt3*** | 0 | 0 | 1 | 0 | 0 | | | |
| ***kyp*** | 0 | 0 | 1 | 0 | 0 | 1 | | |
| ***mom1*** | 1 | 1 | 0 | 0 | 0.01549 | 0 | 0 | |
| ***ibm1*** | 0 | 0 | 0 | 0 | 1 | 0 | 0 | 0.0002 |

**Tyr CHH clustered**

| | **Col0** | ***drm12*** | ***ddm1*** | ***met1*** | ***cmt2*** | ***cmt3*** | ***kyp*** | ***mom1*** |
|---|---|---|---|---|---|---|---|---|
| ***drm12*** | 0.22449 | | | | | | | |
| ***ddm1*** | 0 | 0 | | | | | | |
| ***met1*** | 0 | 0 | 0 | | | | | |
| ***cmt2*** | 0 | 0 | 1 | 0 | | | | |
| ***cmt3*** | 0 | 0 | 0.04539 | 0 | 0.01013 | | | |
| ***kyp*** | 0 | 0 | 1 | 0 | 1 | 0.00004 | | |
| ***mom1*** | 0.00002 | 1 | 0 | 0 | 0 | 0 | 0 | |
| ***ibm1*** | 0.22449 | 1 | 0 | 0 | 0 | 0 | 0 | 0.17813 |

**Supplementary data 2 :** p-value of pairwise Wilcoxson test

**Cys CG clustered**

| | Col0 | *drm12* | *ddm1* | *met1* | *cmt2* | *cmt3* | *kyp* | *mom1* |
|---|---|---|---|---|---|---|---|---|
| ***drm12*** | 1 | | | | | | | |
| ***ddm1*** | 1 | 1 | | | | | | |
| ***met1*** | 1 | 1 | 1 | | | | | |
| ***cmt2*** | 1 | 1 | 1 | 1 | | | | |
| ***cmt3*** | 1 | 1 | 1 | 1 | 1 | | | |
| ***kyp*** | 1 | 1 | 1 | 1 | 1 | 1 | | |
| ***mom1*** | 1 | 1 | 1 | 1 | 1 | 1 | 1 | |
| ***ibm1*** | 1 | 1 | 1 | 1 | 1 | 1 | 1 | 1 |

**Cys CHG clustered**

| | Col0 | *drm12* | *ddm1* | *met1* | *cmt2* | *cmt3* | *kyp* | *mom1* |
|---|---|---|---|---|---|---|---|---|
| ***drm12*** | 1 | | | | | | | |
| ***ddm1*** | 1 | 1 | | | | | | |
| ***met1*** | 1 | 1 | 1 | | | | | |
| ***cmt2*** | 1 | 1 | 1 | 1 | | | | |
| ***cmt3*** | 1 | 1 | 1 | 1 | 1 | | | |
| ***kyp*** | 1 | 1 | 1 | 1 | 1 | 1 | | |
| ***mom1*** | 1 | 1 | 1 | 1 | 1 | 1 | 1 | |
| ***ibm1*** | 1 | 1 | 1 | 1 | 1 | 1 | 1 | 1 |

**Cys CHH clustered**

| | Col0 | *drm12* | *ddm1* | *met1* | *cmt2* | *cmt3* | *kyp* | *mom1* |
|---|---|---|---|---|---|---|---|---|
| ***drm12*** | 1 | | | | | | | |
| ***ddm1*** | 1 | 1 | | | | | | |
| ***met1*** | 1 | 1 | 1 | | | | | |
| ***cmt2*** | 1 | 1 | 1 | 1 | | | | |
| ***cmt3*** | 1 | 1 | 1 | 1 | 1 | | | |
| ***kyp*** | 1 | 1 | 1 | 1 | 1 | 1 | | |
| ***mom1*** | 1 | 1 | 1 | 1 | 1 | 1 | 1 | |
| ***ibm1*** | 1 | 1 | 1 | 1 | 1 | 1 | 1 | 1 |

**Supplementary data 2 :** p-value of pairwise Wilcoxson test

**Pro CG dispersed**

| | Col0 | drm12 | ddm1 | met1 | cmt2 | cmt3 | kyp | mom1 |
|---|---|---|---|---|---|---|---|---|
| ***drm12*** | 1 | | | | | | | |
| ***ddm1*** | 1 | 1 | | | | | | |
| ***met1*** | 0,00002 | 0,00001 | 0,00012 | | | | | |
| ***cmt2*** | 1 | 1 | 1 | 0 | | | | |
| ***cmt3*** | 1 | 1 | 1 | 0 | 1 | | | |
| ***kyp*** | 1 | 1 | 1 | 0 | 1 | 1 | | |
| ***mom1*** | 1 | 1 | 1 | 0 | 1 | 1 | 1 | |
| ***ibm1*** | 1 | 1 | 1 | 0,00008 | 1 | 1 | 1 | 1 |

**Pro CHG dispersed**

| | **Col0** | ***drm12*** | ***ddm1*** | ***met1*** | ***cmt2*** | ***cmt3*** | ***kyp*** | ***mom1*** |
|---|---|---|---|---|---|---|---|---|
| ***drm12*** | 1 | | | | | | | |
| ***ddm1*** | 1 | 1 | | | | | | |
| ***met1*** | 1 | 1 | 1 | | | | | |
| ***cmt2*** | 1 | 1 | 1 | 1 | | | | |
| ***cmt3*** | 1 | 1 | 1 | 1 | 1 | | | |
| ***kyp*** | 1 | 1 | 1 | 1 | 1 | 1 | | |
| ***mom1*** | 1 | 1 | 1 | 1 | 1 | 1 | 1 | |
| ***ibm1*** | 0.00056 | 0.00192 | 0.05741 | 0.00283 | 0.20522 | 0.00053 | 0.00034 | 0.00566 |

**Pro CHH dispersed**

| | **Col0** | ***drm12*** | ***ddm1*** | ***met1*** | ***cmt2*** | ***cmt3*** | ***kyp*** | ***mom1*** |
|---|---|---|---|---|---|---|---|---|
| ***drm12*** | 1 | | | | | | | |
| ***ddm1*** | 1 | 1 | | | | | | |
| ***met1*** | 1 | 1 | 1 | | | | | |
| ***cmt2*** | 1 | 0.09376 | 1 | 0.31755 | | | | |
| ***cmt3*** | 1 | 1 | 1 | 1 | 1 | | | |
| ***kyp*** | 1 | 1 | 1 | 1 | 0.22891 | 1 | | |
| ***mom1*** | 1 | 1 | 1 | 1 | 1 | 1 | 1 | |
| ***ibm1*** | 1 | 0.02096 | 1 | 0.05222 | 1 | 1 | 0.04238 | 1 |

**Supplementary data 2 :** p-value of pairwise Wilcoxson test

**Ser CG dispersed**

| | Col0 | *drm12* | *ddm1* | *met1* | *cmt2* | *cmt3* | *kyp* | *mom1* |
|---|---|---|---|---|---|---|---|---|
| ***drm12*** | 1 | | | | | | | |
| ***ddm1*** | 0.87208 | 1 | | | | | | |
| ***met1*** | 0.11795 | 1 | 1 | | | | | |
| ***cmt2*** | 1 | 1 | 1 | 0.08189 | | | | |
| ***cmt3*** | 1 | 1 | 1 | 0.28781 | 1 | | | |
| ***kyp*** | 1 | 1 | 0.51015 | 0.00481 | 1 | 1 | | |
| ***mom1*** | 1 | 1 | 1 | 0.69927 | 1 | 1 | 1 | |
| ***ibm1*** | 1 | 1 | 0.04524 | 0.00045 | 1 | 1 | 1 | 1 |

**Ser CHG dispersed**

| | Col0 | *drm12* | *ddm1* | *met1* | *cmt2* | *cmt3* | *kyp* | *mom1* |
|---|---|---|---|---|---|---|---|---|
| ***drm12*** | 1 | | | | | | | |
| ***ddm1*** | 1 | 1 | | | | | | |
| ***met1*** | 0.36406 | 1 | 1 | | | | | |
| ***cmt2*** | 1 | 1 | 0.50195 | 0.06146 | | | | |
| ***cmt3*** | 0.91208 | 1 | 1 | 1 | 0.82183 | | | |
| ***kyp*** | 1 | 1 | 1 | 1 | 1 | 1 | | |
| ***mom1*** | 1 | 1 | 1 | 1 | 1 | 1 | 1 | |
| ***ibm1*** | 1 | 1 | 0.0111 | 0.00093 | 1 | 0.0063 | 0.02772 | 1 |

**Ser CHH dispersed**

| | Col0 | *drm12* | *ddm1* | *met1* | *cmt2* | *cmt3* | *kyp* | *mom1* |
|---|---|---|---|---|---|---|---|---|
| ***drm12*** | 0.8407 | | | | | | | |
| ***ddm1*** | 1 | 1 | | | | | | |
| ***met1*** | 1 | 1 | 1 | | | | | |
| ***cmt2*** | 1 | 0.13693 | 1 | 1 | | | | |
| ***cmt3*** | 1 | 1 | 1 | 1 | 1 | | | |
| ***kyp*** | 1 | 1 | 1 | 1 | 1 | 1 | | |
| ***mom1*** | 1 | 1 | 1 | 1 | 1 | 1 | 1 | |
| ***ibm1*** | 1 | 0.01652 | 1 | 0.22843 | 1 | 0.11613 | 0.36626 | 1 |

**Supplementary data 2 :** p-value of pairwise Wilcoxson test

**Tyr CG dispersed**

| | Col0 | *drm12* | *ddm1* | *met1* | *cmt2* | *cmt3* | *kyp* | *mom1* |
|---|---|---|---|---|---|---|---|---|
| ***drm12*** | 1 | | | | | | | |
| ***ddm1*** | 1 | 1 | | | | | | |
| ***met1*** | 1 | 1 | 1 | | | | | |
| ***cmt2*** | 1 | 1 | 1 | 1 | | | | |
| ***cmt3*** | 1 | 1 | 1 | 1 | 1 | | | |
| ***kyp*** | 1 | 1 | 1 | 1 | 1 | 1 | | |
| ***mom1*** | 1 | 1 | 1 | 1 | 1 | 1 | 1 | |
| ***ibm1*** | 1 | 1 | 1 | 1 | 1 | 1 | 1 | 1 |

***Tyr CHG dispersed***

| | Col0 | *drm12* | *ddm1* | *met1* | *cmt2* | *cmt3* | *kyp* | *mom1* |
|---|---|---|---|---|---|---|---|---|
| ***drm12*** | 1 | | | | | | | |
| ***ddm1*** | 1 | 1 | | | | | | |
| ***met1*** | 1 | 1 | 1 | | | | | |
| ***cmt2*** | 1 | 1 | 1 | 1 | | | | |
| ***cmt3*** | 1 | 1 | 1 | 1 | 1 | | | |
| ***kyp*** | 1 | 1 | 1 | 1 | 1 | 1 | | |
| ***mom1*** | 1 | 1 | 1 | 1 | 1 | 1 | 1 | |
| ***ibm1*** | 1 | 1 | 1 | 1 | 1 | 0.90572 | 1 | 1 |

***Tyr CHH dispersed***

| | Col0 | *drm12* | *ddm1* | *met1* | *cmt2* | *cmt3* | *kyp* | *mom1* |
|---|---|---|---|---|---|---|---|---|
| ***drm12*** | 1 | | | | | | | |
| ***ddm1*** | 1 | 1 | | | | | | |
| ***met1*** | 1 | 1 | 1 | | | | | |
| ***cmt2*** | 1 | 1 | 1 | 1 | | | | |
| ***cmt3*** | 1 | 1 | 1 | 1 | 1 | | | |
| ***kyp*** | 1 | 1 | 1 | 1 | 1 | 1 | | |
| ***mom1*** | 1 | 1 | 1 | 1 | 1 | 1 | 1 | |
| ***ibm1*** | 1 | 1 | 1 | 1 | 1 | 1 | 1 | 1 |

**Supplementary data 2 :** p-value of pairwise Wilcoxson test

**Cys CG dispersed**

| | Col0 | *drm12* | *ddm1* | *met1* | *cmt2* | *cmt3* | *kyp* | *mom1* |
|---|---|---|---|---|---|---|---|---|
| ***drm12*** | 1 | | | | | | | |
| ***ddm1*** | 1 | 1 | | | | | | |
| ***met1*** | 0.30961 | 1 | 1 | | | | | |
| ***cmt2*** | 1 | 1 | 1 | 1 | | | | |
| ***cmt3*** | 1 | 1 | 1 | 1 | 1 | | | |
| ***kyp*** | 1 | 1 | 1 | 1 | 1 | 1 | | |
| ***mom1*** | 1 | 1 | 1 | 1 | 1 | 1 | 1 | |
| ***ibm1*** | 1 | 1 | 1 | 0.30961 | 1 | 1 | 1 | 1 |

**Cys CHG dispersed**

| | Col0 | *drm12* | *ddm1* | *met1* | *cmt2* | *cmt3* | *kyp* | *mom1* |
|---|---|---|---|---|---|---|---|---|
| ***drm12*** | 1 | | | | | | | |
| ***ddm1*** | 1 | 1 | | | | | | |
| ***met1*** | 1 | 1 | 1 | | | | | |
| ***cmt2*** | 1 | 1 | 1 | 1 | | | | |
| ***cmt3*** | 1 | 1 | 1 | 1 | 1 | | | |
| ***kyp*** | 1 | 1 | 1 | 1 | 1 | 1 | | |
| ***mom1*** | 1 | 1 | 1 | 1 | 1 | 1 | 1 | |
| ***ibm1*** | 1 | 1 | 1 | 1 | 1 | 1 | 1 | 1 |

**Cys CHH dispersed**

| | Col0 | *drm12* | *ddm1* | *met1* | *cmt2* | *cmt3* | *kyp* | *mom1* |
|---|---|---|---|---|---|---|---|---|
| ***drm12*** | 1 | | | | | | | |
| ***ddm1*** | 1 | 1 | | | | | | |
| ***met1*** | 1 | 1 | 1 | | | | | |
| ***cmt2*** | 1 | 1 | 1 | 1 | | | | |
| ***cmt3*** | 1 | 1 | 1 | 1 | 1 | | | |
| ***kyp*** | 1 | 1 | 1 | 1 | 1 | 1 | | |
| ***mom1*** | 1 | 1 | 1 | 1 | 1 | 1 | 1 | |
| ***ibm1*** | 1 | 1 | 1 | 1 | 1 | 1 | 1 | 1 |

**Supplementary data 2 :** p-value of pairwise Wilcoxson test

**Ala CG dispersed**

| | **Col0** | ***drm12*** | ***ddm1*** | ***met1*** | ***cmt2*** | ***cmt3*** | ***kyp*** | ***mom1*** |
|---|---|---|---|---|---|---|---|---|
| ***drm12*** | 0.00379 | | | | | | | |
| ***ddm1*** | 1 | 0 | | | | | | |
| ***met1*** | 0.03362 | 1 | 0.00001 | | | | | |
| ***cmt2*** | 1 | 0.00038 | 1 | 0.00207 | | | | |
| ***cmt3*** | 1 | 0.00103 | 1 | 0.01131 | 1 | | | |
| ***kyp*** | 1 | 0.00007 | 1 | 0.00023 | 1 | 1 | | |
| ***mom1*** | 1 | 0.01556 | 1 | 0.26251 | 1 | 1 | 1 | |
| ***ibm1*** | 1 | 0.00101 | 1 | 0.00463 | 1 | 1 | 1 | 1 |

**Ala CHG dispersed**

| | **Col0** | ***drm12*** | ***ddm1*** | ***met1*** | ***cmt2*** | ***cmt3*** | ***kyp*** | ***mom1*** |
|---|---|---|---|---|---|---|---|---|
| ***drm12*** | 0.124 | | | | | | | |
| ***ddm1*** | 1 | 0.03072 | | | | | | |
| ***met1*** | 1 | 1 | 1 | | | | | |
| ***cmt2*** | 1 | 1 | 1 | 1 | | | | |
| ***cmt3*** | 1 | 0.2257 | 1 | 1 | 1 | | | |
| ***kyp*** | 1 | 0.08647 | 1 | 1 | 1 | 1 | | |
| ***mom1*** | 1 | 0.90774 | 1 | 1 | 1 | 1 | 1 | |
| ***ibm1*** | 1 | 0.20975 | 1 | 1 | 1 | 1 | 1 | 1 |

**Ala CHH dispersed**

| | **Col0** | ***drm12*** | ***ddm1*** | ***met1*** | ***cmt2*** | ***cmt3*** | ***kyp*** | ***mom1*** |
|---|---|---|---|---|---|---|---|---|
| ***drm12*** | 0.00452 | | | | | | | |
| ***ddm1*** | 1 | 0.00004 | | | | | | |
| ***met1*** | 0.8216 | 0.87515 | 0.04512 | | | | | |
| ***cmt2*** | 1 | 0.03151 | 1 | 1 | | | | |
| ***cmt3*** | 1 | 0.06578 | 1 | 1 | 1 | | | |
| ***kyp*** | 1 | 0.00851 | 1 | 1 | 1 | 1 | | |
| ***mom1*** | 1 | 1 | 1 | 1 | 1 | 1 | 1 | |
| ***ibm1*** | 1 | 0.00744 | 1 | 1 | 1 | 1 | 1 | 1 |
